# Genetic dissection of a *Leishmania* flagellar proteome demonstrates requirement for directional motility in sand fly infections

**DOI:** 10.1101/476994

**Authors:** Tom Beneke, François Demay, Edward Hookway, Nicole Ashman, Heather Jeffery, James Smith, Jessica Valli, Tomas Becvar, Jitka Myskova, Tereza Lestinova, Shahaan Shafiq, Jovana Sadlova, Petr Volf, Richard Wheeler, Eva Gluenz

## Abstract

The protozoan parasite *Leishmania* possesses a single flagellum, which is remodelled during the parasite’s life cycle from a long motile flagellum in promastigote forms in the sand fly to a short immotile flagellum in amastigotes residing in mammalian phagocytes. This study examined the protein composition and *in vivo* function of the promastigote flagellum. Protein mass spectrometry and label free protein enrichment testing of isolated flagella and deflagellated cell bodies defined a flagellar proteome for *L. mexicana* promastigote forms (available via ProteomeXchange with identifier PXD011057). This information was used to generate a CRISPR-Cas9 knockout library of 100 mutants to screen for flagellar defects. This first large-scale knockout screen in a *Leishmania* sp. identified 56 mutants with altered swimming speed (52 reduced and 4 increased) and defined distinct mutant categories (faster swimmers, slower swimmers, slow uncoordinated swimmers and paralysed cells, including aflagellate promastigotes and cells with curled flagella and disruptions of the paraflagellar rod). Each mutant was tagged with a unique 17-nt barcode, providing a simple barcode sequencing (bar-seq) method for measuring the relative fitness of *L. mexicana* mutants *in vivo*. In mixed infections of the permissive sand fly vector *Lutzomyia longipalpis*, paralysed promastigotes and uncoordinated swimmers were severely diminished in the fly after defecation of the bloodmeal. Subsequent examination of flies infected with a single mutant lacking the central pair protein PF16 showed that these paralysed promastigotes did not reach anterior regions of the fly alimentary tract. These data show that *L. mexicana* need directional motility for successful colonisation of sand flies.

**Author Summary:** *Leishmania* are protozoan parasites, transmitted between mammals by the bite of phlebotomine sand flies. Promastigote forms in the sand fly have a long flagellum, which is motile and used for anchoring the parasites to prevent clearance with the digested blood meal remnants. To dissect flagellar functions and their importance in life cycle progression, we generated here a comprehensive list of >300 flagellar proteins and produced a CRISPR-Cas9 gene knockout library of 100 mutant *Leishmania*. We studied their behaviour *in vitro* before examining their fate in the sand fly *Lutzomyia longipalpis*. Measuring mutant swimming speeds showed that about half behaved differently compared to the wild type: a few swam faster, many slower and some were completely paralysed. We also found a group of uncoordinated swimmers. To test whether flagellar motility is required for parasite migration from the fly midgut to the foregut from where they reach the next host, we infected sand flies with a mixed mutant population. Each mutant carried a unique tag and tracking these tags up to nine days after infection showed that paralysed and uncoordinated *Leishmania* were rapidly lost from flies. These data indicate that directional swimming is important for successful colonisation of sand flies.

## Introduction

Eukaryotic flagella / cilia are complex multifunctional organelles conserved from protists to humans [1]. Protists use flagella for swimming, feeding, cell-to-cell communication, adherence to substrates and morphogenesis [2]. Single-celled organisms, most commonly among them the green algae *Chlamydomonas reinhardtii,* have served as important model organisms to study molecular mechanisms of ciliogenesis and ciliary function [3], spurred on by the recognition that ciliary defects cause human genetic disorders collectively termed “ciliopathies” [4]. The eukaryotic flagellum is a complex, highly structured organelle and dissection of the molecular mechanisms underpinning its diverse functions requires detailed knowledge of its component parts. Proteomic studies of isolated flagella or axonemes from diverse species typically identified at least 300 distinct proteins [5-9] and phylogenetic profiling identified a set of 274 evolutionarily conserved ciliary genes [10]. All of these datasets comprise many “hypothetical” proteins still awaiting functional characterisation in addition to well-characterised core components of the microtubule axoneme, associated motor proteins and regulatory complexes.

Insights into conserved ciliary biology have helped elucidation of flagellar function in eukaryotic microbes, with a particular focus on human pathogens [11,12]. Among these, flagella have been most extensively studied in the causative agent of African trypanosomiasis, *Trypanosoma brucei* [13], which uses flagellar motility for locomotion and immune evasion [14] and exhibits close spatio-temporal coordination between flagellum assembly and cell morphogenesis during division [15]. The *T. brucei* bloodstream form is particularly sensitive to the loss of flagellar function [6,16], highlighting a potential Achilles’ heel that might be exploitable for new anti-parasitic treatments.

The *Leishmania* flagellum is also a multi-functional organelle, which undergoes striking structural changes during the parasite’s life cycle [17-19]. Amastigote forms proliferating in mammalian macrophages possess a short sensory-type 9+0 microtubule axoneme, which is remodelled to a canonical long motile 9+2 axoneme during differentiation to promastigote forms, which live in blood-feeding phlebotomine sand flies (Diptera: Psychodidae). In the fly, nectomonad promastigote forms attach via their flagella to the microvilli of the posterior midgut [20] to protect the parasites from being cleared during defecation of remnants of the blood meal. In the oesophageal valve, broad haptomonad forms attach to the cuticular lining via their flagellar tips, forming hemidesmosomes [20]. These life cycle descriptions [21] imply that periods of attachment must be followed by migration to more anterior regions of the alimentary tract and the propulsive function of the *Leishmania* flagellum is presumed to drive this forward migration but this has not been directly tested.

To enable a detailed genetic dissection of flagellar functions and mechanisms in *Leishmania*, we defined here a flagellar proteome for motile *L. mexicana* promastigotes. We used new CRISPR-Cas9 genome editing methods [22] to generate a *Leishmania* knockout library of 100 mutants, over half of which showed altered swimming speed. We also developed a barcode sequencing (bar-seq) protocol to test the fitness of mutants in the permissive sand fly vector *Lutzomyia longipalpis*. This study identified new genes required for flagellar motility and shows that whilst culture-form promastigotes tolerated loss of the flagellum, paralysed mutants and uncoordinated swimmers failed to colonise sand flies indicating that directional flagellar motility is required for completion of the parasite’s life cycle.

## Results

### Defining the promastigote flagellar proteome

To enable a systematic genetic dissection of flagellar functions we sought to isolate *L. mexicana* promastigote flagella comprising the axoneme, extra-axonemal structures and the surrounding membrane for subsequent analysis by protein mass spectrometry (MS). Mechanical shearing in the presence of 75 mM Ca^2+^ successfully separated cells into flagella (F) and deflagellated cell bodies (CB) (Figure 1A, B). Subsequent centrifugation on sucrose gradients allowed isolation of F and CB fractions with little cross-contamination: the CB fraction contained only 2.03% (±0.69%) isolated flagella and the F fractions contained 0.56% (±0.15%) deflagellated cell bodies (S1 Figure). Isolated flagella still retained their membrane: First, examination of F fractions by transmission electron microscopy (TEM) confirmed that most axonemes were bounded by a membrane (S2 Figure) and second, tracking an abundant promastigote flagellar membrane protein, the small myristoylated protein 1 (SMP-1, [23]) tagged with enhanced green fluorescent protein (eGFP) showed that it remained associated with isolated flagella (Figure 1). Analysis of the SMP-1∷eGFP signal also facilitated flagellar length measurements in whole cells, F and CB fractions, which showed that flagella were separated from the cell body near the exit point from the flagellar pocket. The average break point was 2.7 μm distal to the base of the flagellum. The length of the isolated flagella was similar to those on intact cells, indicating that isolated flagella remained in one piece, with little fragmentation (S3 Figure). Two independently prepared sets of F and CB fractions were separated into detergent soluble (s) and insoluble fractions (i), yielding four fractions, F_S_, F_I_, CB_S_ and CB_I_ (Figure 1C). All four fractions for both replicates were analysed by liquid chromatography tandem mass spectrometry (MS), which detected a total of 2711 distinct proteins (Figure 1D). Enrichment of detected proteins between biological replicates correlated well (Pearson’s r > 0.72, Spearman’s *r*_*s*_ > 0.83, S4 Figure). To discover proteins enriched in each of the four fractions, we used a label-free normalized spectral index quantitation method (SINQ, [24]; S1, S2 and S3 Table) to generate a SINQ enrichment plot (Figure 2A). The promastigote flagellar proteome, defined as proteins enriched in F vs. CB fractions consisted of 701 unique proteins detected in at least one MS run; 352 of these were enriched in F vs. CB fractions in both MS runs.

**Figure 1.**
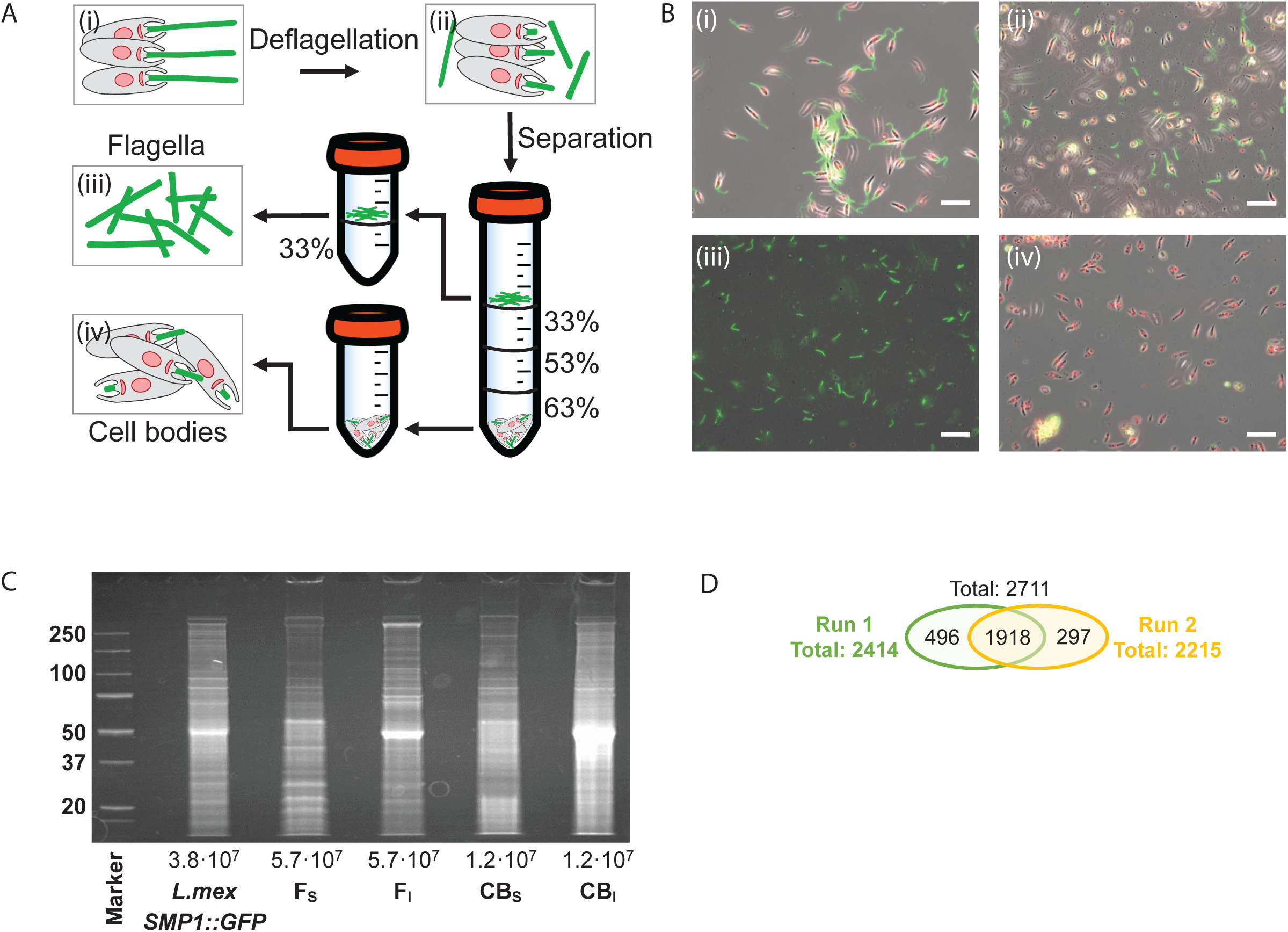
Isolation of flagella and deflagellated cell bodies. **(A)** Overview of the deflagellation and differential centrifugation protocol. Percentage sucrose concentration (w/v) is indicated. (**B)** Micrographs show merged phase and fluorescence channel (SMP-1∷GFP, green; Hoechst DNA stain, red) for each isolation stage (i-iv) depicted in (A). (**i)** *L. mexicana* SMP1∷GFP cells before deflagellation, (**ii)** cells after deflagellation, (**iii)** isolated flagella (F) and (**iv)** deflagellated cell bodies (CB). Scale bars represent 20 μm. (**C)** Protein gel stained with SYPRO RUBY. Numbers on the left indicate molecular weight in kDa, numbers below indicate cell equivalent of protein loaded on each lane. Each sample lane of F_S_, F_I_, C_S_ and C_I_ was cut into eight pieces and analysed by mass spectrometry (two biological replicates). (**D)** Venn diagram shows total number of all detected proteins (≥ 2 peptides detected, p-value > 0.95) of both replicates.

**Figure 2.**
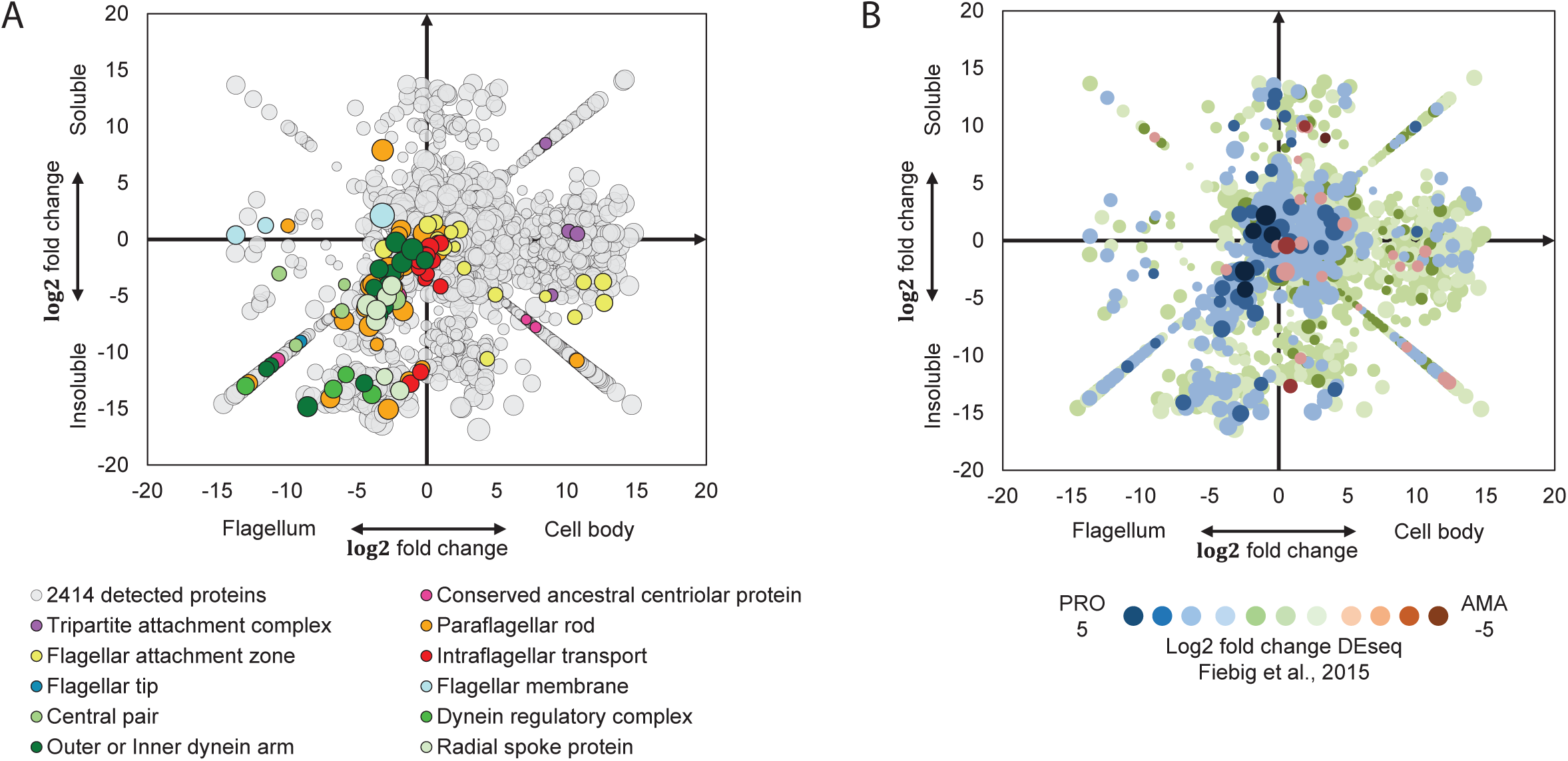
Enrichment plot of flagellar and cell body proteins. Proteins detected by MS were quantified with SINQ. Plotted on the X-axis is the log2 fold-change of the spectral index (C_S_+C_I_ / F_S_+F_I_) and on the Y-axis the log2 fold change of the spectral index (F_S_+C_S_ / F_I_+C_I_). To allow plotting of proteins that were only detected in one fraction, a value of 10^-10^ was inserted for missing spectral indices. The resulting diagonal lines in each quadrant represent proteins uniquely detected in the respective fraction. (**A)** Each data point represents one of 2414 proteins detected in MS run 1 and bubble size reflects protein abundance. Coloured circles indicate representative proteins for different flagellar sub-structures (GeneIDs in S2 Table). The plot can be interactively explored on http://www.leishgedit.net/leishgedit_db/. (**B)** Correlation with RNA-seq data. All 2414 proteins detected in MS run 1 were plotted as in (A) and colour-coded according to the log2 fold change of differentially expressed transcripts between promastigotes (PRO) and amastigotes (AMA) [28].

### Comparison with existing datasets validates flagellar proteome

To validate the data, we mapped well-characterised flagellar proteins onto the enrichment data plot (Figure 2A). Axonemal, paraflagellar rod (PFR), flagellar tip and flagellar membrane proteins mapped to the F_I_ and F_S_ quadrants. Basal body, FAZ and tripartite attachment complex (TAC) proteins mapped to the CB_I_ and CB_S_ quadrants because F fractions contained exclusively the cell-external portion of the flagellum. Intraflagellar transport (IFT) proteins clustered around the midpoint of the plot, indicating their abundance was similar in the F and CB fractions, which is consistent with their known dynamic association with the flagellar basal body and axoneme.

We also found substantial overlaps between *L. mexicana* proteins in the F_I_ quadrant and proteins detected in previously published flagellar proteomes of *L. donovani* and *T. brucei (*S5 Figure A-D). However, *L. mexicana* proteins in the F_S_ quadrant showed only a moderate overlap with reported soluble *T. brucei* flagellar proteins (S5 Figure C). We designed a website (www.leishgedit.net/leishgedit_db) for interactive browsing of proteins in the enrichment plots shown in Figures 2 and S5.

Proteins with predicted trans-membrane domains (TMD) were predominantly detected in the detergent soluble fractions (S5 Figure F). Overall, TMD proteins were underrepresented in the proteome compared to their frequency in the genome (Chi-squared test, p < 0.0001), as were proteins smaller than 10 kDa (S6 Figure), most likely due to well-known technical limitations of protein MS [25]. Although ribosomal proteins were detected in individual F fractions, the enrichment plot clustered them around the midpoint, with many enriched in the cell body fractions (S5 Figure E). Our simple strategy of testing for enrichment thus successfully filtered out likely contaminating proteins from the promastigote flagellar proteome, as recently observed for enrichment of other cytoskeleton structures in *T. brucei* [26,27].

Interestingly, a comparison of these proteomics data with *L. mexicana* RNA-seq data from promastigotes and amastigotes [28] showed that proteins enriched in the flagellar fractions were significantly more likely to have higher RNA abundance in promastigotes vs. amastigotes, compared to proteins detected in the cell body fraction (Figure 2B; Chi-squared test, p < 0.0001). This is consistent with the disassembly of the motile axoneme during differentiation from promastigotes to amastigotes [17]. Whilst on a global scale transcript levels correlate poorly with protein abundance in *Leishmania* spp. [29] these data indicate that modulation of mRNA levels is a key regulatory step in *Leishmania* flagellar biogenesis and differentiation from a 9+2 to a 9+0 flagellum.

### Selection of candidates for a motility mutant screen

Many of the proteins detected in the F fractions had orthologs in previously defined flagellar and ciliary proteomes yet lacked any functional characterisation. Arguably, endowing cells with motility is the primary function of the promastigote flagellum and we took advantage of our high-throughput CRISPR-Cas9 toolkit [22] to identify proteins required for motility and subsequently study the phenotypes of the mutant *Leishmania*. In our knockout (KO) library (S4 Table) we included 19 highly conserved axonemal proteins involved in the regulation of flagellar beating, three intraflagellar transport (IFT) proteins, 60 flagellar proteins with transcript enrichment in promastigotes [28] and eight additional soluble and four insoluble flagellar proteins. Twenty of the selected proteins were detected in the promastigote flagellar proteome but have to our knowledge not been linked to flagella before. We also made deletion mutants for two genes implicated in membrane protein trafficking, *BBS2* and *Kharon1*. Flagellar localisation of a subset of proteins was independently examined by generating cell lines expressing proteins tagged with a fluorescent protein at the N- and/or C-terminus (S7 Figure).

Orthofinder [30] was used to generate genome-wide orthologous protein sequence families using genome sequences of 33 ciliated and 15 non-ciliated species from across eukaryotic life, including *L. mexicana* and *T. brucei (*S5 Table). Twenty-two proteins were kinetoplastid-specific (*L. mexicana* and *T. brucei*), 30 were conserved specifically in ciliated organisms and 23 widely conserved across eukaryotes whilst the remainder showed no clear pattern. In the following, we refer to genes of unknown function by their GeneID from TriTrypDB.org [31] and where we identified named orthologs we used these gene names.

### Screening CRISPR-Cas9 knockout mutants for motility defects

The target genes were then deleted as described previously [22]. To facilitate high-throughput generation of knockout (KO) cell lines, PCR reactions and transfections were performed in 96-well plates. Analysis of drug-resistant transfectants by PCR confirmed loss of the target ORF and integration of the drug-resistance gene in 94 of 98 cell lines (S8 Figure). This 96% success rate highlights the power of our gene deletion strategy. The reason for the presence of the target ORF in the remaining four cell lines was not further investigated, but was confirmed by diagnostic PCR of two independently isolated samples of genomic DNA from the relevant mutants.

The flagellar mutants generated in this study, the previously generated paralysed cell line Δ*PF16* [22], the parental line *L. mex* Cas9 T7, and wild type promastigotes were subjected to motility assays using dark field microscopy to track the swimming behaviour of cells and measure swimming speed and directionality as previously described [32]. Parental cells immobilised though formaldehyde fixation were also measured. More than half of all mutant lines showed a significant deviation from the normal average swimming speed measured for the parental cell line and wild type controls (Figure 3A, B): 52 (53.6%) mutants showed a significant reduction in speed and 4 (4.1%) swam faster (Student’s t-test, p<0.005; Figure 3A, S4 Table). Plotting mean swimming speed against mean directionality (velocity/speed) shows broad groups of mutants (Figure 3B): Those which are paralysed, slower swimmers, slow uncoordinated swimmers, faster swimmers and a single mutant that had faster and more directional swimming (Δ*LmxM.36.3620*). The mechanistic contribution to swimming behaviour remains to be clarified for many proteins in this set. Loss of flagellar waveform modulators would cause altered motility patterns, and this is exemplified by two mutants in this set: the Δ*dDC2* mutant, which lacks the outer dynein arm docking complex protein dDC2 and can perform a ciliary beat but no flagellar beat [33] clusters with the uncoordinated group. By contrast, Δ*LC4-like*, which lacks a distal regulator of outer dynein arms and spends more time doing a flagellar beat at a higher beat frequency [33], was among the faster swimmers.

**Figure 3.**
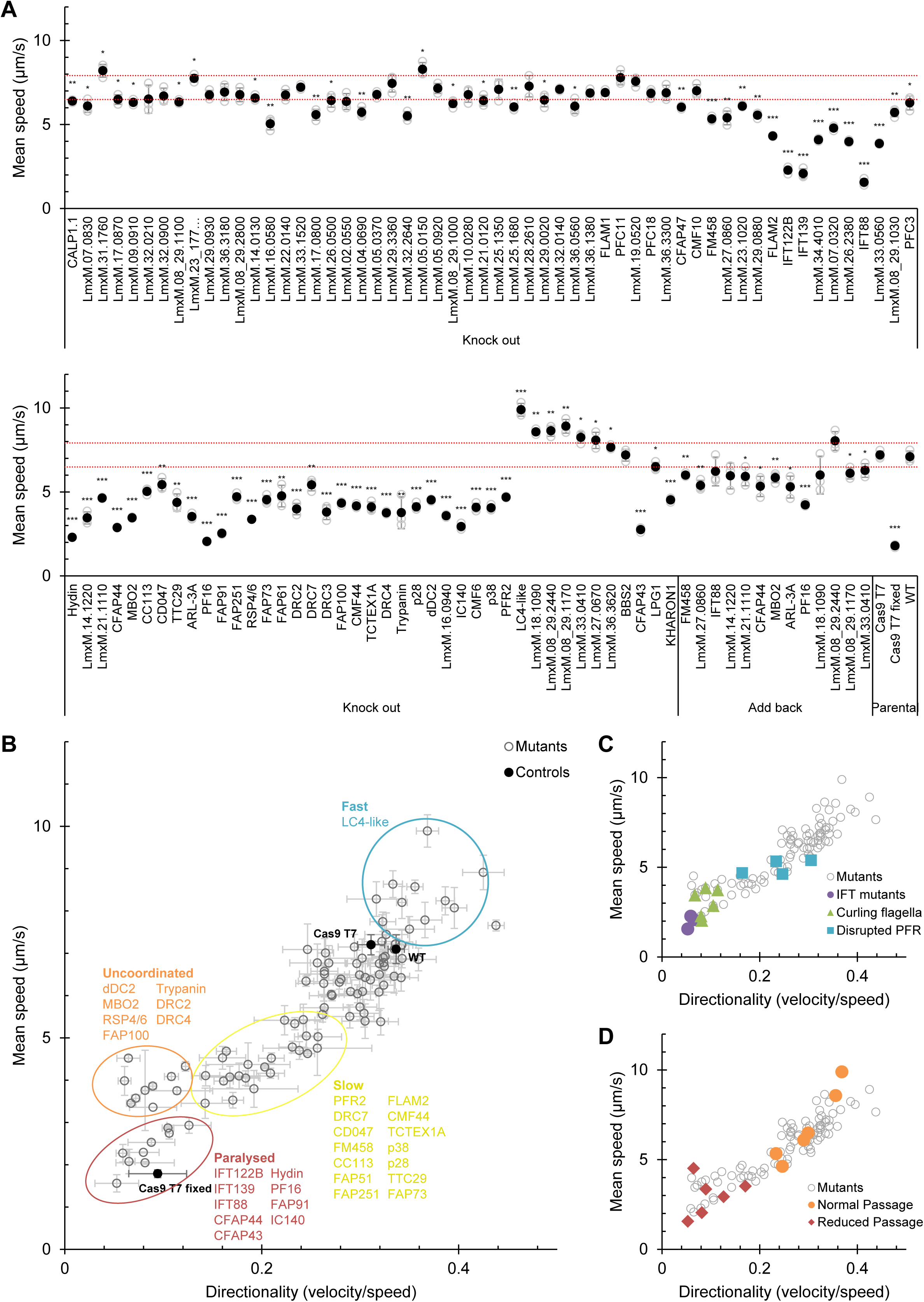
Identification of motility phenotypes. All deletion cell lines generated in this study and the Δ*PF16* mutant [22] were analysed for defects in swimming speed or directionality (the ratio of velocity to speed). (**A)** Mean swimming speeds. Speeds were measured three times and the mean of all three replicates (•) and the individual replicates (○) are shown. Error bars represent the standard deviation. Red dotted lines indicate two standard deviations above and below the parental cell line (Cas9 T7) mean swimming speed. Cas9 T7 cells killed with 1% formaldehyde (Cas9 T7 fixed) were used as a completely immotile control. *** p<0.0005, ** p<0.005, * p<0.05 (Student’s t-test compared to the parental cell line). For a sub-set of mutants, an addback copy of the deleted gene was introduced and swimming speeds restored toward the wild-type. (**B)** The swimming speeds of all knockout mutants (○), as in (A), plotted against mean directionality. Error bars represent the standard deviation of the three replicates. Four main mutant phenotype clusters are apparent: Paralysed (including mutants lacking a long flagellum), uncoordinated (which move slowly, but with greatly reduced directionality), slow (which move slowly, but with reduced directionality and speed) and fast (which move faster, with a tendency for higher directionality). (**C)** Speed and directionality of all knockout mutants in (B) with deletion of IFT components, mutants with a tendency for curling of the flagellum (S9 Figure) or disrupted PFR structure (Figure 5) highlighted. (**D)** Speed and directionality of knockout mutants in (B) with those passaged though sand flies (Figure 6) highlighted.

The most severe loss of motility was observed in three cell lines that had no visible external flagellum (Figure 4); all of these were deletions of conserved intraflagellar transport (IFT) proteins (Δ*IFT122B*, Δ*IFT139* and Δ*IFT88*). Ablation of the central pair (CP) protein hydin also resulted in almost complete paralysis, comparable to the deletion of the CP protein PF16 [22].

**Figure 4.**
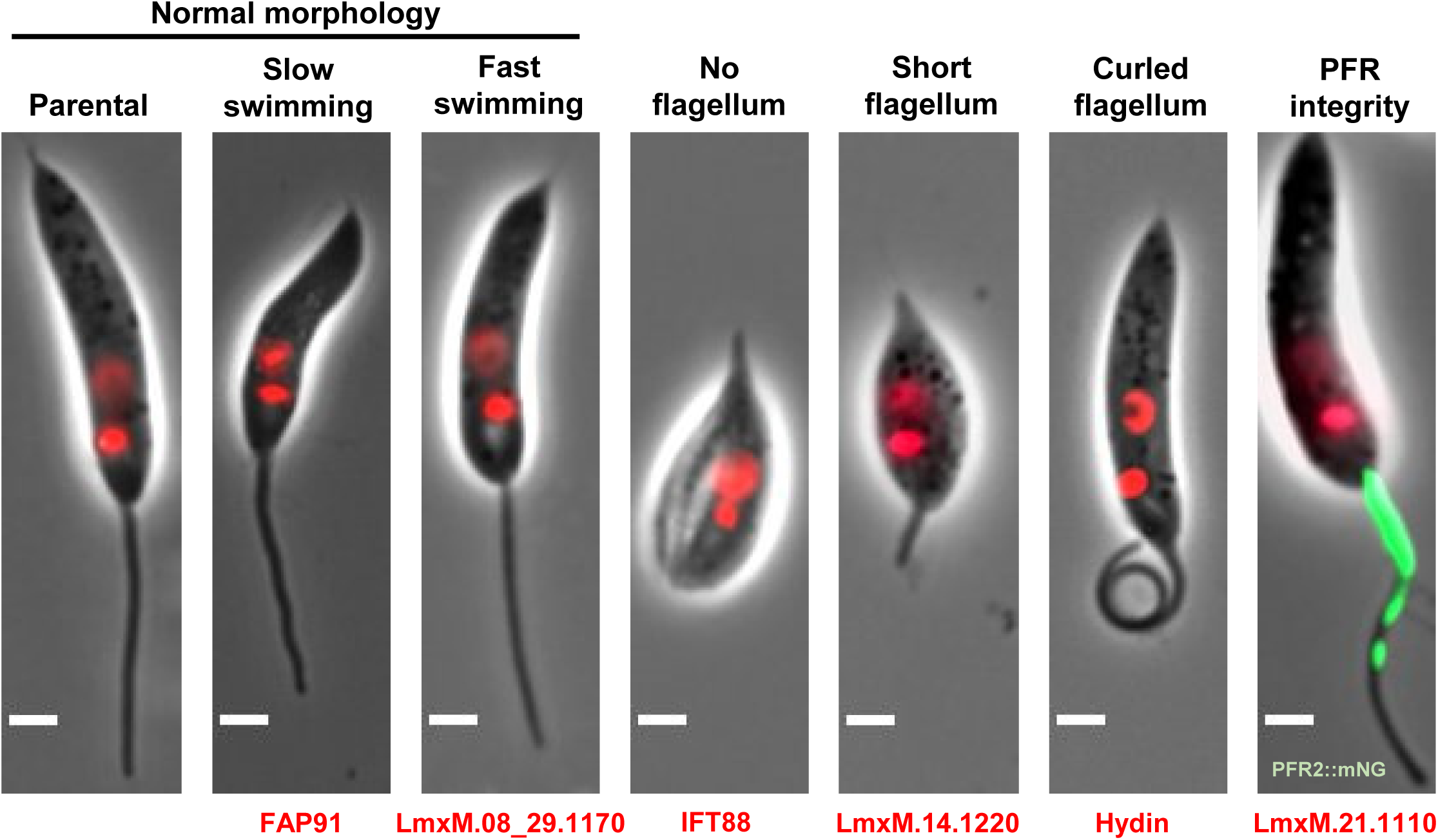
Phenotype categories. Categories of mutant phenotypes: among cells with normal morphology there were normal swimmers as well as slower or faster swimmers. Four categories of distinct morphological defects were observed, which all lead to impaired motility: no flagellum, short flagellum, curled flagellum and disrupted PFR. One representative mutant is shown for each category, the deleted gene is indicated. Green, PFR2∷mNG signal; red, Hoechst-stained DNA. Scale bar, 2 μm.

In a subset of paralysed or slow-swimming uncoordinated mutants (Figure 3C) we noted that the flagella tended to be in a curled rather than straight conformation. Δ*hydin* mutants had the highest proportion of curled-up flagella (62.6%, Figure 4 and S9 Figure) while fewer than 1% of flagella were curled-up in the parental cell line and many other slow swimming mutants (S9 Figure). A high proportion (>10%) of curled-up flagella was also found in four paralysed KO lines (inner dynein arm intermediate chain protein mutant Δ*IC140*, 57%; Δ*PF16*, 14%; tether and tether head complex protein mutants Δ*CFAP44*, 15% and Δ*CFAP43*, 19%) and three uncoordinated KO lines (Δ*MBO2*, 26%; nexin-dynein regulatory complex protein mutant Δ*DRC4*, 13%; Δ*LmxM.33.0560*, 12%). The curls were observed in aldehyde fixed cells as well as in live cells in culture, indicating they were not an artefact of microscopy sample preparation. This novel phenotype might be caused by disrupted dynein regulation and warrants further investigation.

Null mutants for the major PFR protein PFR2, lacking the paracrystalline PFR lattice structure, are known to have impaired motility [34]. To compare motility of a Δ*PFR2* mutant with other mutants generated in this study, we used CRISPR-Cas9 to delete both allelic copies of the *PFR2* array (*PFR2A, PFR2B* and *PFR2C*) and confirmed loss of PFR2 expression by Western blot (S10 Figure). This Δ*PFR2* line had slower and less directional swimming compared to the parental cells, clustering with other slow swimming mutants defined in Figure 3B. To test whether gene deletion in other slow swimming mutants had a major disruptive effect on the PFR, which might explain their motility defect, we expressed PFR2∷mNG in KO lines and looked for changes to PFR length or loss of PFR integrity (defined as gaps in the PFR2∷mNG signal) (Figures 4 and 5). Three mutants had shorter flagella compared to the parental cell line, but the PFR remained proportional to the overall flagellar length and was uninterrupted (Δ*ARL-3A*, Δ*CFAP44*, and Δ*FLAM2*). Six mutants had PFR-specific defects (Figure 5B): a shorter flagellum with a disproportionately shorter PFR (Δ*LmxM.27.0860;* Δ*TTC29;* Δ*LmxM.14.1220*), a normal-length flagellum with a shorter PFR (Δ*FM458*) or a shorter PFR with gaps (Δ*LmxM.21.1110,* 25.3% of all flagella; Δ*MBO2* only 4.1% of all flagella). Interestingly, these comparatively subtle alterations to PFR length and integrity reduced swimming speed to similar levels as PFR2 deletion (Figure 3C).

**Figure 5.**
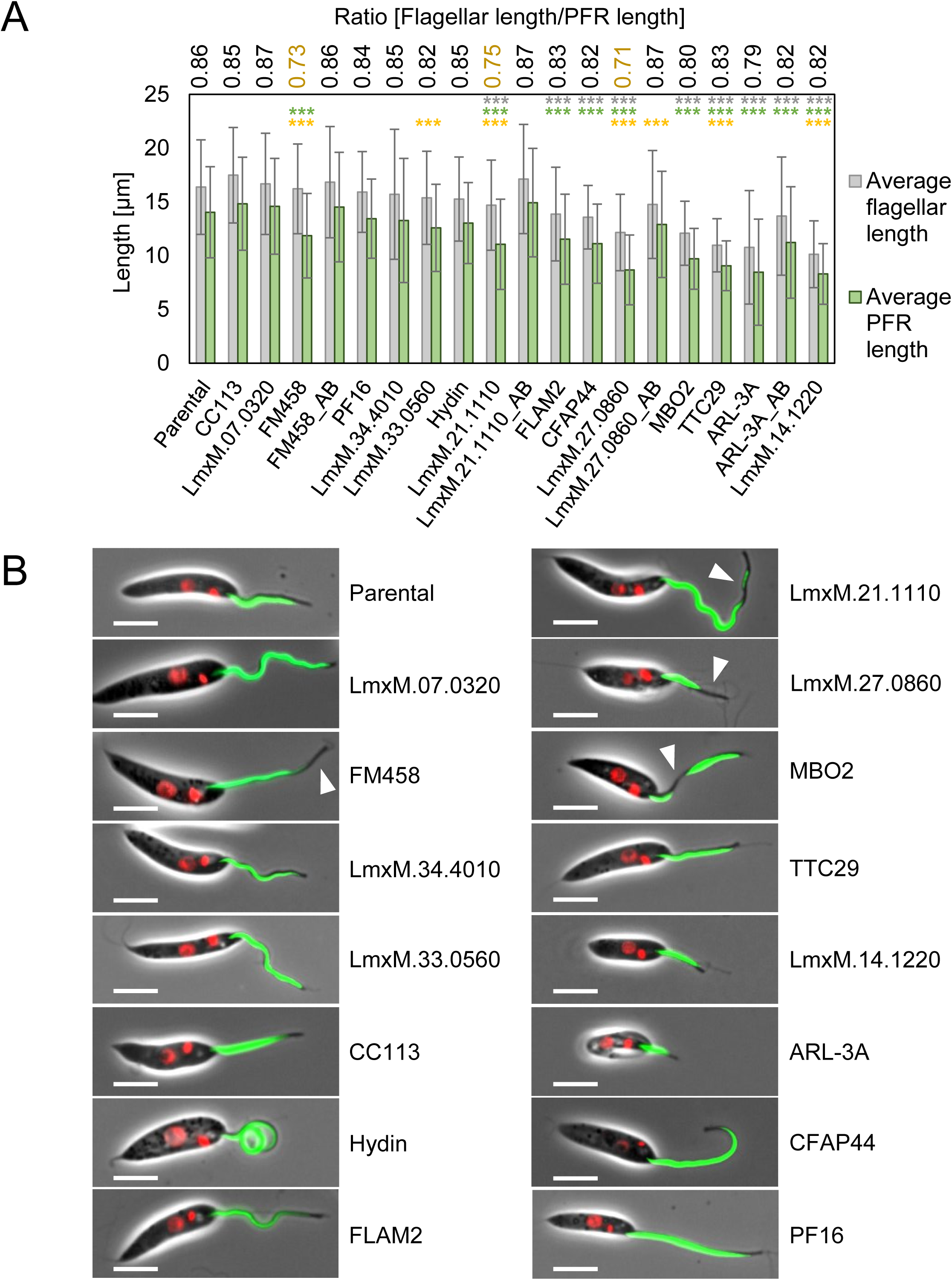
Measurement of flagellar and PFR length in motility mutants. **(A)** Measurements of flagellar length (measured from kinetoplast DNA to flagellar tip; grey bar) and PFR2∷mNG signal (green bar; data in S8 Table). Error bars show standard deviation. At least 70 measurements were recorded per cell line. The GeneIDs / gene names indicate the deleted gene. Numbers above bars show PFR : flagellar length ratio. Measurements were compared to the parental cell line expressing PFR2∷mNG and p-values (Students t-test) for flagellar length (grey) and PFR length (green) are indicated: *** p≤0.001. (**B)** Fluorescence micrographs showing tagged mutant cell lines used for measurements in (A). Green, PFR2∷mNG signal; red, Hoechst-stained DNA. Scale bar, 5 μm. White arrows indicate PFR defects.

We generated 13 add-back cell lines to rescue mutant phenotypes by transfecting episomes containing the deleted ORF. Four complemented mutants fully recovered parental swimming speed (Figure 3) and complemented Δ*CFAP44* and Δ*MBO2* lines showed fewer curled flagella (S9 Figure). Complementation of the other slow swimming mutants resulted in a significant increase in swimming speed close to parental levels (Figure 3) and reduction of curling compared to the KO lines (S9 Figure).

Thus, our screen readily identified promastigote mutants with impaired motility and even the most severe phenotype, ablation of flagellar assembly caused by loss of IFT components, was compatible with promastigote survival *in vitro*, in line with earlier reports [35], [36], [37].

### Paralysed promastigotes are cleared from sand flies

Whilst flagellar motility is generally believed to be required for development in sand flies, enabling *Leishmania* migration from the midgut to the mouthparts [38-40], this has not been directly tested. To interrogate the phenotypes of larger cohorts of *Leishmania* mutants in parallel, we developed a multiplexed bar-seq strategy inspired by pioneering phenotyping screens in yeast [41] and the malaria parasite *Plasmodium berghei* [42,43]. We pooled mutant *L. mexicana* lines that were each tagged with a unique 17 bp barcode. This enabled us to measure the relative abundance of each line at different time points after sand fly infection (S11 Figure). Seventeen were flagellar mutants described above and five were parental control cell lines tagged with unique barcodes in their small subunit (SSU) ribosomal RNA locus. We also generated a barcoded Δ*LPG1* KO mutant, which is only defective in LPG synthesis [22,44] and three barcoded mutants defective in the pathway leading to mannose activation for synthesis of LPG and other glycoconjugates: KOs of phosphomannose isomerase [45] (Δ*PMI*), phosphomannomutase [46] (ΔPMM) and GDP-mannose pyrophosphorylase [47] (ΔGDP-MP). These mutants were included as control lines expected to be outcompeted by the parental cell lines. The barcoded cell lines were pooled in equal proportions and used to infect *L. longipalpis*. The relative abundance of each line was determined by sequencing DNA isolated from the mixed promastigote pool and from flies at two, six and nine days after infection. The results show progressively diminishing proportions for the control mutants defective in LPG synthesis (Δ*LPG1*) or a broader range of glycoconjugates including LPG (Δ*PMI*, Δ*PMM* and Δ*GDP-MP*) (Figure 6, S9 Table) indicating that parasites lacking these molecules were at a competitive disadvantage in these infections. This effect was apparent as early as two days after infection, consistent with a protective role for PG-containing glycoconjugates in the digesting bloodmeal [48] and a role for LPG in *L. mexicana* attachment to *L. longipalpis* [49].

**Figure 6.**
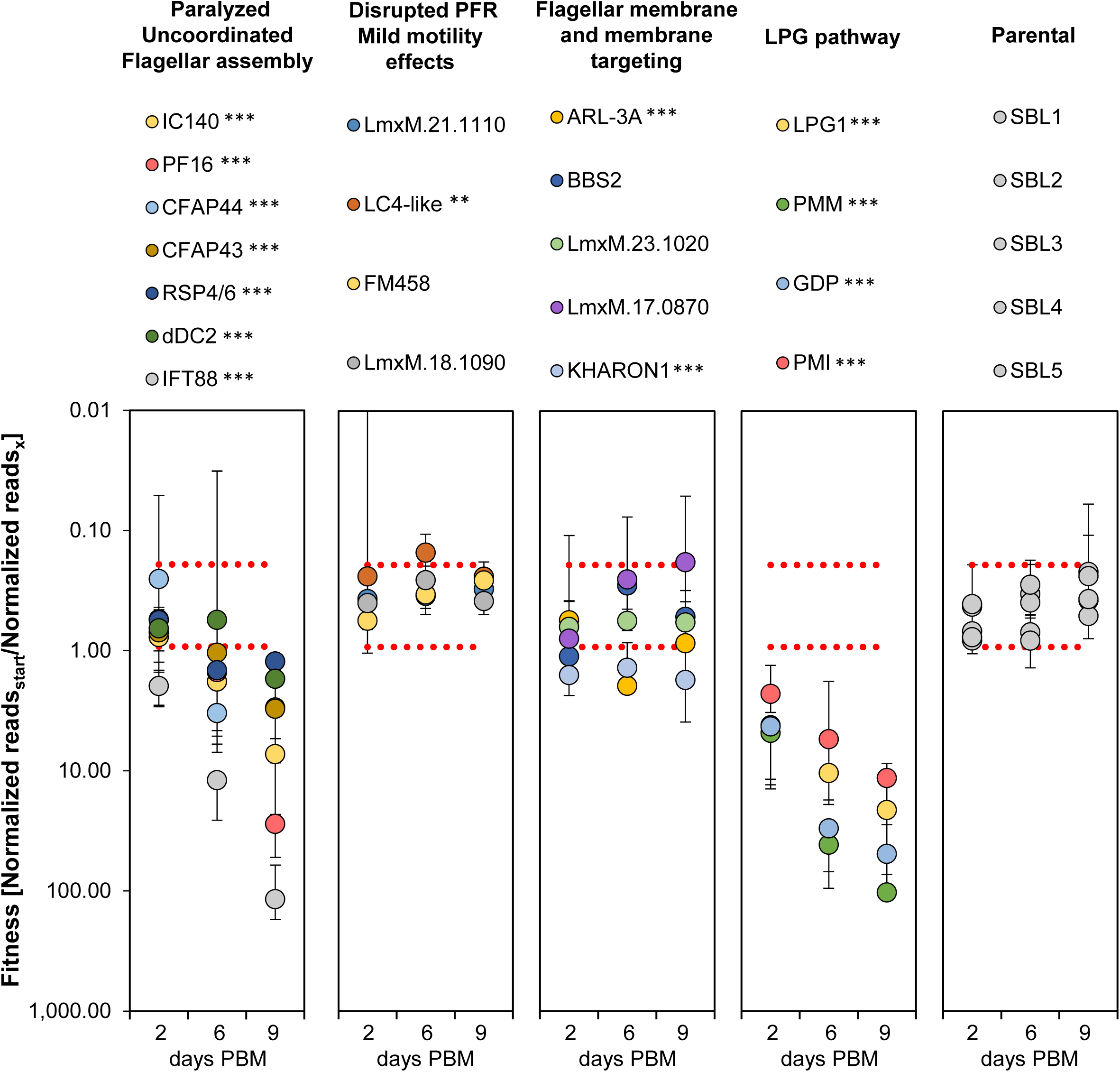
Relative fitness of *Leishmania* mutants in *L. longipalpis* infections. The plots display abundance of barcodes at time points post bloodmeal (PBM) relative to the abundance of this barcode in the mixed parasite population used to infect *L. longipalpis*. Mutants are grouped according to the function of the deleted gene and severity of motility phenotype (**A**) flagellar mutants with severe motility defects (paralyzed, uncoordinated swimmers and aflagellate cells), (**B**) flagellar mutants with mild motility defects, (**C**) mutants lacking flagellar membrane proteins or proteins involved in protein trafficking to the flagellar membrane, (**D**) mutants lacking key enzymes for the synthesis of LPG and other glycoconjugates, (**E**) control mutants with wild type motility. Data points represent the average of three replicates. Error bars show the standard deviation of the mean of the three replicates. Dotted red lines indicate two standard deviations above and below the parental cell line (SBL1-5). Measurements were compared (two-sided t-test) to the average of all five parental controls and p-values are indicated: *≤0.05, **≤0.005, ***≤0.0005.

Paralysed and uncoordinated mutants also became noticeably scarcer as the infection progressed (Figure 6, S9 Table). The aflagellate Δ*IFT88* mutant showed the most severe phenotype and a significant decrease over time was also measured for Δ*PF16,* Δ*CFAP43,* Δ*CFAP44,* Δ*IC140,* Δ*dDC2* and Δ*RSP4/6.* By contrast, mutants with a mild swimming defect (slower swimmers Δ*LmxM.21.1110,* Δ*FM458* and Δ*LmxM.18.1090* and faster Δ*LC4-like*) (Figure 3D) remained as abundant as the normal swimmers throughout the infection (Figure 6, S9 Table). The exceptions were the slower swimmers Δ*Kharon1,* which is also defective in the transport of a flagellar glucose transporter [50], and Δ*ARL-3A,* which has a short flagellum (Figure 5). Both of these were rarer in the fly compared to the starting pool. To gain anatomical resolution and determine whether an immotile mutant fails to migrate to anterior portions of the fly gut, we infected separate batches of *L. longipalpis* with motile parasite lines and complemented KO lines as controls, with the motile Δ*BBS2* mutant, which lacks a component of the BBSome complex [51] which is expected to play a role in flagellar membrane trafficking, and with the paralysed Δ*PF16* mutant (Figure 7). The Δ*PF16* mutants are among the least motile cells that retain a long flagellum (Figure 5), while having only moderate levels of flagellar curling (S9 Figure). The axonemal defect resulting in paralysis is a well-characterised disruption of the central pair in kinetoplastids (Figure 3B and [22,52,53]) and is similar to the defect of the *pf16 Chlamydomonas reinhardtii* mutant [54] indicating it is a well-conserved core axoneme component. Two days post blood-meal (PBM), the *L. mexicana* wild type and *L. mex* Cas9 T7 [22] control cell lines and the Δ*BBS2* mutant developed well, with infection rates above 70%; the Δ*PF16* mutant produced the lowest infection rate (below 50%). The introduction of an add-back copy of *PF16* into the Δ*PF16* line restored infection levels (Figure 7A). In all lines, promastigotes were localized in the abdominal midgut, within the bloodmeal enclosed in the peritrophic matrix (Figure 7B). After defecation (day 6 PBM), all control lines and the Δ*BBS2* mutant replicated well and developed late-stage infections with colonisation of the whole mesenteron including the stomodeal valve (Figure 7B) which is a prerequisite for successful transmission. Their infection rates ranged from 56% to 83%. By contrast, Δ*PF16 Leishmania* failed to develop; the infection rate was less than 2% (a single positive fly out of 62 dissected (Figure 7A), with parasites restricted to the abdominal midgut (Figure 7B)), indicating that Δ*PF16* parasites were lost during defecation and were unable to develop late stage infections in *L. longipalpis*. Our data provide strong evidence that flagellar motility is an essential requirement for successful development in sand flies and, by implication, transmission.

**Figure 7.**
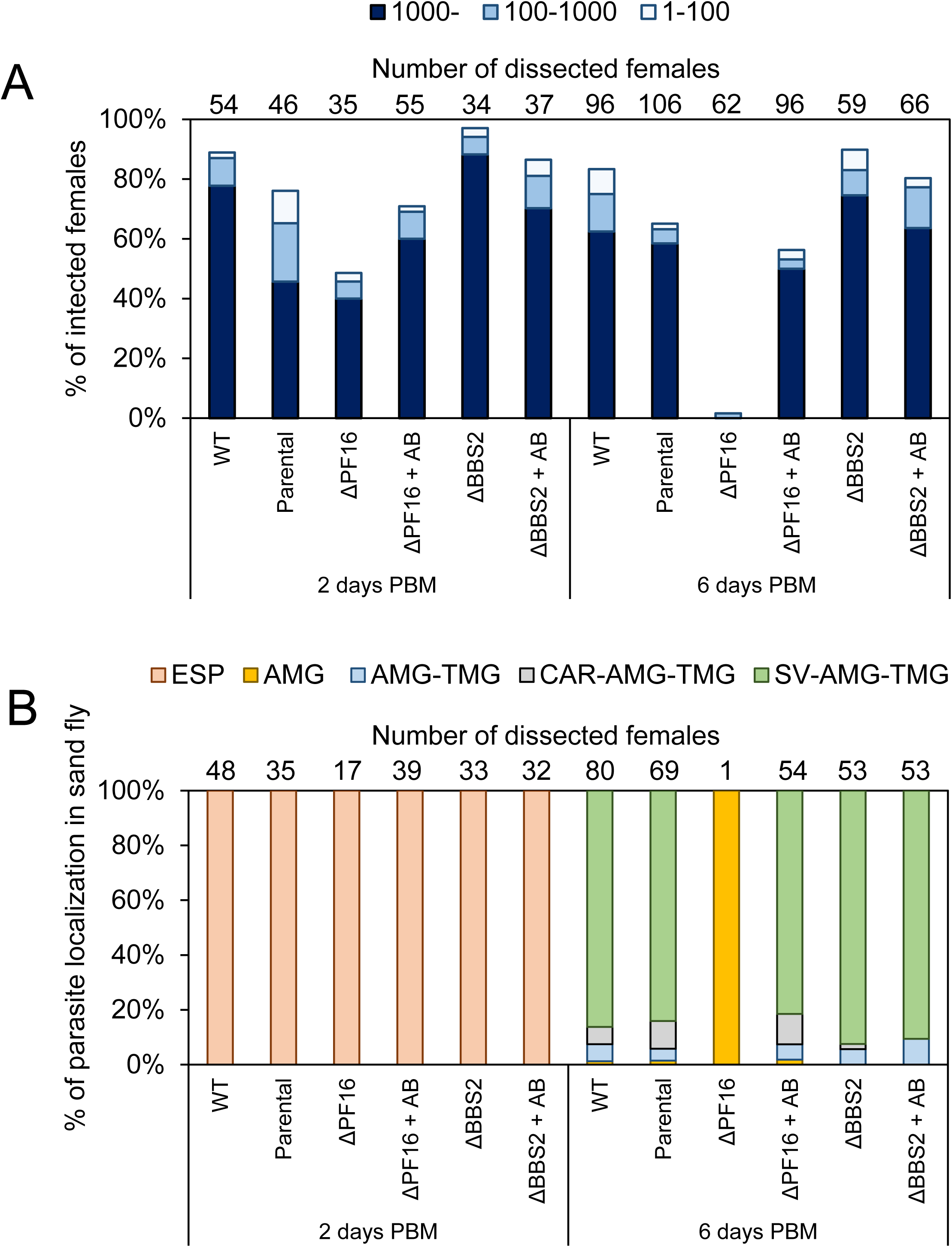
Development of *L. mexicana* ΔPF16 in *L. longipalpis*. **(A)** Infection rates (% of infected females) and intensities of infections (four categories) of *L. mexicana* in *L. longipalpis*. Numbers above the bars indicate numbers of dissected females in the group. (**B)** Localization of *L. mexicana* infections in *L. longipalpis*. ESP: endoperitrophic space, AMG: abdominal midgut, TMG: thoracic midgut, CAR: cardia, SV: stomodeal valve. Numbers above the bars indicate numbers of infected females in the group.

## Discussion

This study demonstrates the power of high-throughput CRISPR-Cas9 knockout screens to discover mutant phenotypes in *Leishmania*. We defined a high-confidence flagellar proteome and used these data in conjunction with transcriptomics data and prior knowledge of conserved axonemal proteins to demonstrate a role in motility for >50 genes from a set of one hundred. We also show the importance of flagellar motility in the colonisation of sand flies. The data from the pooled mutant population show a progressive loss of paralysed or uncoordinated swimmers over nine days from infection. Because these data report total abundance of each genotype in the whole fly without discriminating between regions of the gut, we probed this question further in infections with the Δ*PF16* mutant, which is essentially paralysed and incapable of sustained directional motility due to a defined defect in the central pair complex of the axoneme [22]. The results show that Δ*PF16 Leishmania* were rapidly lost from most of the dissected flies, consistent with the depletion of this mutant from the mixed pool, and additionally shows that none of the few remaining parasites reached anterior parts of the alimentary tract. These findings show that parasite motility is required for completion of the *Leishmania* life cycle, in line with the essential role of motility in other vector-transmitted protists. For example, Rotureau *et al.*, [55] showed that loss of forward motility, caused by ablation of outer dynein arms though KO of *DNAI1*, rendered *T. brucei* unable to reach the tsetse fly foregut. It seems likely that loss of motility also contributed to the inability of *L. amazonensis* to progress beyond the abdominal midgut of *L. longipalpis* when the parasites overexpressed GTP-locked ADP-ribosylation factor-like protein 3A (Arl-3A) and as a result grew only short flagella [56].

Observations of attached *Leishmania* in dissected sand flies show adhesion specifically via the flagellum but the precise molecular interactions between flagellum and the microvillar gut lining remain to be clarified. The dominant cell surface glycoconjugate LPG which covers the entire parasite surface including the flagellum is known to be important in *Leishmania* attachment to sand fly guts [57] and our results support the view that LPG plays an important role in *L. mexicana* infection of *L. longipalpis* [49]. The proportion of Δ*LPG1* mutants had decreased by two days after infection and reduced further as infection progressed. The observed loss of fitness of the Δ*PMM*, Δ*GDP-MP* and Δ*PMI* mutants is likely the cumulative effect of the loss of LPG and a broader range of mannose-containing glycoconjugates which were shown to protect *Leishmania* in the digesting bloodmeal [48]. The consistency of the pooled mutant data with the reported phenotypes of individual glycoconjugate-deficient mutants demonstrates the power of this new rapid method for mutant phenotyping in *Leishmania*. However, whilst a role for LPG in *L. mexicana* attachment to the fly is well established, the possible contribution of flagellum-specific surface molecules [58] has not yet been conclusively resolved. Zauli et al., [36] reported isolation of *L. braziliensis* from a patient’s skin lesion which differentiated to promastigotes with an “atypical” morphology. These cells had a short flagellum barely protruding from the flagellar pocket, with an amorphous tip suggestive of a defect in flagellum elongation. In experimental infections of *L. longipalpis*, these parasites persisted in the fly following defecation of the blood meal, suggesting that they remained sufficiently anchored without a long flagellum. It would be interesting to follow up the subsequent development of this mutant in the fly.

Several lines of evidence suggest a role for the trypanosomatid flagellum in environmental sensing [40,59-61]. Evidence for specific signal transduction pathways aiding promastigote navigation through the sand fly is however limited. Cyclic nucleotide signal transduction pathways may have important roles in coupling environmental sensing with regulation of flagellar beat patterns [62,63]. In our flagellar proteome we identified several adenylate cyclases (ACs), cAMP-specific phosphodiesterases (PDEs) and PKA subunits and mapped their localisations to distinct flagellar subdomains by protein tagging (S7 Figure). The motility assays showed that deletion of PKA subunits (Δ*LmxM.34.4010 (*partial KO only) and Δ*FM458*) reduced swimming speed, whereas deletion of two different PDEs (Δ*LmxM.18.1090* and Δ*LmxM.08_29.2440*) increased it, pointing to an activating role for cAMP in *Leishmania* motility. Knockout of receptor-type adenylate cyclase a-like protein LmxM.36.3180 had no effect on swimming speed in our motility assay but given the possible redundancy with other flagellar ACs, this preliminary finding should be followed up by examination of other AC mutants individually and in combinations. In our pooled KO screen in sand flies, KOs of PDE LmxM.18.1090 and PKA RSU (FM458) remained as abundant as the controls, indicating that the mild motility phenotypes measured *in vitro* did not significantly impair colonisation of flies.

Perturbation of the flagellar membrane might be expected to interfere with sensory functions mediated through the flagellum. Ablation of membrane proteins LmxM.17.0870 and LmxM.23.1020 (S7 Figure) did not significantly enhance or reduce the relative abundance of the respective mutants in sand flies over the nine-day observation period. BBS2 is an integral part of the core BBSome complex which is highly conserved across ciliated eukaryotes [64] and functions as a cargo adaptor for ciliary membrane protein trafficking in *Chlamydomonas* flagella and metazoan cilia [65]. Our pooled mutant data and infections with the *BBS2* deletion mutant alone found that loss of this gene had no discernible detrimental effect on survival in sand flies and the parasites’ ability to reach the anterior gut. By contrast, KO of Kharon1, a protein shown to be required for trafficking of the glucose transporter LmGT1, and perhaps other proteins, to the promastigote flagellum [50] led to slightly reduced fitness in the flies from the earliest time point. The Δ*Arl-3A* mutants were also less abundant compared to the controls. This is reminiscent of the previously published abortive phenotype of *L. amazonensis* overexpressing the constitutively GTP bound *Ld*ARL-3A-Q70L [56]. This mutant formed only a short flagellum, similar to the Δ*Arl-3A* mutant generated in the present study. Failed attachment as a result of the shortened flagellum was thought to be a likely cause for the rapid clearance of *Ld*ARL-3A-Q70L-expressing parasites but it was noted that an inability to migrate at later stages of development would also lead to the disappearance of the mutants [56]. In our study the phenotype of the Δ*Arl-3A* mutants was however mild compared to the aflagellate (Δ*IFT88*) or paralysed mutants. Arl-3A acts as guanine nucleotide exchange factor in the transport of lipidated proteins to the flagellar membrane [66] and protein mis-targeting could contribute to the phenotype in addition to flagellar shortening. Further insights into the contribution of flagellar membrane proteins to attachment or directional swimming behaviour may be uncovered by further biochemical studies into flagellar membrane composition and subjecting different mutants (with or without overt motility phenotypes in culture) to chemotaxis assays and fly infections.

In contrast to the absolute requirement of motility for movement through the sand fly vector, flagellar motility is dispensable for promastigote proliferation in culture. Promastigotes are viable and able to divide even if they fail to assemble a flagellum at all, as demonstrated originally by the deletion of cytoplasmic dynein-2 heavy chain gene *LmxDHC2.2* [35] and IFT140 [37] and the phenotypes of knockouts of anterograde and retrograde IFT components in the present study. The ensuing prediction that most gene deletions affecting flagellar function are expected to yield viable promastigotes in the laboratory is borne out by our high success rate of obtaining 96% of attempted knockouts. Thus, in *Leishmania*, flagellar mutant phenotypes can be observed in replication-competent cells over many cell cycles and our mutant library enables detailed systematic studies of KO phenotypes to probe protein functions in flagellum assembly, motility and signal transduction.

A fruitful area for further studies will be dissection of PFR function and assembly mechanisms. This extra-axonemal structure is required for motility as demonstrated through deletion of the major structural PFR components, PFR1 and PFR2 in *Leishmania* [34,67] and ablation of PFR2 by RNAi in *T. brucei* procyclic forms [68] but its precise role remains unclear. The PFR comprises more than 40 proteins, some with structural roles, others with roles in adenine nucleotide homeostasis, cAMP signalling, calcium signalling and many uncharacterised components [69,70] and it may anchor metabolic and regulatory proteins as well as influencing the mechanical properties of the flagellum. Our results showed that fragmentation of the PFR caused by loss of LmxM.21.1110 reduced swimming speed to levels similar to the structurally more severe PFR2 KO. Whether LmxM.21.1110 is required for correct PFR assembly or stabilisation is currently unknown.

Motility mutants analysed in our screen also included deletions of genes with human orthologs linked to ciliopathies (such as *hydin*) or male infertility (*CFAP43* and *CFAP44*) [71]. *Leishmania* offers a genetically tractable system to gain further mechanistic insight into their functions. The *hydin* mutant has been extensively characterised in other species: in mammals, mutations in the *hydin* gene cause early-onset hydrocephalus [72] and subsequent studies on *C. reinhardtii, T. brucei* and mice showed that hydin localises to the C2 projection of the central pair complex [73], and that loss of hydin function causes mispositioning and loss of the CP [74] and motility defects [73-75]. The motility phenotype in the *L. mexicana* Δ*hydin* mutant was consistent with these existing data and we made the new observation that the mutant flagella show extensive curling (Figure 4, S9 Figure). Interestingly, *hydin* knockdown in *C. reinhardtii* caused flagella to arrest at the switch point between effective and recovery stroke, leaving cells with one flagellum pointing up and the other down, prompting speculation that this may indicate a role for hydin in signal transmission to dynein arms [73]. Consistent with this hypothesis, cilia of *hy3/hy3* mouse mutants frequently stalled at the transition point between the effective and recovery stroke [75]. Curling may represent the failure of flagellum bending to reverse during progression of the normal flagellum waveform down the flagellum, leaving the flagellum locked at one extreme of bending, analogous to the ciliary beat *hydin* phenotype. In *L. mexicana*, the Δ*hydin* mutant presented the most severe manifestation of the curling phenotype, which was also observed in a lower proportion of other mutants (S9 Figure). This phenotype may be a consequence of mis-regulated dyneins and the set of mutants exhibiting curling will facilitate further experiments to establish the mechanistic basis for flagellar curling.

Genetic, biochemical and structural studies have provided elegant and detailed models for the mechanisms of flagellar motility [76,77]. Phylogenetic profiling and comparative proteomics studies have yielded insights into the evolutionary history, core conserved structures and lineage-specific adaptations of eukaryotic flagella. Our CRISPR-Cas9 KO method enables rapid targeted gene deletion and characterisation of loss-of-function phenotypes for large cohorts of *Leishmania* genes *in vitro* and *in vivo* and hence new opportunities to interrogate the functions of hitherto poorly characterised flagellar proteins in motility regulation, environmental sensing and axoneme remodelling from 9+2 to 9+0. The bar-seq strategy for phenotyping of mutants can also be used to probe parasite-host interactions in mammals.

## Materials and Methods

### Cell culture

Promastigote-form *L. mexicana (*WHO strain MNYC/BZ/62/M379) were grown at 28°C in M199 medium (Life Technologies) supplemented with 2.2 g/L NaHCO_3_, 0.005% haemin, 40 mM 4-(2-Hydroxyethyl)piperazine-1-ethanesulfonic acid (HEPES) pH 7.4 and 10% FCS. The modified cell line *L. mexicana SMP1:TYGFPTY* [17] was cultured in supplemented M199 with the addition of 40 μg/ml G-418 Disulfate. L. *mex* Cas9 T7 [22] was cultured in supplemented M199 with the addition of 50 μg/ml Nourseothricin Sulphate and 32 μg/ml Hygromycin B.

### Deflagellation protocol

To avoid proteolytic degradation, all procedures were performed on ice. 2·10^9^ *L. mexicana SMP1:TYGFPTY* cells were collected at 800*g* for 15 min at 4°C, washed once in 20 ml phosphate buffered saline (PBS) and resuspended in 5 ml 10 mM PIPES [10 mM NaCl, 10 mM piperazine-N,N′-bis(2-ethanesulfonic acid, 1 mM CaCl_2_, 1 mM MgCl_2_, 0.32 M sucrose, adjusted to pH 7.2]. 0.375 ml of 1 M Ca^2+^ solution (final conc. 0.075 M) and a protease inhibitor cocktail [final concentration, 5 μM E-64, 50 μM Leupeptin hydrochloride, 7.5 μM Pepstatin A and 500 μM Phenylmethylsulfonyl fluoride (PMSF)] were added to the cell suspension. Cells were deflagellated by passing them 100 times through a 200 μl gel loading pipette tip (Starlab) attached to a 10 ml syringe. Flagella and cell bodies were separated through density gradient centrifugation, using a modified version of the protocol in [78]. The sample was loaded on top of the first sucrose-bed containing three layers of 10 mM PIPES with 33% (upper), 53% (middle) and 63% (bottom) w/v sucrose [10 mM NaCl, 10 mM piperazine-N,N′-bis(2-ethanesulfonic acid, 1 mM CaCl_2_, 1 mM MgCl_2_, adjusted to pH 7.2 with either 0.96M, 1.55M or 1.84M sucrose] and centrifuged at 800*g* for 15 min at 4°C. The pellet in the 63% sucrose layer was diluted with 10 ml 10 mM PIPES and centrifuged at 800*g* for 15 min at 4°C. The supernatant was discarded and the pellet resuspended in 40 μl 10 mM PIPES. This was the cell body fraction. The top layer of the first sucrose-bed, containing flagella, was collected and sucrose sedimentation was repeated with a second sucrose-bed containing only one layer of 10 mM PIPES with 33% w/v sucrose. The resulting top layer of the second sucrose bed was transferred to an ultra-centrifugation tube (Beckmann tubes) and collected by ultra-centrifugation at 100,000*g* for 1 h at 4°C (Beckman Coulter). The supernatant was discarded and the pellet resuspended in 40 μl 10 mM PIPES. This was the flagellar fraction. All other sucrose layers contained a mixture of flagella and cell bodies and were discarded. 1 μl of flagellar and cell body fractions was used for counting and imaging and 36 μl of each fraction were used for proteomic analysis.

### Protein Mass spectrometry

Cell body and flagellar fractions were supplemented with 4 μl protease inhibitor cocktail (see above) and 10 μl octylglycoside (1% (w/v) final conc.), incubated for 20 min on ice and centrifuged at 18,500*g* for 1 h at 4°C to separate soluble (supernatant) from insoluble (pellet) proteins. 50 μl ice cold reducing 2x Laemmli buffer was added to the resulting supernatant. Pellets were dissolved in 100 μl 1x Laemmli buffer. To avoid aggregation of hydrophobic proteins, fractions were not boiled prior to SDS-PAGE [79]. 20 μl of flagella fractions and 10 μl of cell body fractions (∼5 – 20 μg protein) were pre-fractionated on a 10% polyacrylamide gel, stained overnight with SYPRO Ruby Protein Gel Stain (Molecular Probes) and destained for 30 min in 10% (v/v) Methanol / 7% (v/v) acetic acid. Sample preparation in the following was carried out as described in [80]. Briefly, gel pieces were destained with 50% acetonitrile, reduced with 10mM TCEP (Tris(2-carboxyethyl)phosphine hydrochloride) for 30 minutes at RT, followed by alkylation with 55 mM Iodoacetamide for 60 minutes in the dark at RT. Samples were deglycosylated with PNGase F over two days at RT and digested overnight at 37°C with 100 ng trypsin. Samples were acidified to pH 3.0 using 0.1% trifluoroacetic acid and desalted by reversed phase liquid chromatography. Samples were analysed on an Ultimate 3000 RSLCnano HPLC (Dionex, Camberley, UK) system run in direct injection mode coupled to a QExactive Orbitrap mass spectrometer (Thermo Electron, Hemel Hempstead, UK).

### MS data analysis and SINQ enrichment plot

MS-data were converted from .RAW to .MGF file using ProteoWizard (S6 Table) and uploaded to the Central Proteomics Facilities Pipeline (CPFP [81]). Protein lists were generated by using CPFP meta-searches (S6 Table) against the predicted *L. mexicana* proteome (gene models based on [28], followed by label-free SINQ quantification (S1 and S6 Table). For SINQ enrichment plots detected geneIDs were filtered (p ≥ 0.95, ≥ 2 peptides) and plotted using normalized spectral indices. For missing indices pseudo spectral indices of 10^-10^ were inserted. The mass spectrometry proteomics data have been deposited to the ProteomeXchange Consortium via the PRIDE [82] partner repository with the dataset identifier PXD011057.

### Gene tagging

Tagging was achieved by insertion of a drug-selectable marker cassette and fluorescent protein gene into the endogenous gene to produce an in-frame gene fusion. The fusion PCR method described in Dean *et al*., [83] was used for tagging with eYFP, using pJ1170 (pLENT-YB) as the template plasmid for PCR and selection with 5 μg/ml Blasticidin-S deaminase. The CRISPR-Cas9 method described in Beneke et al., [22] was used for tagging with mNeonGreen. The online primer design tool www.LeishGEdit.net was used to design primers for amplification of the 5’ or 3’ sgRNA template and primers for amplification of donor DNA from pPLOTv1 blast-mNeonGreen-blast or pPLOTv1 puro-mNeonGreen-puro. Transfectants were selected with either 5 μg/ml Blasticidin-S deaminase or 20 μg/ml Puromycin Dihydrochloride.

### CRISPR-Cas9 gene knockouts

Gene deletions were essentially done as described in Beneke et al., [22]. The online primer design tool www.LeishGEdit.net was used to design primers for amplification of the 5’ and 3’ sgRNA templates and for amplification of donor DNA from pTBlast and either pTPuro or pTNeo. Primers for deletion of *PFR2A-C* were designed with the EuPaGDT primer design tool [84] using the *PFR2* array sequence from *L. mex* Cas9 T7. For amplification of the sgRNA template DNA, primers:

Nsg: 5’-gaaattaatacgactcactataggTGCGTGCGGAGGTTTGCACGgttttagagctagaaatagc-3’ / Csg: 5’-gaaattaatacgactcactataggAAGGGTGGACGCCATCTCCGgttttagagctagaaatagc-3’.

For amplification of a pTNeo donor DNA fragment with 80 bp homology arms, primers: F: 5’GCCCACCCCTTTCACTCTTTCGCTGCTCTCTCACCTCACCGACCCCCGCCTCTTT CCATCTCTCACTGTGTGCTCCACCTgtataatgcagacctgctgc-3’ / R: 5’-AGCAGCCTTGAGACGACACCTGTAACAAAACCATCACCACAAGCTCCAAGGCGA CAACATCGCGGGAAGACTTCGCCCCAccaatttgagagacctgtgc-3’.

For transfections on 96-well plates the protocol was modified as follows (similar to descriptions in [85]): 52 × 10^7^ late log phase *L. mex* Cas9 T7 cells (1 × 10^7^ cells per reaction) were collected by centrifugation at 800*g* for 15 min. A transfection buffer was prepared by mixing 2 ml 1.5 mM CaCl_2_, 6.5 ml modified 3x Tb-BSF buffer (22.3 mM Na_2_HPO_4_, 7.67 mM NaH_2_PO_4_, 45 mM KCl, 450 mM sucrose, 75 mM HEPES pH 7.4) and 1.5 ml ddH_2_O. The cells were re-suspended in 3 ml of this transfection buffer and centrifuged again as above. The heat-sterilised sgRNA and donor DNA PCR products were placed into 48 wells of a 96-well disposable electroporation plate (4 mm gap, 250 μl, BTX) such that a given well combined all of the donor DNAs and the sgRNA templates for a given target gene. After centrifugation, cells were re-suspended in 5.2 ml transfection buffer and 100 μl of the cell suspension dispensed into each of the 48 wells containing the PCR products. Plates were sealed with foil and placed into the HT-200 plate handler of a BTX ECM 830 Electroporation System. Transfection used the following settings: 1500 V, 24 pulses, 2 counted pulses, 500 ms interval, unipolar, 100 μs. After transfection cells were recovered in 1 ml supplemented M199 in 24-well plates and incubated for 8-16 h at 28°C / 5% CO_2_ before addition of the relevant selection drugs by adding 1 ml of supplemented M199 with double the concentration of the desired drugs. Mutants were selected with 5 μg/ml Blasticidin-S deaminase in combination with either 20 μg/ml Puromycin Dihydrochloride or 40 μg/ml G-418 Disulfate and further incubated. Drug resistant populations typically emerged after one week.

### Diagnostic PCR for knockout validation

Drug-selected populations were passaged at least twice (one with at least a 1:100 dilution) before extraction of genomic DNA using the protocol described in [86]. A diagnostic PCR was done to test for the presence of the target gene ORF in putative KO lines and the parental cell line (S8 Figure). Primer3 [87] was used to design, for the entire *L. mexicana* genome (gene models based on [28]), primers to amplify a short 100 – 300 bp unique fragment of the target gene ORF (S7 Table). In a second reaction, primers 518F: 5’-CACCCTCATTGAAAGAGCAAC-3’ and 518R: 5’-CACTATCGCTTTGATCCCAGGA-3’ were used to amplify the blasticidin resistance gene from the same genomic DNA samples. For Δ*PFR2* additional primers were used to confirm deletions (S10 Figure; 1F: 5’-GCAGAAGGAGAAGAGCGAGC-3’; 1R: 5’-CCAGGAACTGCTGGTACTCC-3’; 2F: 5’-CGCAGAAGGAGAAGAGCGAG-3’; 2R: 5’-GTTGTACACGGACAGCTCCA-3’; 3F: 5’-ACCCCTTTCACTCTTTCGCTG-3’; 4R: 5’-ACCAACGACGTACACAGCAG-3’).

### Light and electron microscopy

*Leishmania* cells expressing fluorescent fusion proteins were imaged live. Samples were prepared as described in [17]. Cells were immediately imaged with a Zeiss Axioimager.Z2 microscope with a 63× numerical aperture (NA) 1.40 oil immersion objective and a Hamamatsu ORCA-Flash4.0 camera or a 63× NA 1.4 objective lens on a DM5500 B microscope (Leica Microsystems) with a Neo sCMOS camera (Andor Technology) at the ambient temperature of 25–28°C.

For transmission electron microscopy, deflagellated cell bodies and isolated flagella were prepared with a modified version of the chemical fixation protocol described in Höög et al., [88]. Pellets of cell fractions were fixed with 2.5% glutaraldehyde in 10 mM PIPES (buffer as described above) overnight at 4°C. Pellets were washed four times for 15 min in 10 mM PIPES and overlaid with 10 mM PIPES, containing 1% osmium tetraoxide and incubated at 4°C for 1 h in darkness, then washed five times with ddH_2_O for 5 min each time and stained with 500 μl of 0.5% uranyl acetate in darkness at 4°C overnight. Samples were dehydrated, embedded in resin, sectioned and on-section stained as described previously [88]. Electron micrographs were captured on a Tecnai 12 TEM (FEI) with an Ultrascan 1000 CCD camera (Gatan).

### Image processing and analysis

Micrographs were processed using Fiji [89]. To enable comparisons between the parental and tagged cell lines, the same display settings for the green fluorescence channel were used. PFR length (defined by reporter signal) and flagellar length (distance between stained kinetoplast DNA and flagellar tip) was measured with the ROI manager in Fiji [89].

### Motility assays

Mutant and parental cell lines were tracked using the previously described method in [32] with three modifications. Firstly, the scripts were modified for batch analysis of multiple files. Secondly, prior to calculation of swimming statistics any ‘drift’ due to bulk fluid flow in the sample was subtracted. As swimming direction of each cell in the population is in a random direction any drift is visible as a mean population movement between successive frames. We treated drift as an apparent translation and scaling of cell positions between successive video frames, which was then subtracted. Finally, the primary statistics we considered were mean speed (using the path at 200 ms resolution) and directionality (mean velocity as a fraction of mean speed). Swimming behaviour was measured in triplicates at approximately 6·10^6^ cells/ml and analysed from 5 μl of cell culture on a glass slide in a 250-μm deep chamber covered with a 1.5 mm cover slip using darkfield illumination with a 10× NA 0.3 objective and a Hamamatsu ORCA-Flash4.0 camera on a Zeiss Axioimager.Z2 microscope at the ambient temperature of 25–28°C. The sample was stored inverted prior to use, then turned upright immediately prior to imaging to ensure consistent motion of immotile cells through sedimentation between samples. A sample of the parental cell line killed with a final concentration of 1% formaldehyde was used as a reference for motion of completely paralysed cells through sedimentation and Brownian motion alone.

### *Lutzomyia longipalpis* infections

Sand flies were either infected with pooled mutant populations of *L. mexicana* or individually with *L. mexicana* MNYC/BZ/62/M379 wild type (WT), parental line *L. mex* Cas9 T7, knockout cell line Δ*PF16*, its add-back (*PF16*AB) [22], knockout cell line Δ*BBS2* and its add-back (*BBS2*AB). All parasites were cultivated at 23°C in M199 medium supplemented with 20% foetal calf serum (Gibco), 1% BME vitamins (Sigma), 2% sterile urine and 250 μg/ml amikin (Amikin, Bristol-Myers Squibb). Before experimental infections, logarithmic growing parasites were washed three times in PBS and resuspended in defibrinated heat-inactivated rabbit blood at a concentration of 10^6^ promastigotes/ml. *Lutzomyia longipalpis (*from Jacobina, Brazil) were maintained at 26°C and high humidity on 50% sucrose solution and a 12h light/12h dark photoperiod. Sand fly females, 3-5 days old, were fed through a chick skin membrane as described previously [90]. Fully-engorged females were separated and maintained at 26° C with free access to 50% sucrose solution. They were dissected on days 2 or 6 post blood-meal (PBM) and the guts were checked for localisation and intensity of infection by light microscopy. Parasite load was graded as described previously by [91] into four categories: zero, weak (<100 parasites/gut), moderate (100–1000 parasites/gut) and heavy (>1000 parasites/gut).

### DNA extraction for bar-seq screen

Mutant *Leishmania* lines were grown separately to late log phase and mixed in equal proportions (1·10^7^ cells per cell line). This pool was divided equally into three aliquots. DNA was extracted using the Roche High Pure Nucleic Acid Kit according to the manufacturers instructions. Each aliquot was then used to infect three separate batches of 150 sand flies. The three batches were kept separate and DNA was extracted from 50 whole sand flies two, six and nine days post blood meal, using the same kit as follows: Sand flies were placed in two 1.5 ml microcentrifuge tubes (25 flies per tube), overlaid with 100 μl tissue lysis buffer and frozen at −20°C. The dead flies were defrosted and after addition of 100 μl tissue lysis buffer and 40 μl proteinase K, flies were disrupted with a disposable plastic pestle (Bel-Art) and DNA purified according to the kit manufacturer’s instructions. DNA was eluted with 50 μl elution buffer and eluates from the same timepoint were combined, yielding 100 μl in total.

### Bar-seq library preparation and sequencing

For each sample, 600 ng isolated DNA was incubated with exonuclease VII (NEB) overnight at 37°C, purified using SPRI magnetic beads and amplified using PAGE purified custom designed p5 and p7 primers (Life Technologies), containing indexes for multiplexing and adapters for Illumina sequencing. Amplicons were again bead purified and multiplexed by pooling them in equal proportions. The final sequencing pool was once again bead purified and quantified by qPCR using NEB Library Quant Kit. Library size was determined using Agilent High Sensitivity DNA Kit on a 2100 Bioanalyzer instrument. The pool was diluted to 4 nM and spiked with 30% single indexed *Leishmania* genomic DNA, prepared using Illumina TruSeq Nano DNA Library kit according to the manufacturers instructions. The final library was spiked with 1% PhiX DNA and the Illumina sequencer was loaded with 8 pM to allow low cluster density (∼800 K/mm^2^). Sequencing was performed using MiSeq v3 150 cycles kit following the manufactures instructions with paired-end sequencing (2×75 cycles, 6 and 8 cycles index read).

MiSeq raw files were de-multiplexed using bcl2fastq (Illumina). De-multiplexed samples were subjected to barcode counting using a bash script. Each gene in the *Leishmania* genome was assigned a unique barcode – the number of times each of these barcodes appeared in the sequencing output was recorded. The criteria for barcode counting was a 100% match to the 17 nt total length. Counts for each barcode were normalized for each sample by calculating their abundance relative to all 25 barcodes. One of the mutants selected for the pooled screen was excluded from the analysis because sequencing of the cell line showed eight nucleotide mismatches in the p5 sequence (likely introduced during oligonucleotide synthesis) which precluded amplification of the barcode region. To calculate “fitness” normalized barcode counts in the pooled population before feeding were divided by normalized counts at the relevant time point post blood meal.

### Orthofinder

Orthofinder 1.1.2 was used to generate orthogroups for predicted protein coding genes from 48 eukaryotic genomes (32 ciliated species and 16 non-ciliated species): The 45 previously used by Hodges *et al*. [64] (with *Leishmania major* replaced with *Leishmania mexicana*) and supplemented with *Aspergillus nidulans, Plasmodium berghei* and *Schistosoma mansoni*.

## Supporting information

## Acknowledgements

We like to thank Svenja Hester, Philip Charles and Benjamin Thomas Central Proteomics Facility at the Sir William Dunn School of Pathology for help with protein mass spectrometry, Errin Johnson (Dunn School Bioimaging Facility) for assistance with electron microscopy, Amanda Williams (University of Oxford) for help with Illumina sequencing, Maxime Cesca (Université de Paris Sud) and Laura Makin (University of Oxford) for assistance with mutant characterisation, Oliver Billker (Wellcome Sanger Institute) and members of O.B.’s lab for helpful discussions about the bar-seq method, Sébastien Pomel (Université de Paris Sud) for drawing our attention to the PMI, PMM and GDP-MP mutants, Diane McMahon-Pratt (Yale University) for antibody 2E10 (F-4) and Keith Gull (University of Oxford) for advice and helpful comments on the manuscript, antibodies L8C4 and L13D6 and access to equipment, and members of K.G.’s lab for helpful discussions, particularly Samuel Dean and Jack Sunter for help with development of high-throughput transfection protocols.

