## Supplementary material for "Genetic dissection of a *Leishmania* flagellar proteome demonstrates requirement for directional motility in sand fly infections": S6_Table_mass_spectrometry_search_parameters_Beneke_et_al.docx

| Action | **Filter type** | **value** |
| --- | --- | --- |
| Proteo  Wizard | Threshold | Count 200 most-intense |
|  | Peak picking | True 1- |
| Mascot/X!Tandem kscore/OMSSA | Missed cleavages | 2 |
|  | Precursor Tolerance | 0.1 Da |
|  | Charge State(s) | 1+, 2+ and 3+ |
|  | Fixed Modification | Carbamidomethyl (C) |
|  | Variable Modifications | Deamidated (N), Deamidated (Q), Oxidation (M) |
|  | Quantitation | None |
|  | Quantitation Tolerance | 0.02 |
| CPFP | PSM & Peptide-level FDR | 1.00% |
|  | PSM & Peptide-level q-value | 0.01 |
|  | Protein group FDR | 1.00% |
|  | Protein group q-value | 0.01 |
| SINQ | PROT_MAX_QVAL | 0.01 |
|  | PSM_MAX_QVAL | 0.01 |
|  | FRAGMENT_TOL | 0.1 |
|  | Maximum q-value (local FDR) | 0.01 |
|  | **Peptides per protein** | **at least 2** |

S6 Table. Mass spectrometry search parameters
