## Supplementary material for "Genetic dissection of a *Leishmania* flagellar proteome demonstrates requirement for directional motility in sand fly infections"

Beneke *et al.*

### Supplementary Tables

#### S1 Table SINQ meta analysis, raw data

Table shows the raw output from the CPFP pipeline with label-free SINQ quantification. The data table contains all identified proteins (listed by GeneIDs) from Run 1 and 2 and their relative enrichment across all four fractions. Listed is the spectral count (not normalized enrichment) and spectral index (normalized enrichment).

#### S2 Table GeneIDs Run 1 (p > 0.95, number of peptides > 2)

GeneIDs identified in Run 1 (S1 Table) were filtered (p > 0.95, number of peptides > 2) and listed with additional information. Tab "all proteins Run 1" lists all GeneIDs, Tabs "$C_l$ square Run 1", "$C_S$ square Run 1", "$F_l$ square Run 1" and "$F_S$ square Run 1" list GeneIDs enriched in the respective fractions.

#### S3 Table GeneIDs Run 2 (p > 0.95, number of peptides > 2)

GeneIDs identified in Run 2 (S1 Table) were filtered (p > 0.95, number of peptides > 2) and listed as described for Table S2.

#### S4 Table mutant summary of analysed mutants

#### S5 Table Orthofinder results

Tab Orthofinder Results: Summary of orthologs identified in the genomes of 33 ciliated and 15 non-ciliated species. *Leishmania mexicana* genes analysed in this study are printed in bold. The numbers indicate number of orthologs in each species. Sensitivity is the observed probability of a flagellated organism having at least one member of that orthogroup. Specificity is the observed probability of a non-flagellated organism lacking any members of that orthogroup. A high sensitivity and specificity indicates an orthogroup is a good predictor of a species' ability to build a flagellum and high specificity indicates a low type I (false positive) error rate. Tab Genome Versions: A list of genome versions used for the analysis. Tab Genes Selected Species: Gene accession numbers for orthologs identified in the genome of *Leishmania mexicana*, *Trypanosoma brucei*, *Chlamydomonas reinhardtii* and *Homo sapiens* genomes. Tab Genes All Species: gene accession numbers for all analysed genomes.

#### S6 Table Mass spectrometry search parameters

#### S7 Table Knockout verification primers

List of primer sequences for validation of gene deletion by PCR for all annotated *L. mexicana* genes and novel genes defined by Fiebig et al. [28]. Sequences are written in 5' to 3' orientation and the product length is indicated.

#### S8 Table PFR measurements

#### S9 Table Sand fly reads

### Supplementary Figures

#### S1 Figure. Quantitation of yield and purity

Counts of whole flagellated cells, isolated flagella or deflagellated cell bodies for each stage of the deflagellation procedure. The purified cell body fraction contains 2.03% (±0.69%) isolated flagella; the purified flagella fraction contains 0.56% (±0.15%) deflagellated cell bodies. Error bars represent standard deviations between four biological replicates.

#### S2 Figure. Transmission electron micrographs of isolated cell components

**(A, B)** Sections from the isolated flagella fraction and **(C, D)** from the deflagellated cell body fraction. Arrows indicate **(A)** flagellar tip structure, **(B)** cross-section through flagellum with intact axoneme, associated PFR and surrounding membrane, **(C)** anterior end of a deflagellated cell body and **(D)** intact axonemal structure of flagellum inside the flagellar pocket. Scale bars represent 1 µm.

#### S3 Figure. Length measurements on isolated flagella

**(A)** Measurements of flagellar lengths for intact cells, isolated flagella or deflagellated cell bodies from four independent samples. **(B)** Average lengths derived from data shown in (A). The length measurements of the cell body fractions and flagella fractions are stacked to show the combined total length after deflagellation. Combined standard deviation for isolated flagella and deflagellated cell bodies is calculated by $z = \sqrt{x^2 + y^2}$. **(C)** Electron micrograph of an intact *L. mexicana* promastigote cell. The red line shows the average length of flagellum remaining attached to the cell body.

#### S4 Figure. Pearson's and Spearman's correlation between biological replicates

**(A)** Pearson's correlation coefficient and **(B)** Spearman's rank of spectral indices for 1918 detected proteins (overlap of both biological replicates) of all cell fractions.

#### S5 Figure. Comparison with published proteomes

All 2414 proteins detected in run 1 were plotted as in Figure 2A. Highlighted in colour are **(A)** orthologs of *L. donovani* flagellar proteins [92], **(B)** orthologs of detergent insoluble salt-extracted *T. brucei* flagellar proteins [6], **(C)** orthologs of *T. brucei* flagellar surface and matrix proteins [78], **(D)** orthologs of *T. brucei* proteins detected in mechanically sheared flagella [93], **(E)** *L. mexicana* ribosomal proteins and **(F)** *L. mexicana* proteins with trans-membrane domain predictions. Each plot can be interactively explored on http://www.leishgedit.net/leishgedit_db/.

#### S6 Figure. Molecular weight and isoelectric point distributions of proteins.

Molecular weight and isoelectric point calculated with Isoelectric Point Calculator [94]. The isoelectric point prediction model from EMBOSS is shown. **(A)** All annotated proteins, based on gene models from [28]. **(B)** Proteins detected in MS run 1. Two sample Kolmogorov–Smirnov test shows significant difference in distributions between (A) and (B) (p-value = $9.63e^{-30}$).

#### S7 Figure. Localisation of tagged proteins

Widefield epifluorescence micrographs of *L. mexicana* cells. Left image: merged phase-contrast (grey), red (Hoechst-stained DNA), and green (mNG or eYFP) channels. Right image: greyscale image of green channel (mNG or eYFP signal). For each tagged protein, the relevant parental cell line (wild type for proteins tagged with eYFP, *L. mex* Cas9 T7 for proteins tagged with mNG) was imaged, using the same acquisition settings and image processing parameters as for the tagged cell line. For each protein, panels on the left show N-terminal tags, panel on the right C-terminal tags.

#### S8 Figure. PCR validation of KO cell lines

Cartoons showing PCR strategy: **(A)** amplification of a fragment of the target gene ORF, **(B)** amplification of a fragment of the inserted blasticidin resistance gene (BlastR). **(C)** PCR products from target gene ORF run on agarose gel. Each panel shows the product obtained from the putative knockout cell line (K) and the parental *L. mex* Cas9 T7 cell line (P). The

GeneIDs / gene names indicate the target gene. Absence of the target ORF PCR product was confirmed for all genes, except for LmxM.34.4010, CALP1.1, LmxM.09.0910 and CMF6. See S7 Table for amplicon sizes. Fainter bands below the target gene amplicon are likely primer dimers. (**D**) Presence of the BlastR PCR product was confirmed for all transfected cell lines; no BlastR PCR product was amplified from the parental genome (P).

### S9 Figure. Quantitation of curled flagella

Histogram showing the proportion of cells with curled flagella in the parental *L. mex* Cas9 T7 cell line, 20 different KO mutants and three add-back cell lines (AB). The GeneIDs / gene names indicate the target gene. Numbers above the bars indicate percentage of cells with curly flagella.

### S10 Figure. Deletion of *PFR2* array

(**A**) Strategy for deletion of *PFR2* array, using CRISPR-Cas9 to insert a pTNeo cassette with homology arms flanking the array. (**B**) Cartoon showing location of five primer pairs used to validate KO cell line. (**C**) Results of diagnostic PCR. Primer pairs 1, 2, 3 and 4 specific to PFR2 array sequences only yield a product in the parental cell line (+/+) but not in the KO line (-/-). Primer pair 5, specific to an unrelated gene (PMM, LmxM.36.1960) yields a product in the parental and the KO cell line. (**A**), (**B**) and (**C**) Size of deleted PFR2 array and expected PCR amplicons are indicated. (**D**) Western blots of parental and PFR2 KO protein lysates. Upper panel: blots probed with three different monoclonal antibodies: L8C4 [95] (1:1000 dilution in TBST 1% skim milk powder, sigma) is specific to PFR2, L13D6 [95] (1:20 dilution in TBST 1% skim milk powder) detects PFR1 and 2E10 detects both PFR1 and PFR2 [96] (1:1000 dilution in TBST 5% skim milk powder). No PFR2 signal is detected in the PFR2 KO line. Lower panel: Ponceau red stained membrane.

### S11 Figure. Strategy for pooled sand fly infection

**(A)** Barcoded cell lines in log phase of growth were pooled in equal proportion. **(B)** For each replicate, 150 sand flies were fed on rabbit blood containing the pooled *Leishmania*. **(C)** DNA was isolated from the pooled population before feeding the sand flies. Two, six and nine days post blood meal, DNA was isolated from 50 infected flies. **(D, E)** The 17nt unique barcode and adjacent constant region was amplified from the isolated DNA. Primer binding sites for indexed p5 and p7 Illumina sequencing primers are indicated in red. **(F)** Final amplicon library constructs subjected to Illumina sequencing. Samples were multiplexed using different p5 and p7 indices.

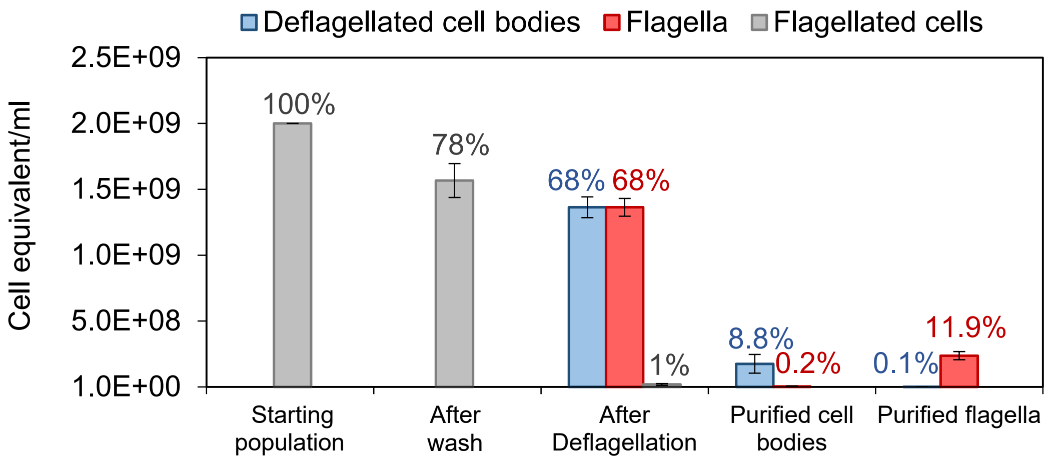
Deflagellated cell bodies
Flagella
Flagellated cells
Cell equivalent/ml
2.5E+09
2.0E+09
1.5E+09
1.0E+09
5.0E+08
1.0E+00
100%
78%
68%
68%
1%
8.8%
0.2%
0.1%
11.9%
Starting population
After wash
After Deflagellation
Purified cell bodies
Purified flagella

Figure S1

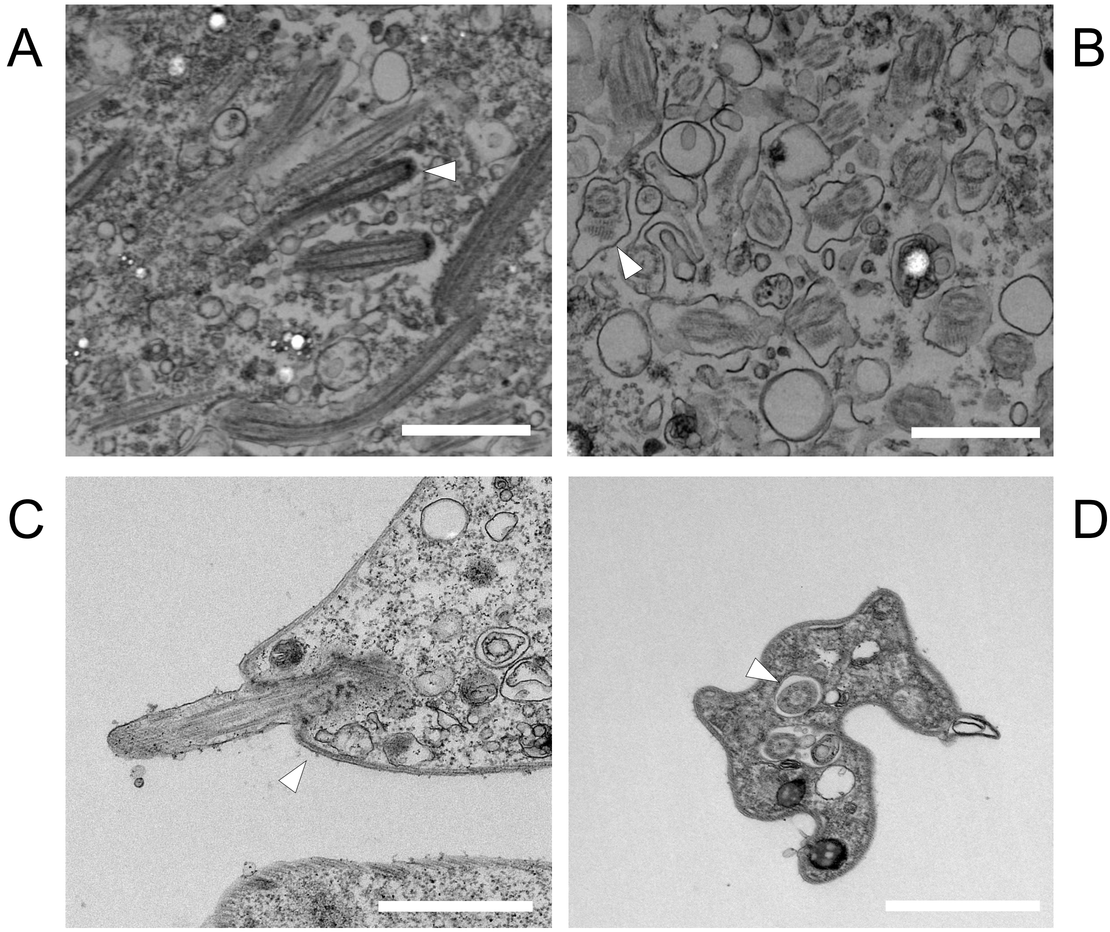

Figure S2

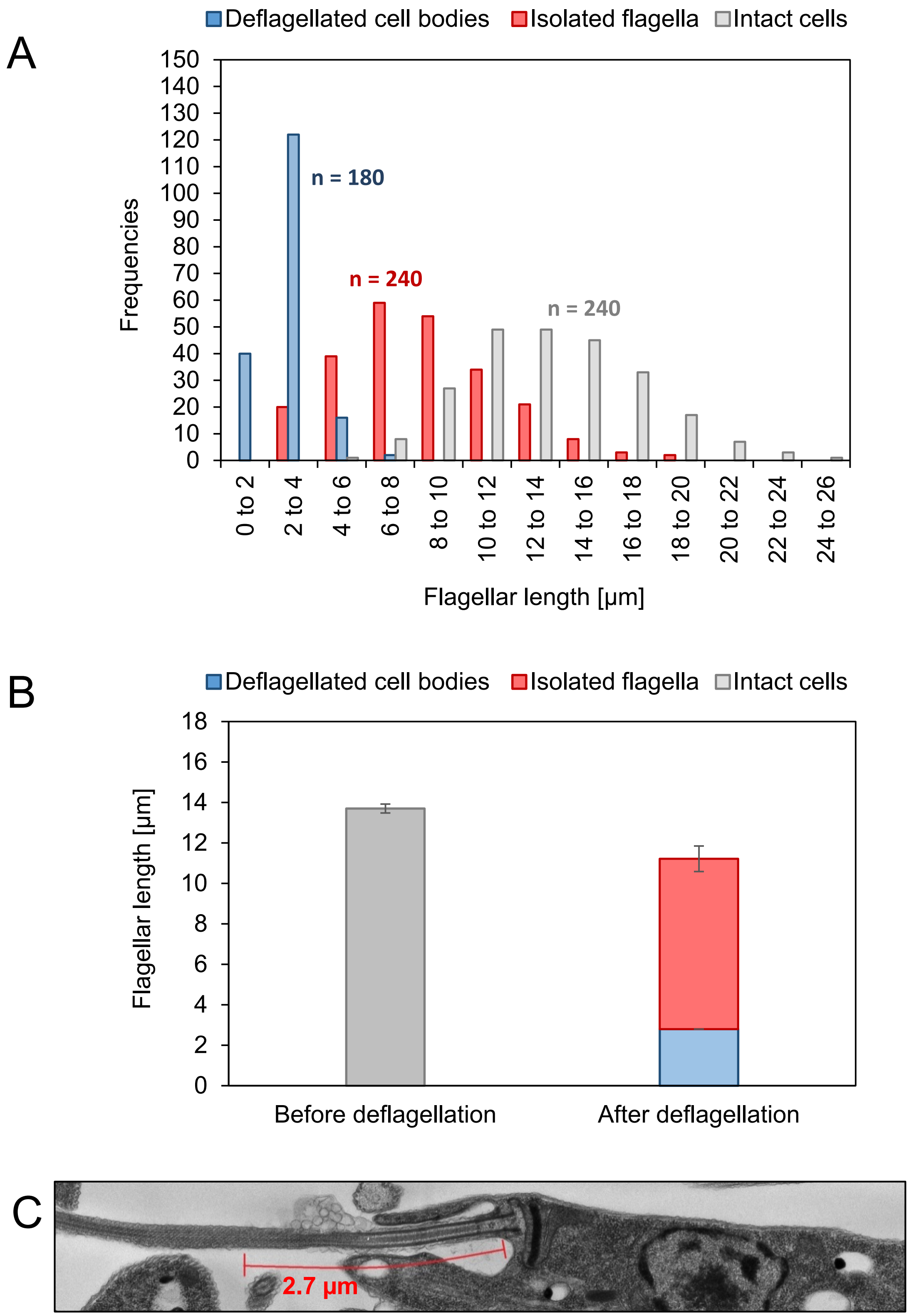
A
Deflagellated cell bodies
Isolated flagella
Intact cells
n = 180
n = 240
n = 240
Frequencies
0
10
20
30
40
50
60
70
80
90
100
110
120
130
140
150
0 to 2
2 to 4
4 to 6
6 to 8
8 to 10
10 to 12
12 to 14
14 to 16
16 to 18
18 to 20
20 to 22
22 to 24
24 to 26
Flagellar length [µm]
B
Deflagellated cell bodies
Isolated flagella
Intact cells
Flagellar length [µm]
0
2
4
6
8
10
12
14
16
18
Before deflagellation
After deflagellation
C
2.7 µm

Figure S3

A

**Pearson correlation coefficient**

| | $C_I$ 2 | $C_I$ 1 | $C_S$ 2 | $C_S$ 1 | $F_I$ 2 | $F_I$ 1 | $F_S$ 2 | $F_S$ 1 |
|---|---|---|---|---|---|---|---|---|
| **$C_I$ 2** | 1.00 | 0.72 | 0.28 | 0.27 | 0.68 | 0.49 | 0.16 | 0.21 |
| **$C_I$ 1** | 0.72 | 1.00 | 0.53 | 0.62 | 0.53 | 0.56 | 0.28 | 0.38 |
| **$C_S$ 2** | 0.28 | 0.53 | 1.00 | 0.78 | 0.58 | 0.44 | 0.58 | 0.61 |
| **$C_S$ 1** | 0.27 | 0.62 | 0.78 | 1.00 | 0.41 | 0.45 | 0.50 | 0.60 |
| **$F_I$ 2** | 0.68 | 0.53 | 0.58 | 0.41 | 1.00 | 0.73 | 0.65 | 0.57 |
| **$F_I$ 1** | 0.49 | 0.56 | 0.44 | 0.45 | 0.73 | 1.00 | 0.40 | 0.50 |
| **$F_S$ 2** | 0.16 | 0.28 | 0.58 | 0.50 | 0.65 | 0.40 | 1.00 | 0.81 |
| **$F_S$ 1** | 0.21 | 0.38 | 0.61 | 0.60 | 0.57 | 0.50 | 0.81 | 1.00 |

B

**Spearman's rank**

| | $C_I$ 2 | $C_I$ 1 | $C_S$ 2 | $C_S$ 1 | $F_I$ 2 | $F_I$ 1 | $F_S$ 2 | $F_S$ 1 |
|---|---|---|---|---|---|---|---|---|
| **$C_I$ 2** | 1.00 | 0.86 | 0.62 | 0.58 | 0.51 | 0.46 | 0.49 | 0.54 |
| **$C_I$ 1** | 0.86 | 1.00 | 0.57 | 0.61 | 0.47 | 0.48 | 0.44 | 0.55 |
| **$C_S$ 2** | 0.62 | 0.57 | 1.00 | 0.88 | 0.36 | 0.32 | 0.72 | 0.74 |
| **$C_S$ 1** | 0.58 | 0.61 | 0.88 | 1.00 | 0.29 | 0.33 | 0.67 | 0.75 |
| **$F_I$ 2** | 0.51 | 0.47 | 0.36 | 0.29 | 1.00 | 0.84 | 0.47 | 0.42 |
| **$F_I$ 1** | 0.46 | 0.48 | 0.32 | 0.33 | 0.84 | 1.00 | 0.43 | 0.49 |
| **$F_S$ 2** | 0.49 | 0.44 | 0.72 | 0.67 | 0.47 | 0.43 | 1.00 | 0.83 |
| **$F_S$ 1** | 0.54 | 0.55 | 0.74 | 0.75 | 0.42 | 0.49 | 0.83 | 1.00 |

Figure S4

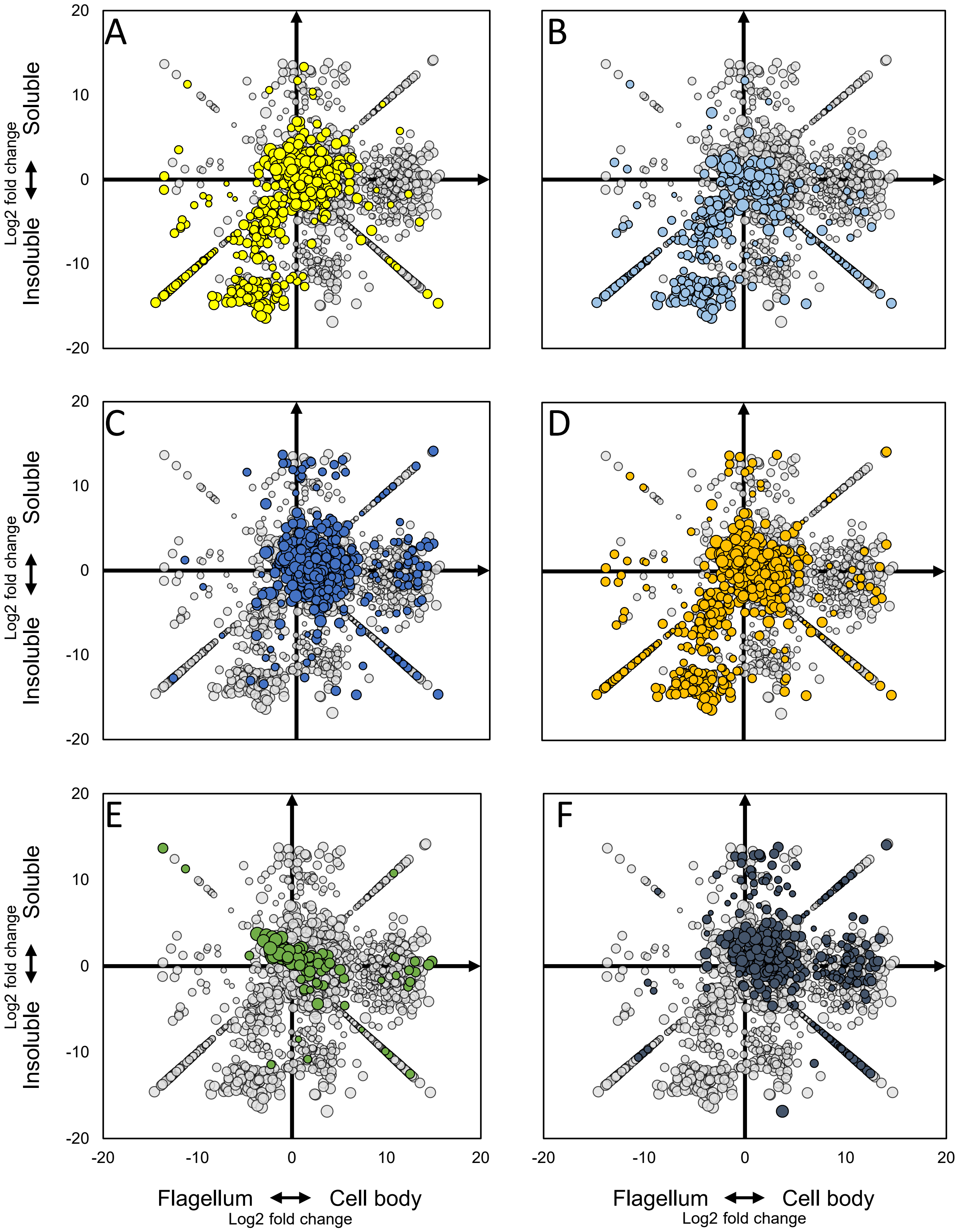
A
B
C
D
E
F
Soluble
Insoluble
Log2 fold change
Flagellum
Cell body
Log2 fold change
20
10
0
-10
-20
-20
-10
0
10
20

Figure S5

A

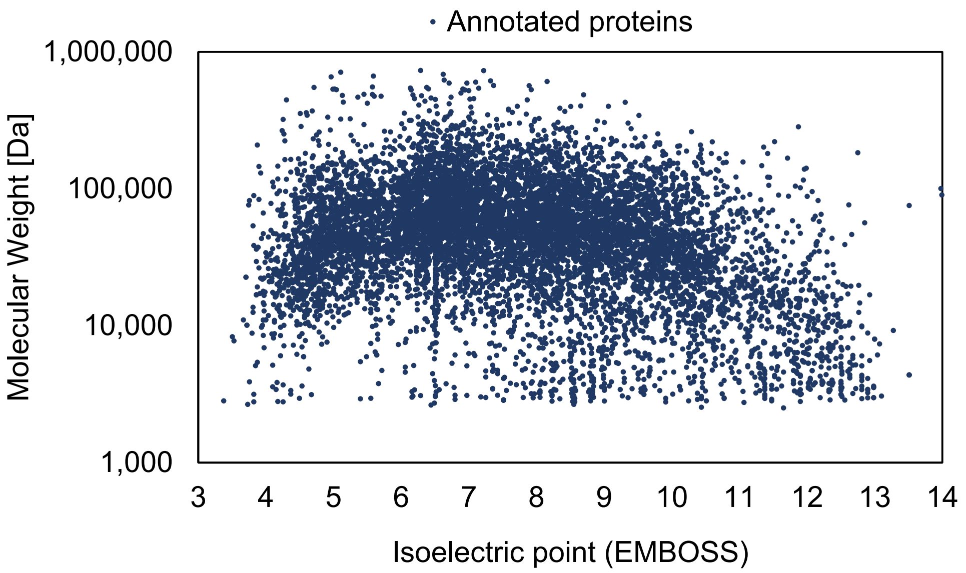

Annotated proteins
1,000,000
100,000
10,000
1,000
Molecular Weight [Da]
Isoelectric point (EMBOSS)

B

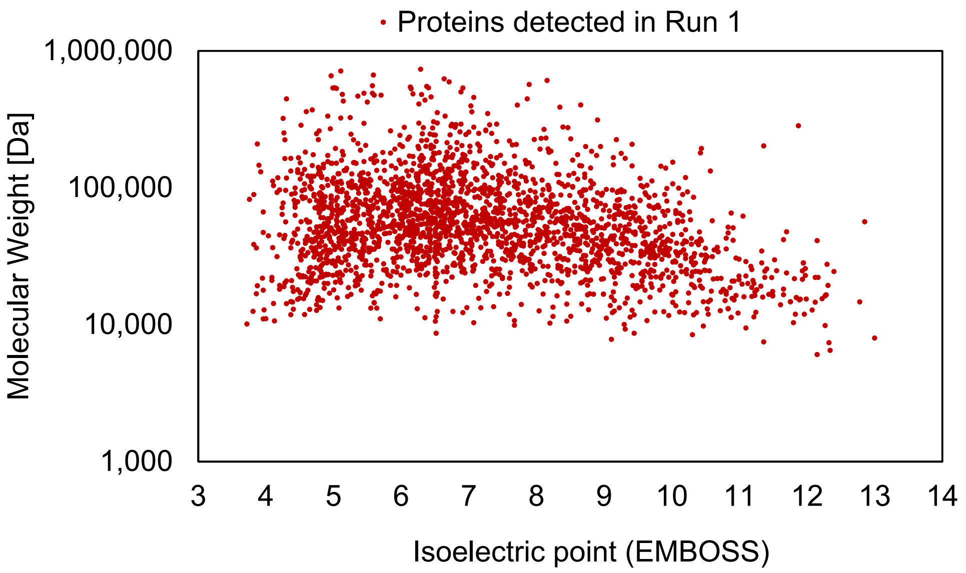

Proteins detected in Run 1
1,000,000
100,000
10,000
1,000
Molecular Weight [Da]
Isoelectric point (EMBOSS)

Figure S6

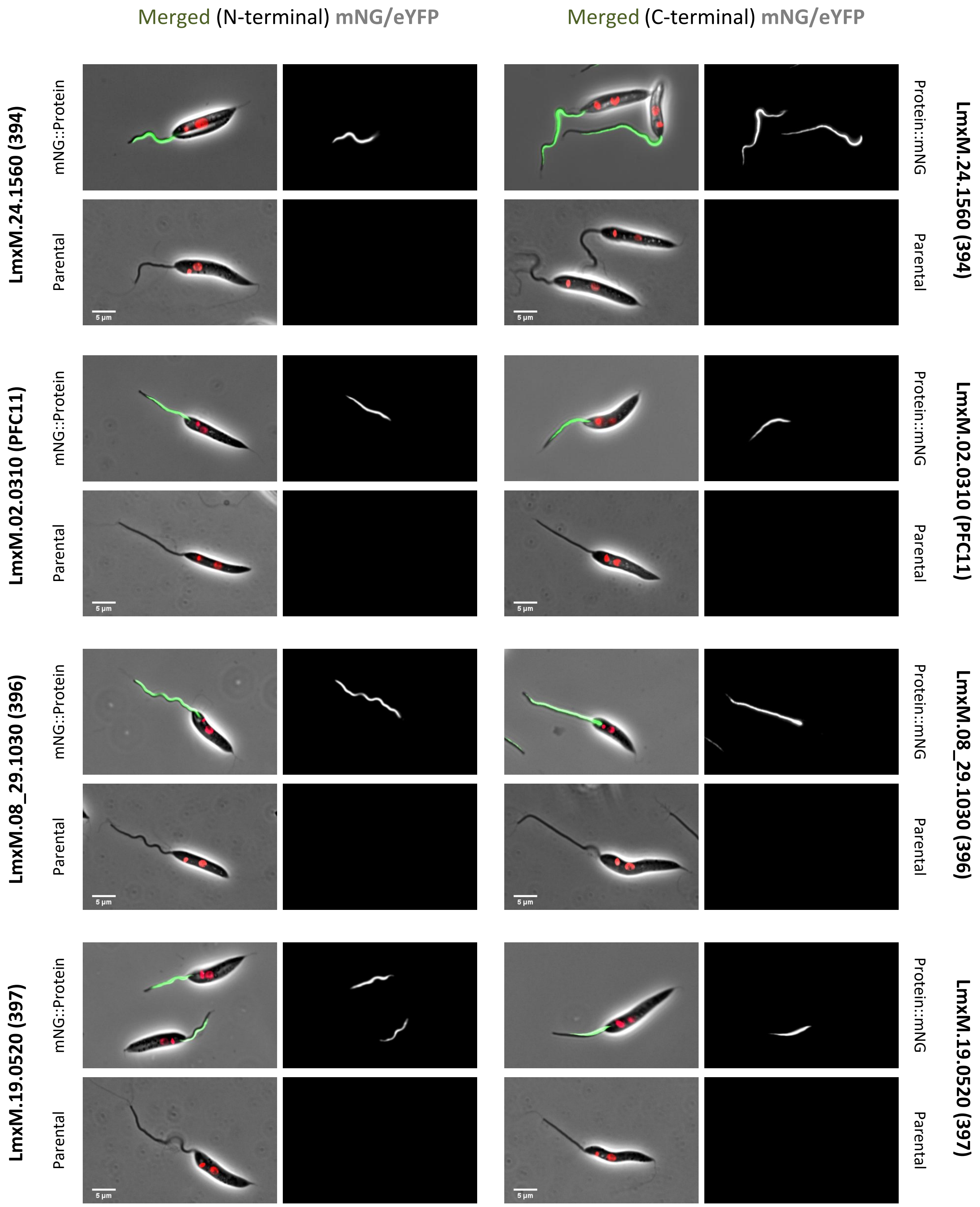
Merged (N-terminal) mNG/eYFP
Merged (C-terminal) mNG/eYFP
LmxM.24.1560 (394)
LmxM.02.0310 (PFC11)
LmxM.08_29.1030 (396)
LmxM.19.0520 (397)
mNG::Protein
Parental
Protein::mNG
5 µm

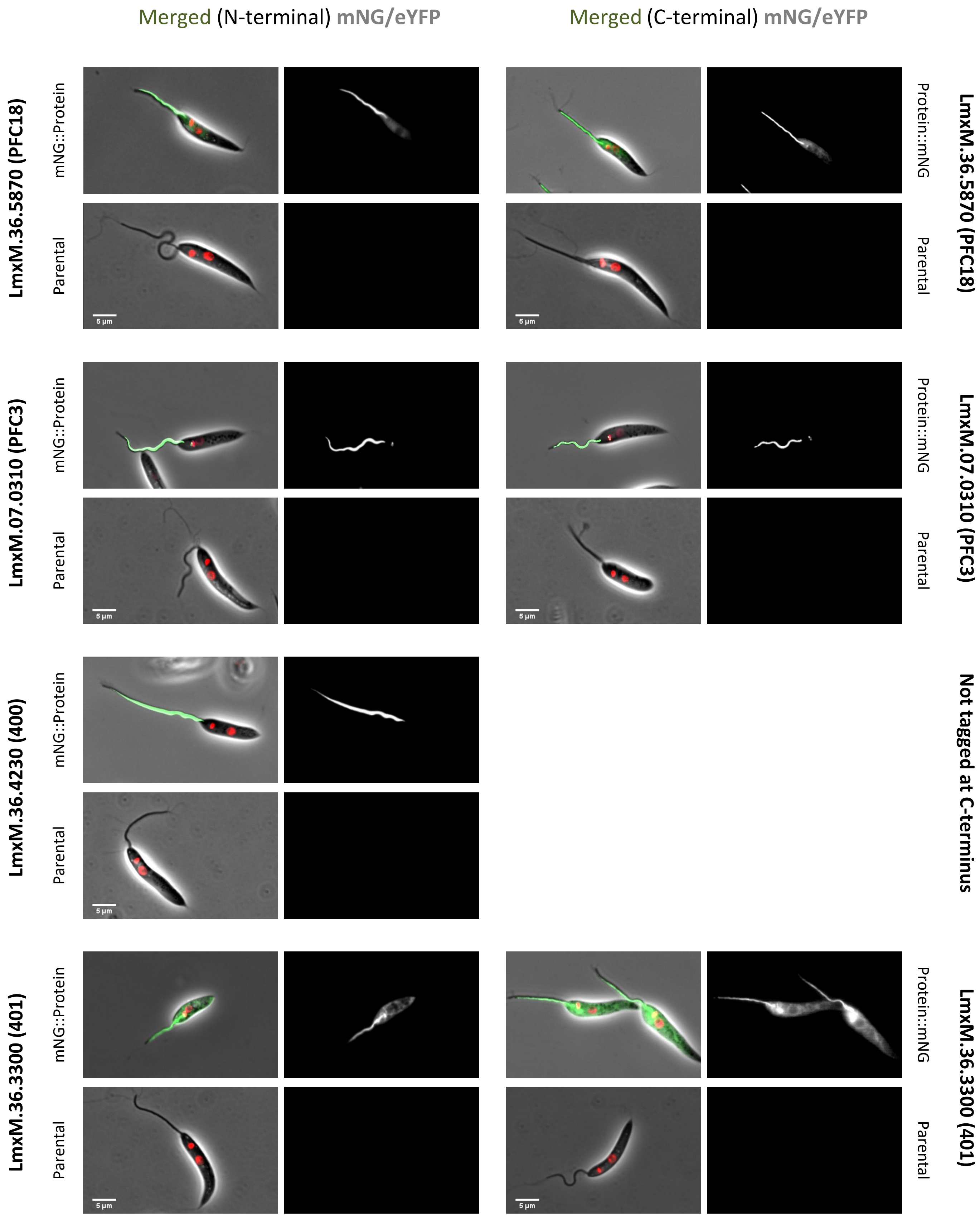

Merged (N-terminal) mNG/eYFP
Merged (C-terminal) mNG/eYFP
LmxM.36.5870 (PFC18)
mNG::Protein
Parental
Protein::mNG
Parental
LmxM.36.5870 (PFC18)
LmxM.07.0310 (PFC3)
mNG::Protein
Parental
Protein::mNG
Parental
LmxM.07.0310 (PFC3)
LmxM.36.4230 (400)
mNG::Protein
Parental
Not tagged at C-terminus
LmxM.36.3300 (401)
mNG::Protein
Parental
Protein::mNG
Parental
LmxM.36.3300 (401)
5 µm

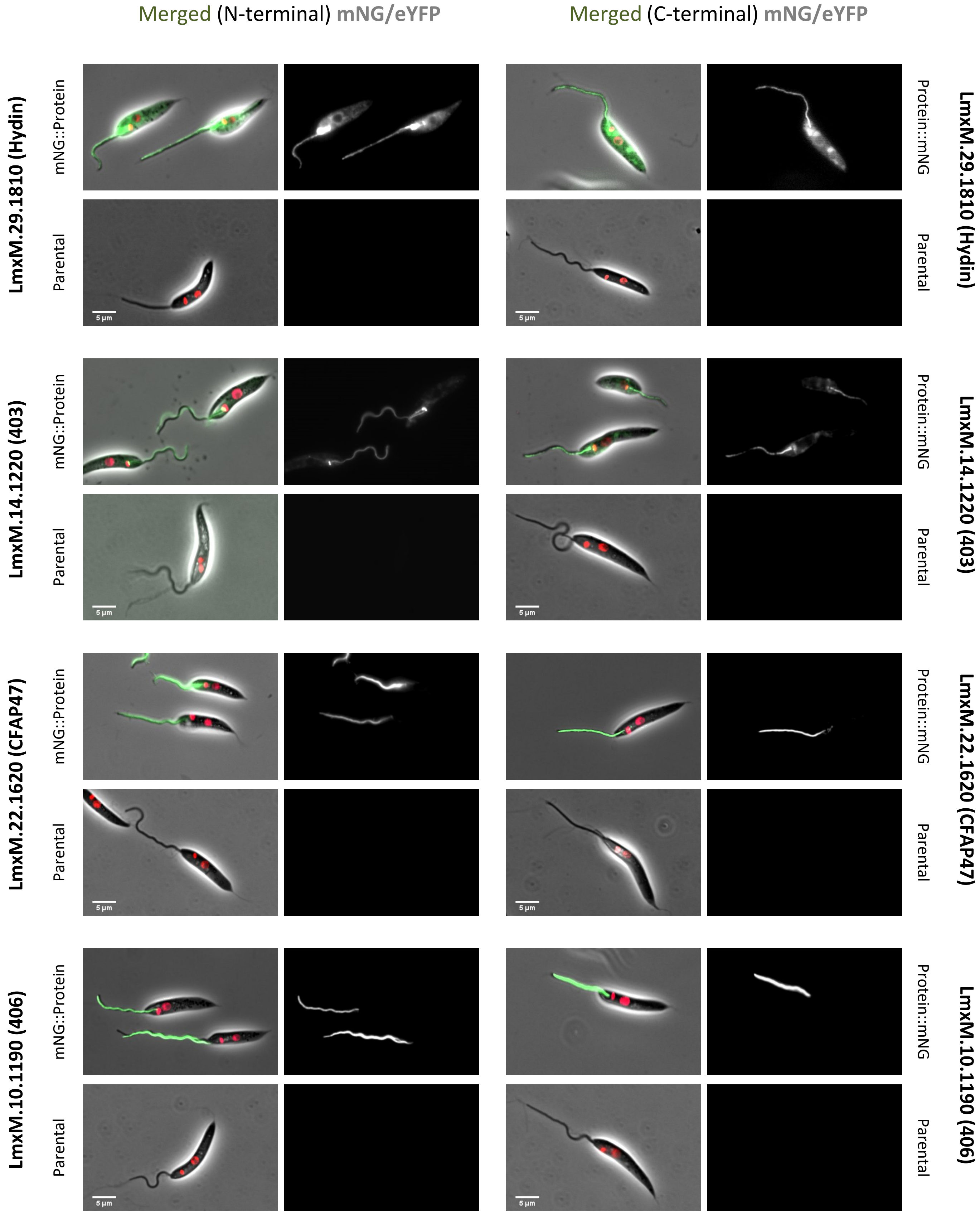

Merged (N-terminal) mNG/eYFP
Merged (C-terminal) mNG/eYFP
LmxM.29.1810 (Hydin)
LmxM.14.1220 (403)
LmxM.22.1620 (CFAP47)
LmxM.10.1190 (406)
mNG::Protein
Protein::mNG
Parental
5 µm

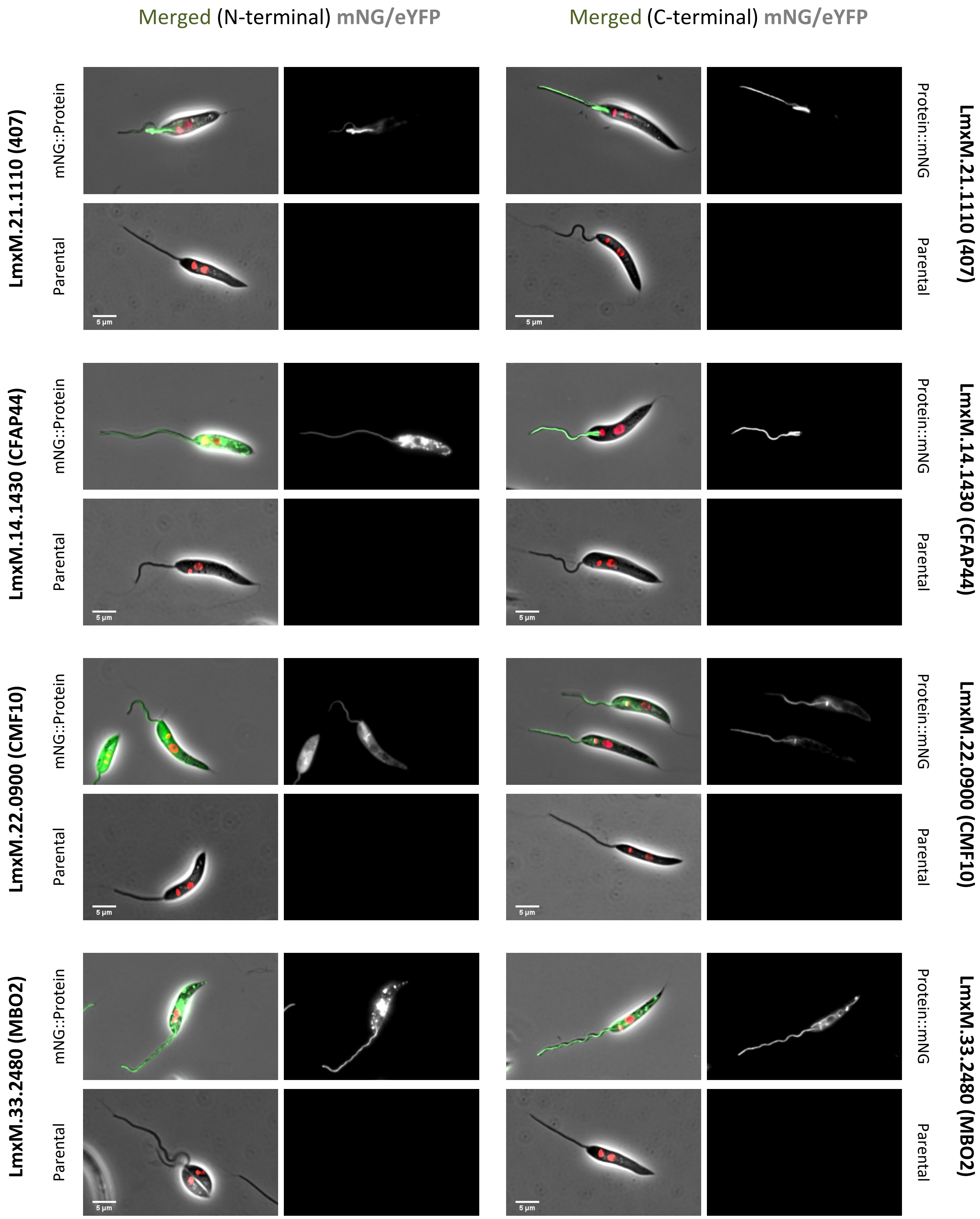

Merged (N-terminal) mNG/eYFP
Merged (C-terminal) mNG/eYFP
LmxM.21.1110 (407)
LmxM.14.1430 (CFAP44)
LmxM.22.0900 (CMF10)
LmxM.33.2480 (MBO2)
mNG::Protein
Protein::mNG
Parental
5 µm

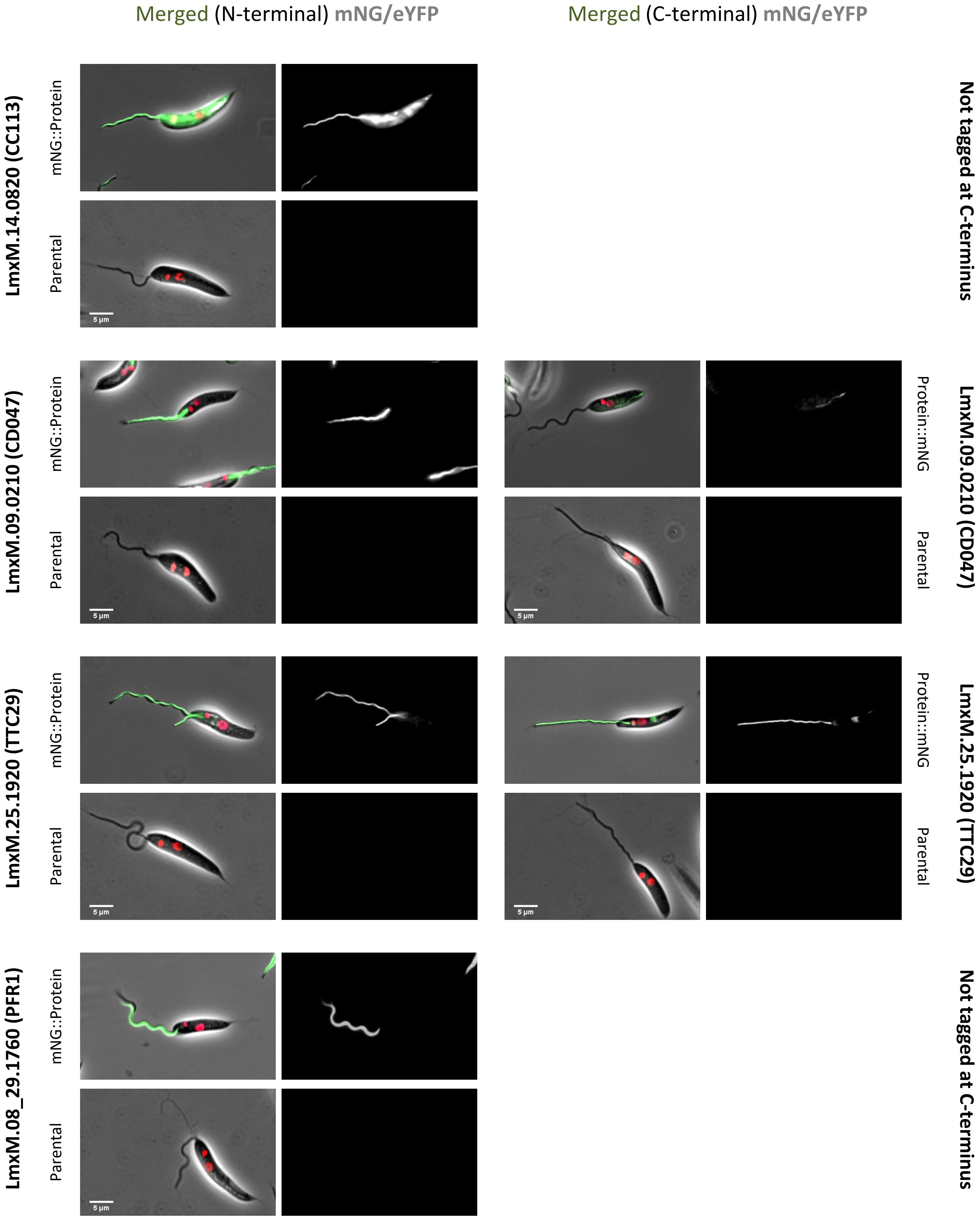
Merged (N-terminal) mNG/eYFP
Merged (C-terminal) mNG/eYFP
LmxM.14.0820 (CC113)
mNG::Protein
Parental
5 µm
Not tagged at C-terminus
LmxM.09.0210 (CD047)
mNG::Protein
Parental
5 µm
Protein::mNG
Parental
5 µm
LmxM.09.0210 (CD047)
LmxM.25.1920 (TTC29)
mNG::Protein
Parental
5 µm
Protein::mNG
Parental
5 µm
LmxM.25.1920 (TTC29)
LmxM.08_29.1760 (PFR1)
mNG::Protein
Parental
5 µm
Not tagged at C-terminus

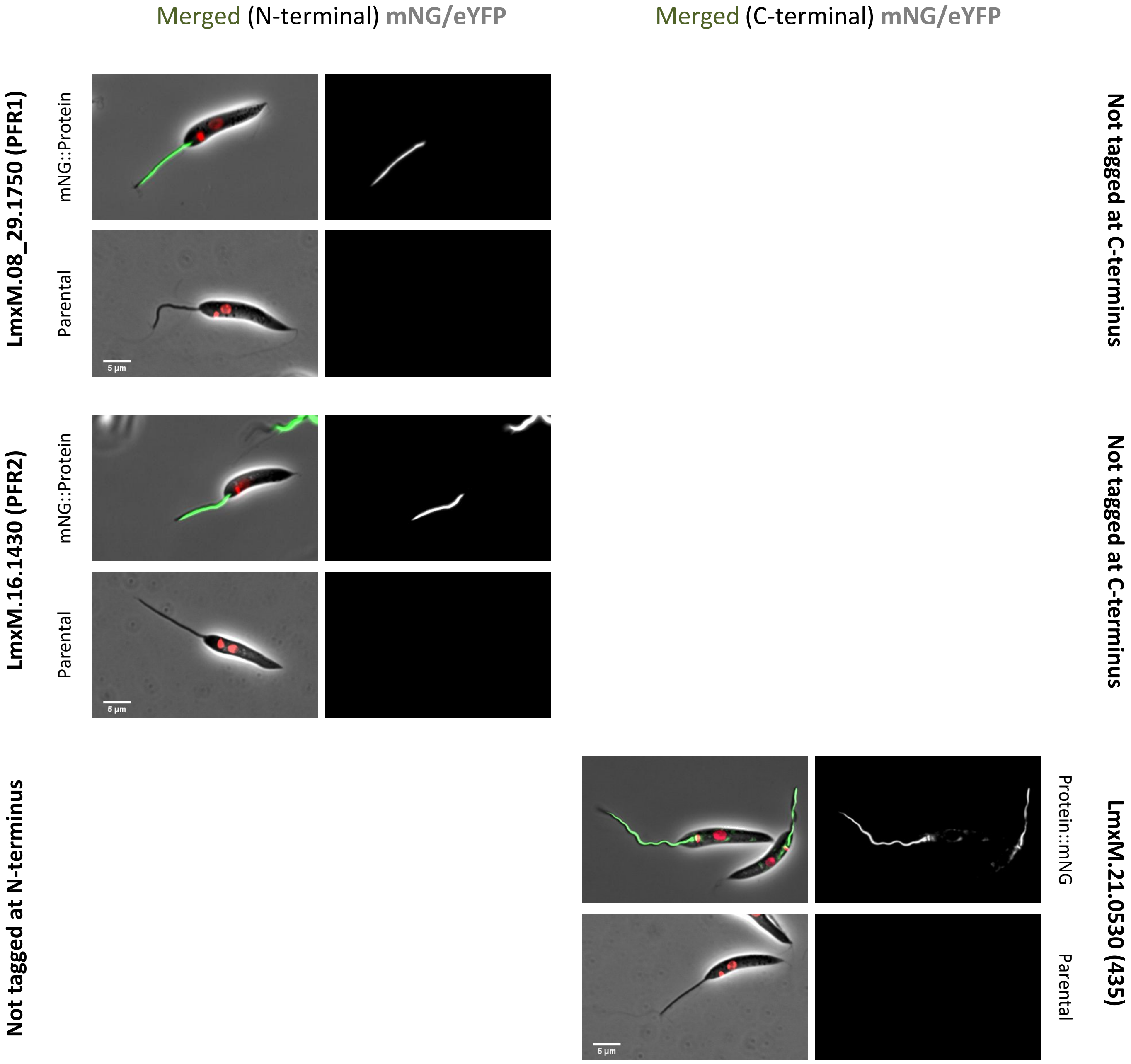
Merged (N-terminal) mNG/eYFP
Merged (C-terminal) mNG/eYFP
LmxM.08_29.1750 (PFR1)
mNG::Protein
Parental
5 µm
Not tagged at C-terminus
LmxM.16.1430 (PFR2)
mNG::Protein
Parental
5 µm
Not tagged at C-terminus
Not tagged at N-terminus
LmxM.21.0530 (435)
Protein::mNG
Parental
5 µm

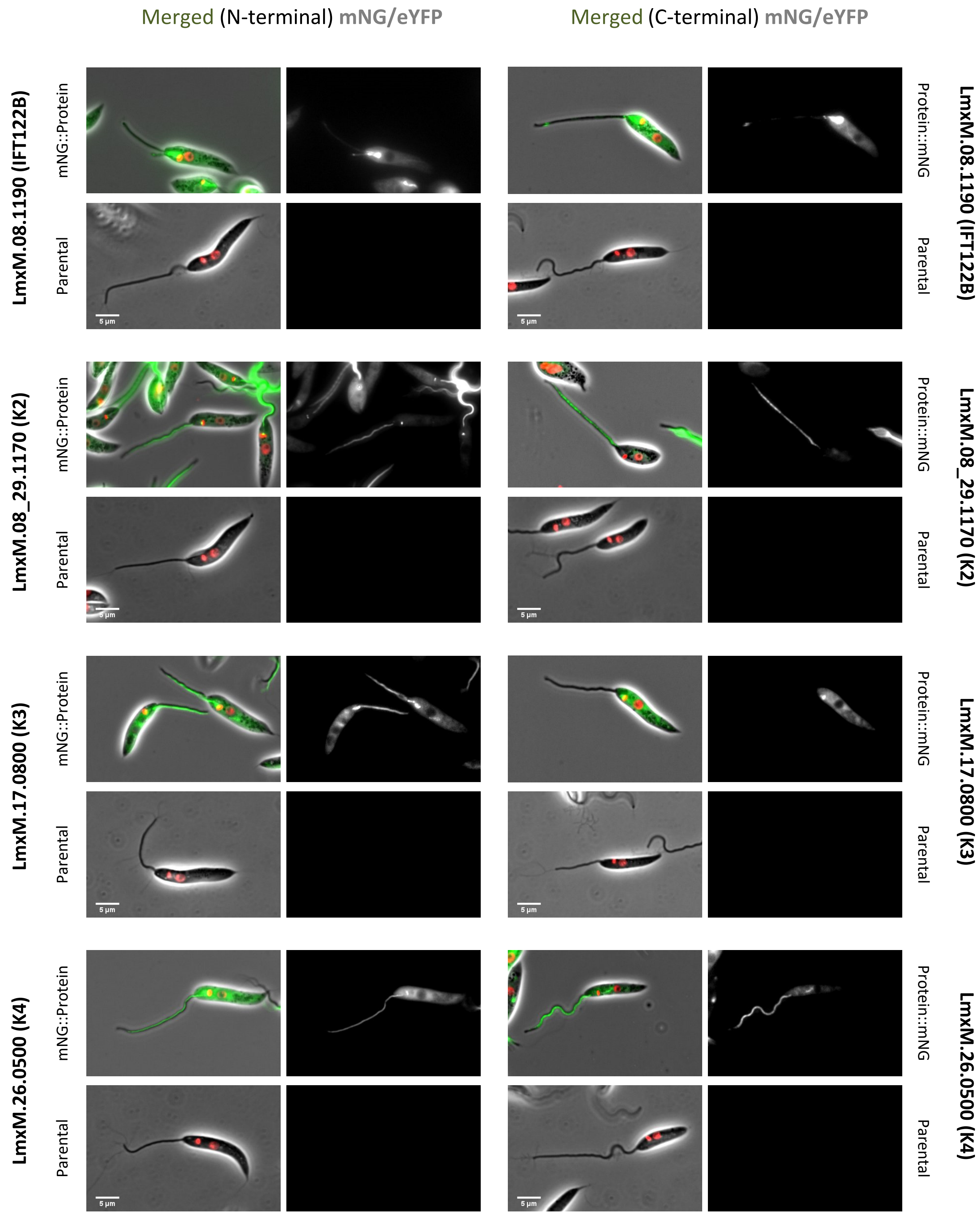

Merged (N-terminal) mNG/eYFP
Merged (C-terminal) mNG/eYFP
LmxM.08.1190 (IFT122B)
mNG::Protein
Parental
Protein::mNG
LmxM.08_29.1170 (K2)
LmxM.17.0800 (K3)
LmxM.26.0500 (K4)
5 µm

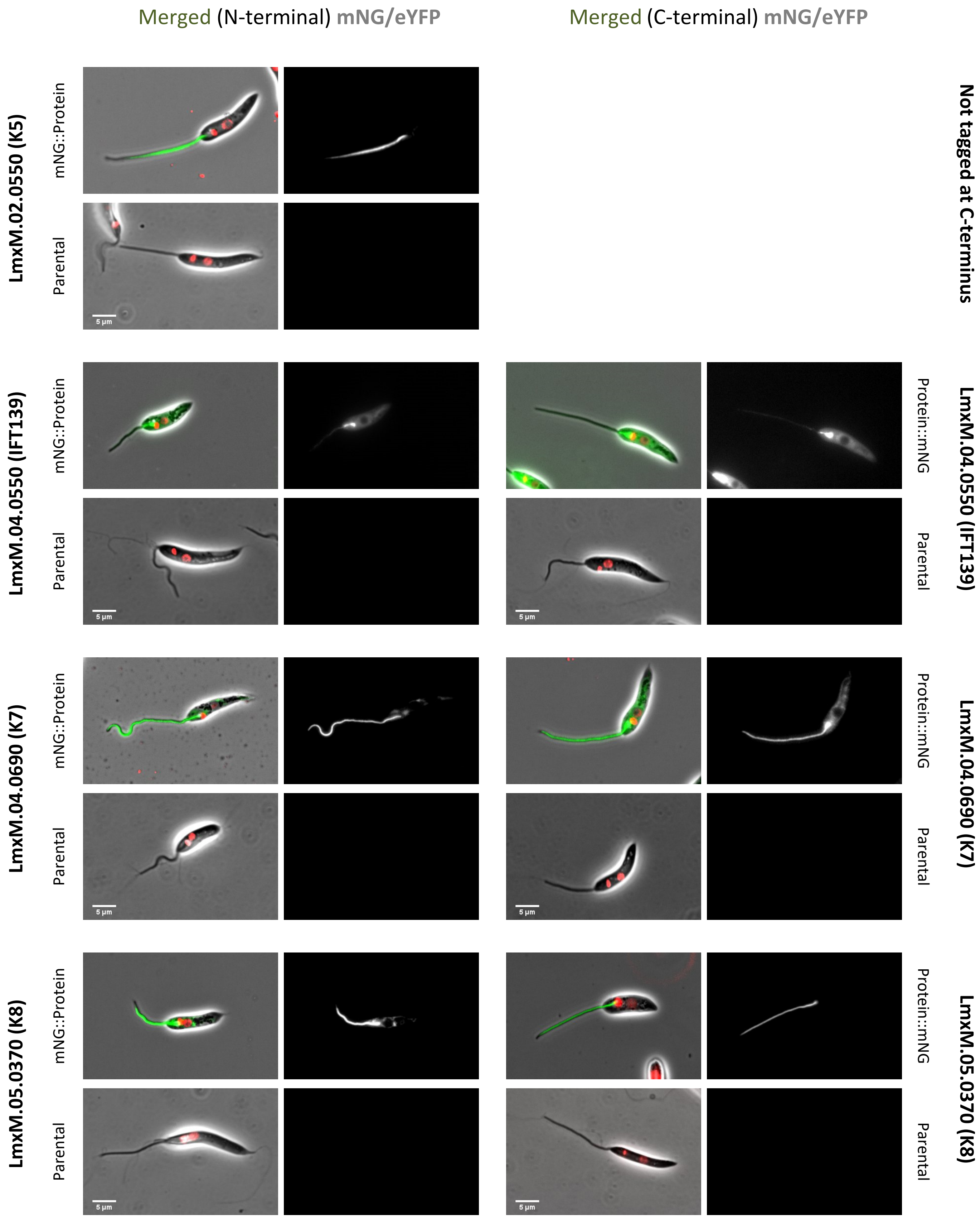

Merged (N-terminal) mNG/eYFP
Merged (C-terminal) mNG/eYFP
LmxM.02.0550 (K5)
mNG::Protein
Parental
Not tagged at C-terminus
LmxM.04.0550 (IFT139)
Protein::mNG
LmxM.04.0690 (K7)
LmxM.05.0370 (K8)
5 µm

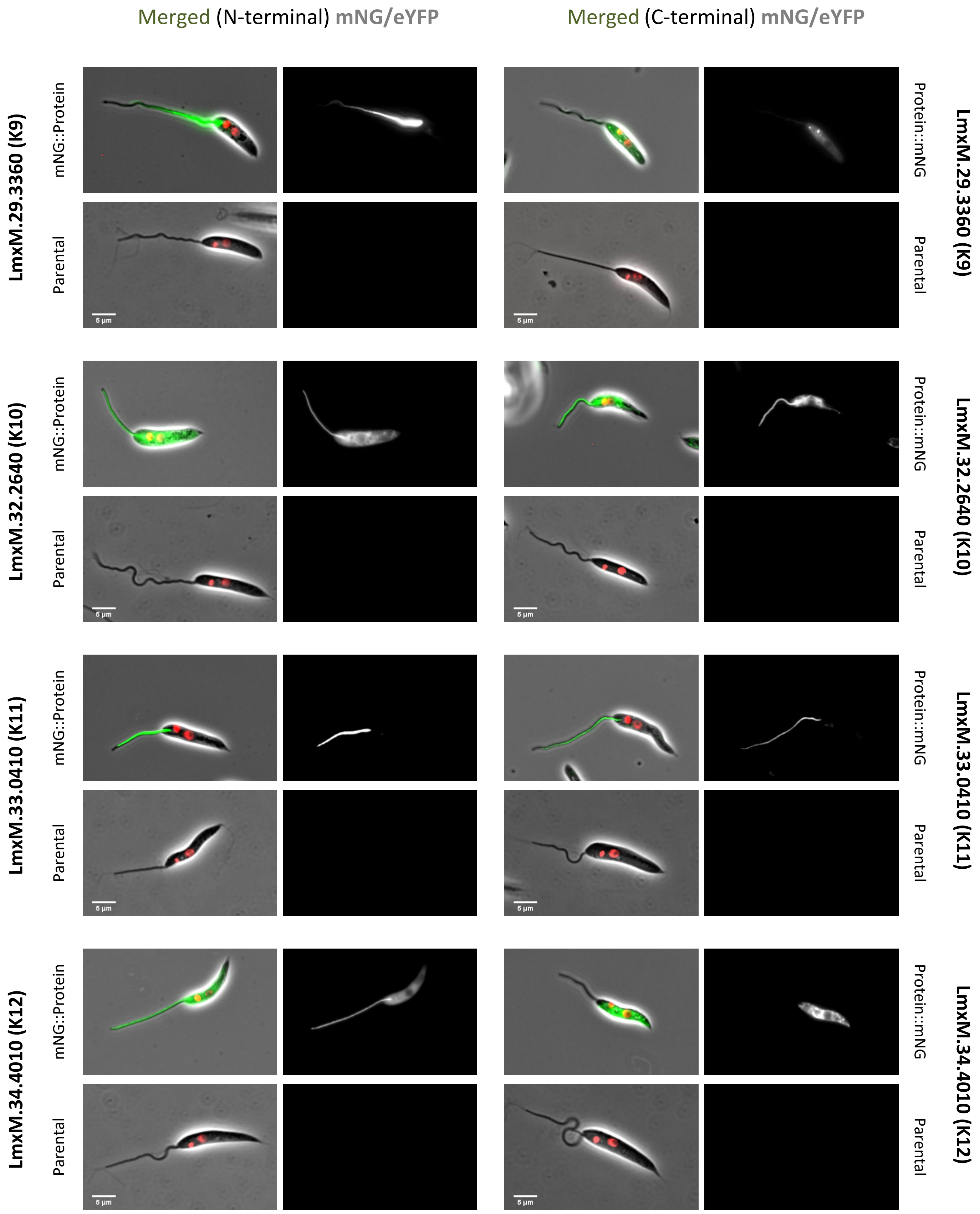

Merged (N-terminal) mNG/eYFP
Merged (C-terminal) mNG/eYFP
LmxM.29.3360 (K9)
mNG::Protein
Parental
Protein::mNG
Parental
LmxM.32.2640 (K10)
LmxM.33.0410 (K11)
LmxM.34.4010 (K12)
5 µm

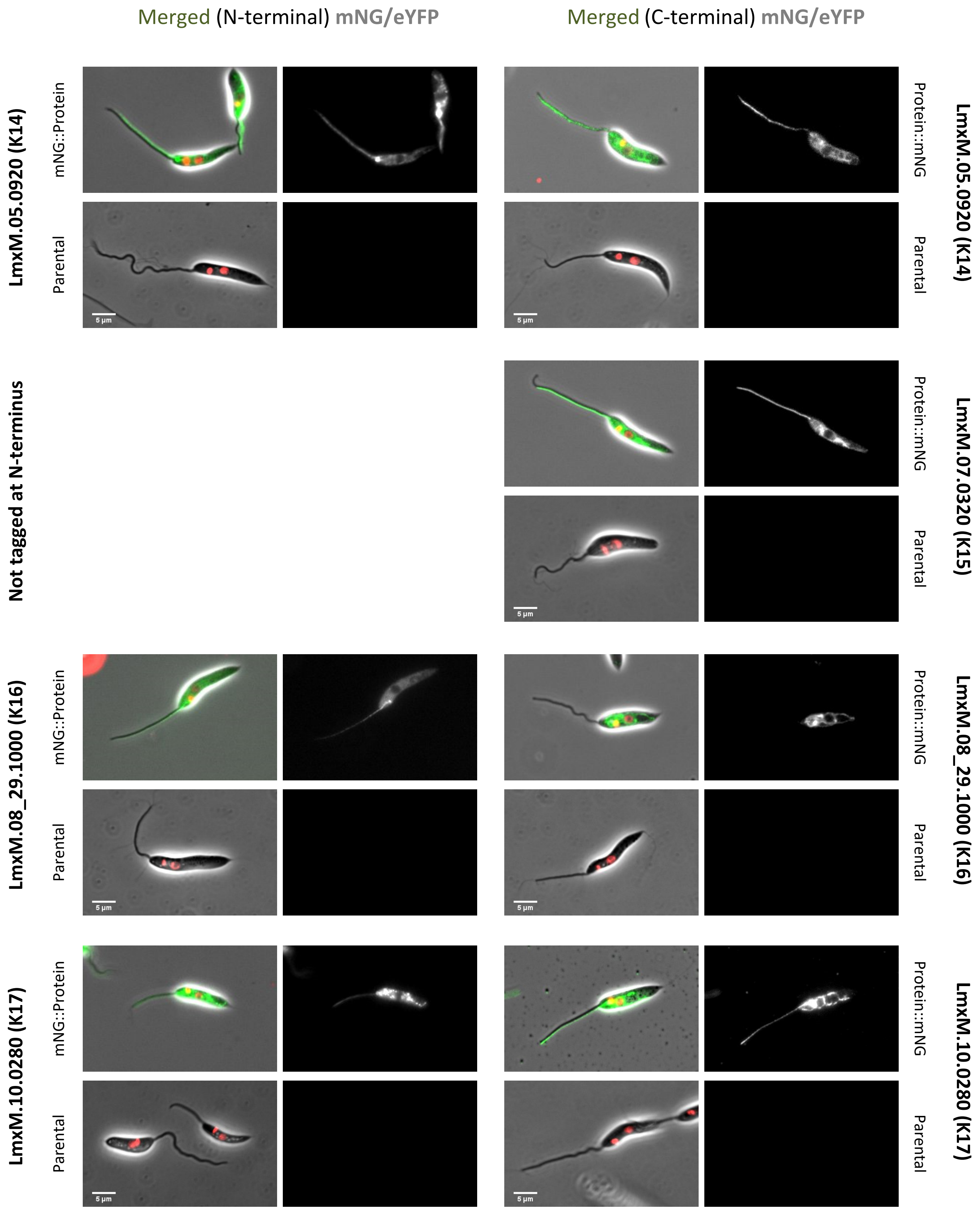

Merged (N-terminal) mNG/eYFP
Merged (C-terminal) mNG/eYFP
LmxM.05.0920 (K14)
mNG::Protein
Parental
Protein::mNG
Parental
Not tagged at N-terminus
LmxM.07.0320 (K15)
LmxM.08_29.1000 (K16)
LmxM.10.0280 (K17)
5 µm

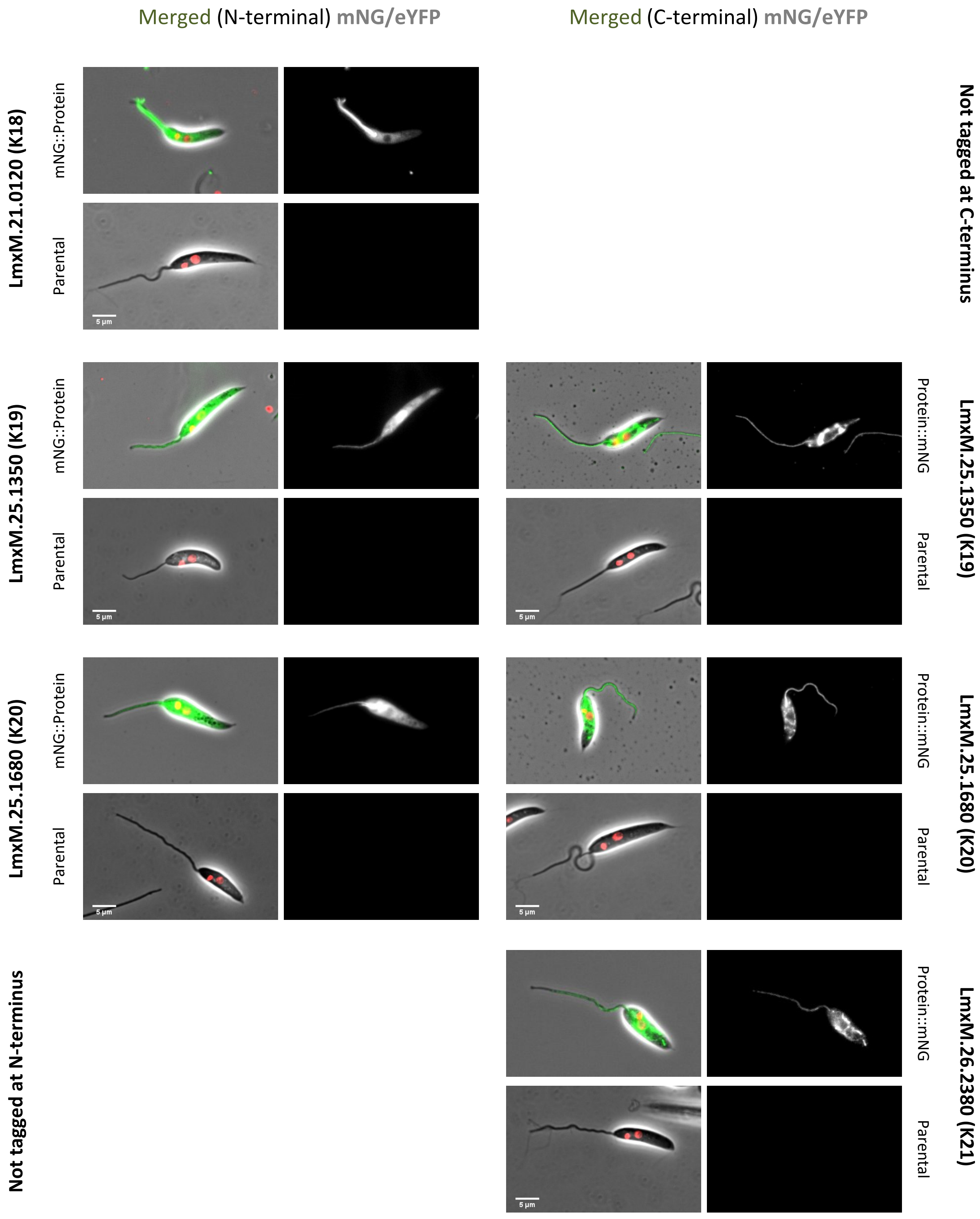
Merged (N-terminal) mNG/eYFP
Merged (C-terminal) mNG/eYFP
LmxM.21.0120 (K18)
mNG::Protein
Parental
Not tagged at C-terminus
LmxM.25.1350 (K19)
Protein::mNG
LmxM.25.1680 (K20)
LmxM.26.2380 (K21)
Not tagged at N-terminus
5 µm

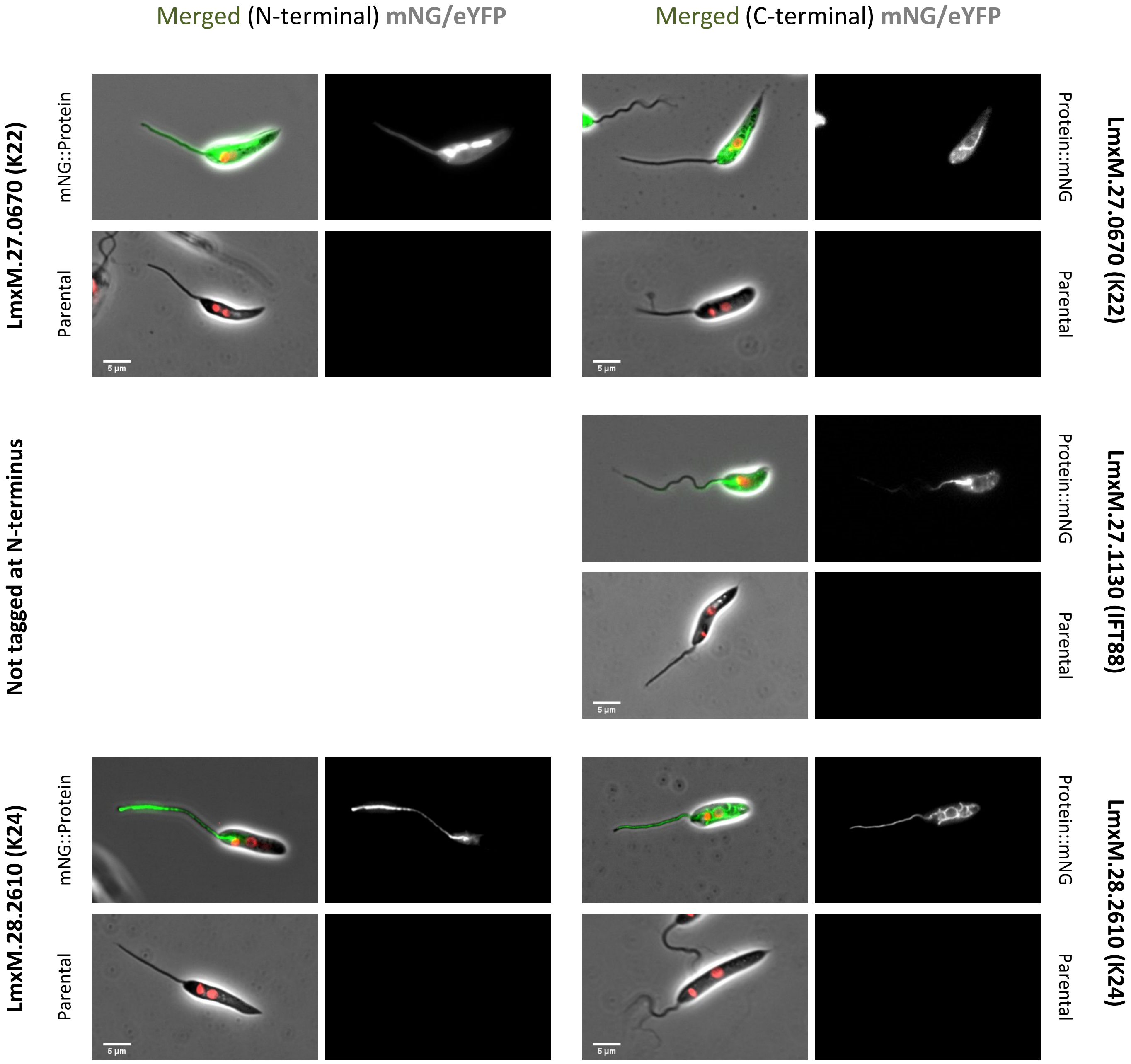
Merged (N-terminal) mNG/eYFP
Merged (C-terminal) mNG/eYFP
LmxM.27.0670 (K22)
mNG::Protein
Parental
Protein::mNG
Parental
LmxM.27.0670 (K22)
Not tagged at N-terminus
Protein::mNG
Parental
LmxM.27.1130 (IFT88)
LmxM.28.2610 (K24)
mNG::Protein
Parental
Protein::mNG
Parental
LmxM.28.2610 (K24)
5 µm

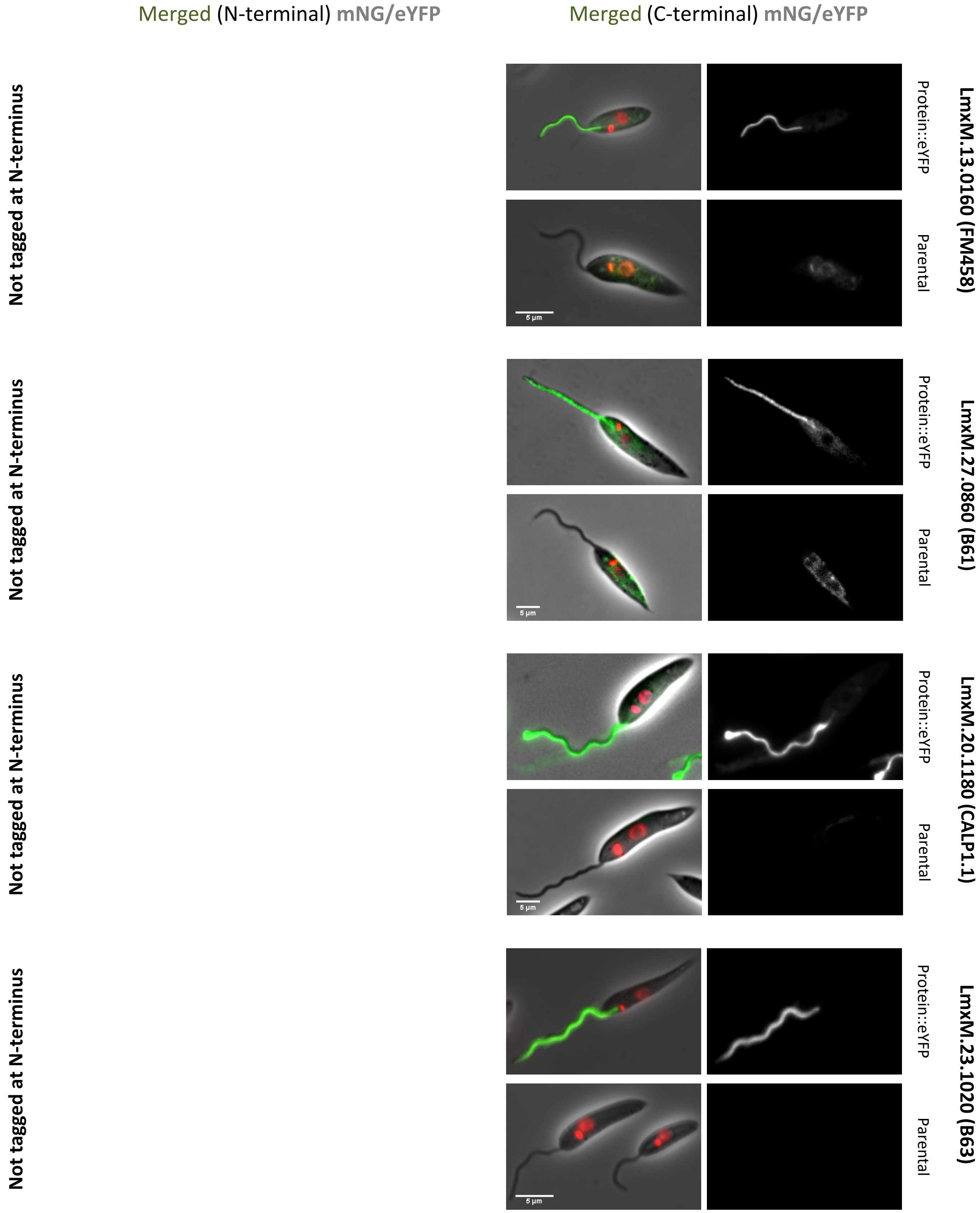

Merged (N-terminal) mNG/eYFP
Merged (C-terminal) mNG/eYFP
Not tagged at N-terminus
Not tagged at N-terminus
Not tagged at N-terminus
Not tagged at N-terminus
Protein::eYFP
Parental
LmxM.13.0160 (FM458)
Protein::eYFP
Parental
LmxM.27.0860 (B61)
Protein::eYFP
Parental
LmxM.20.1180 (CALP1.1)
Protein::eYFP
Parental
LmxM.23.1020 (B63)
5 µm
5 µm
5 µm
5 µm

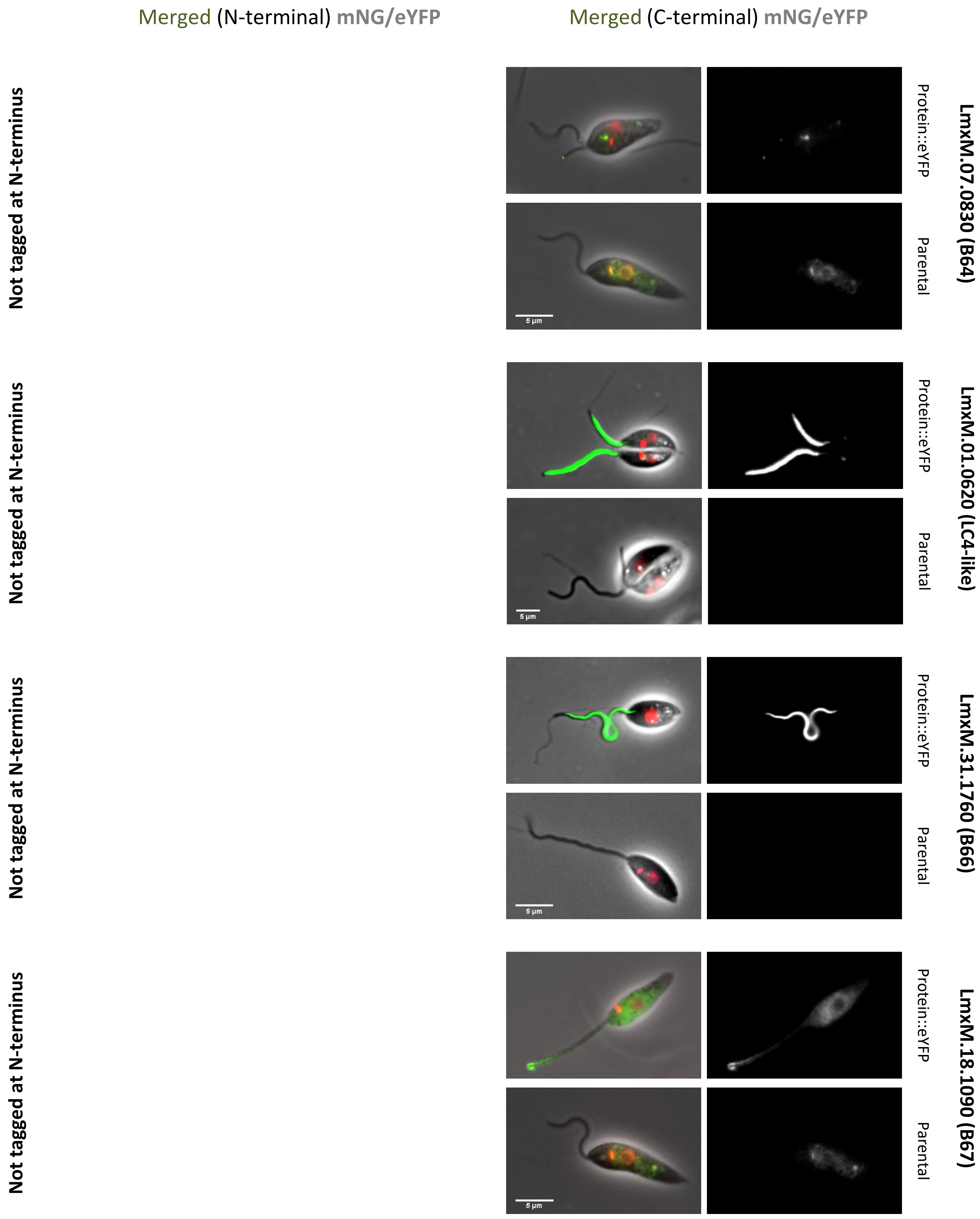
Merged (N-terminal) mNG/eYFP
Merged (C-terminal) mNG/eYFP
Not tagged at N-terminus
LmxM.07.0830 (B64)
Protein::eYFP
Parental
5 µm
Not tagged at N-terminus
LmxM.01.0620 (LC4-like)
Protein::eYFP
Parental
5 µm
Not tagged at N-terminus
LmxM.31.1760 (B66)
Protein::eYFP
Parental
5 µm
Not tagged at N-terminus
LmxM.18.1090 (B67)
Protein::eYFP
Parental
5 µm

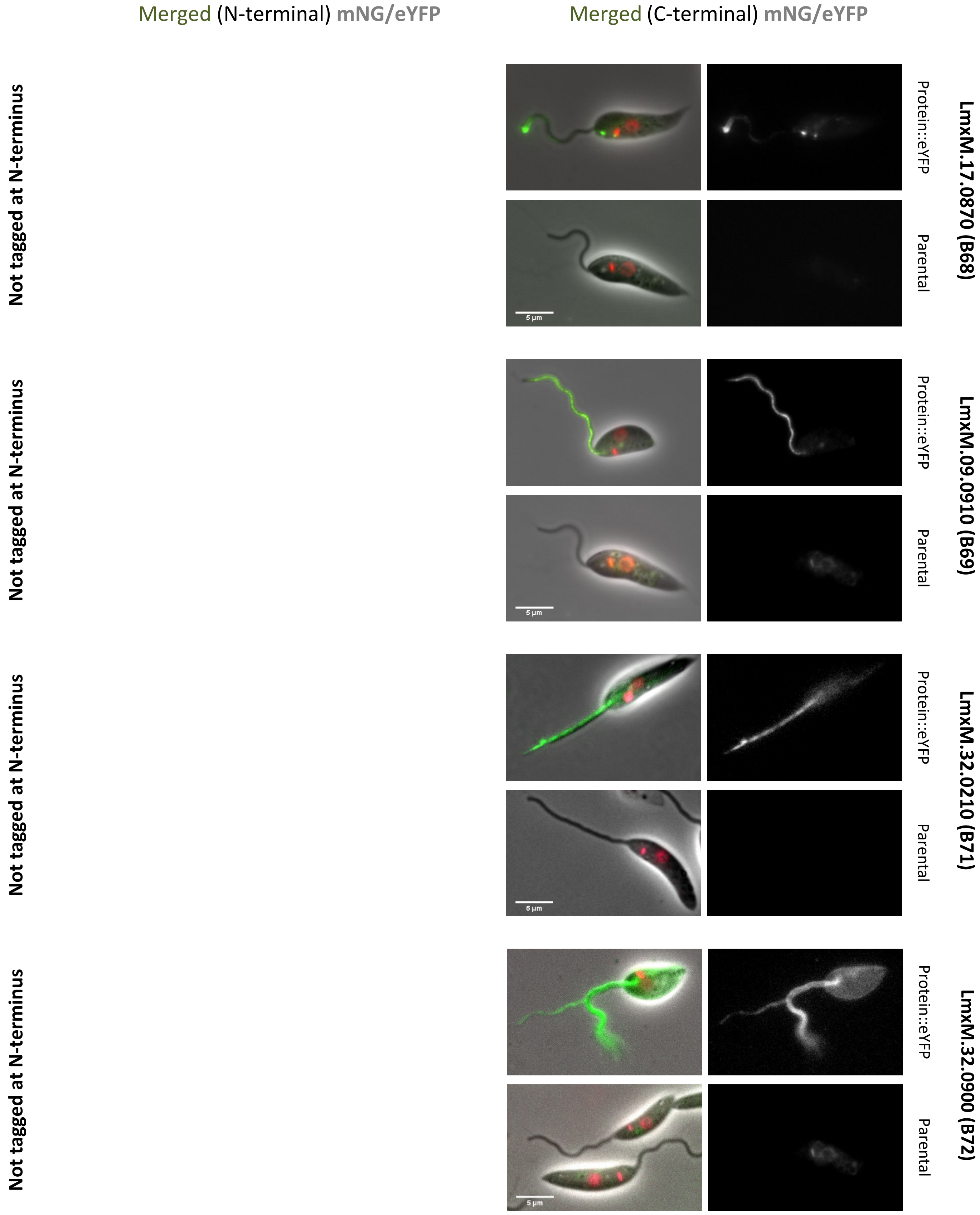

Merged (N-terminal) mNG/eYFP
Merged (C-terminal) mNG/eYFP
Not tagged at N-terminus
Protein::eYFP
Parental
5 µm
LmxM.17.0870 (B68)
Not tagged at N-terminus
Protein::eYFP
Parental
5 µm
LmxM.09.0910 (B69)
Not tagged at N-terminus
Protein::eYFP
Parental
5 µm
LmxM.32.0210 (B71)
Not tagged at N-terminus
Protein::eYFP
Parental
5 µm
LmxM.32.0900 (B72)

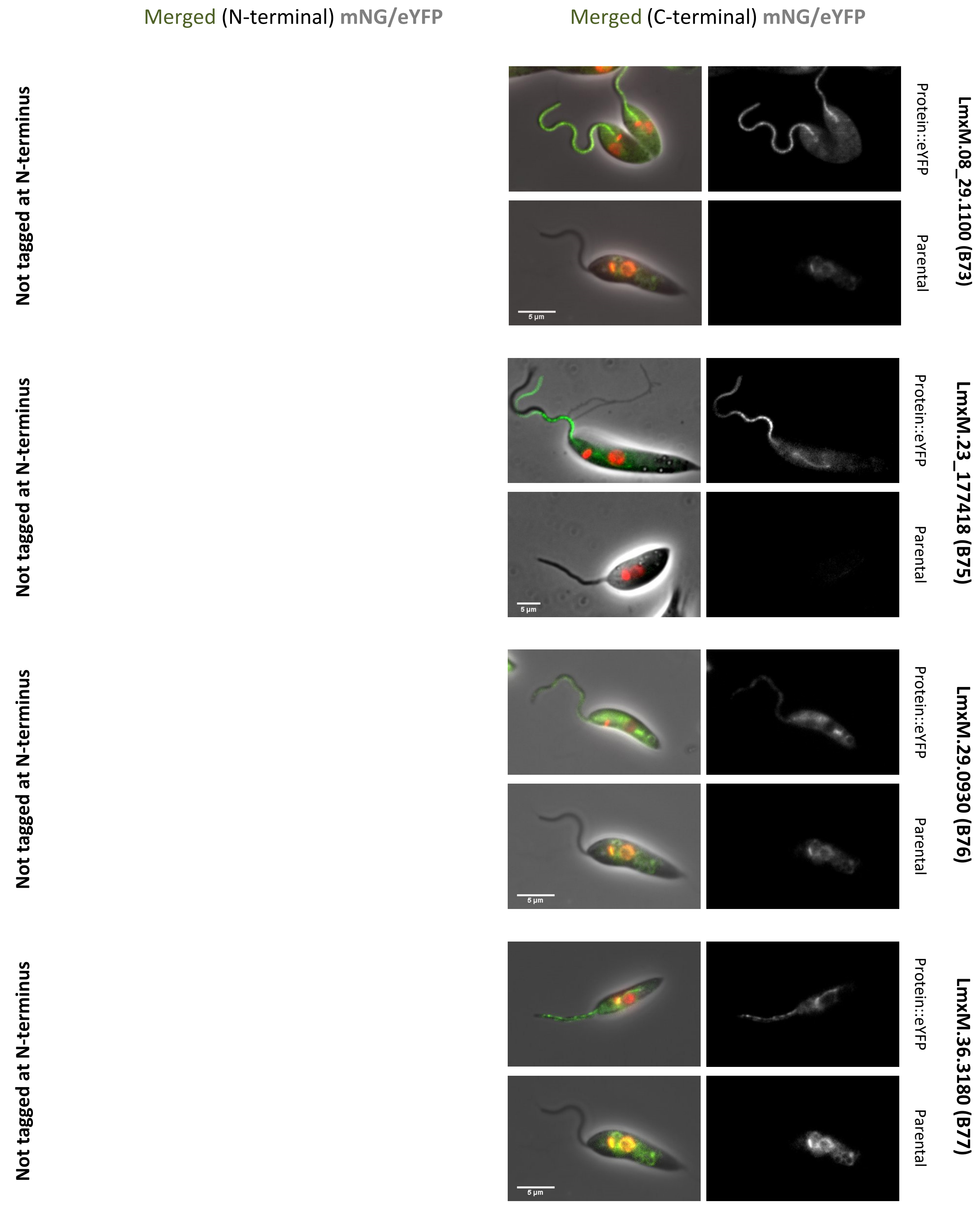
Merged (N-terminal) mNG/eYFP
Merged (C-terminal) mNG/eYFP
Not tagged at N-terminus
Protein::eYFP
Parental
LmxM.08_29.1100 (B73)
5 µm
Not tagged at N-terminus
Protein::eYFP
Parental
LmxM.23_177418 (B75)
5 µm
Not tagged at N-terminus
Protein::eYFP
Parental
LmxM.29.0930 (B76)
5 µm
Not tagged at N-terminus
Protein::eYFP
Parental
LmxM.36.3180 (B77)
5 µm

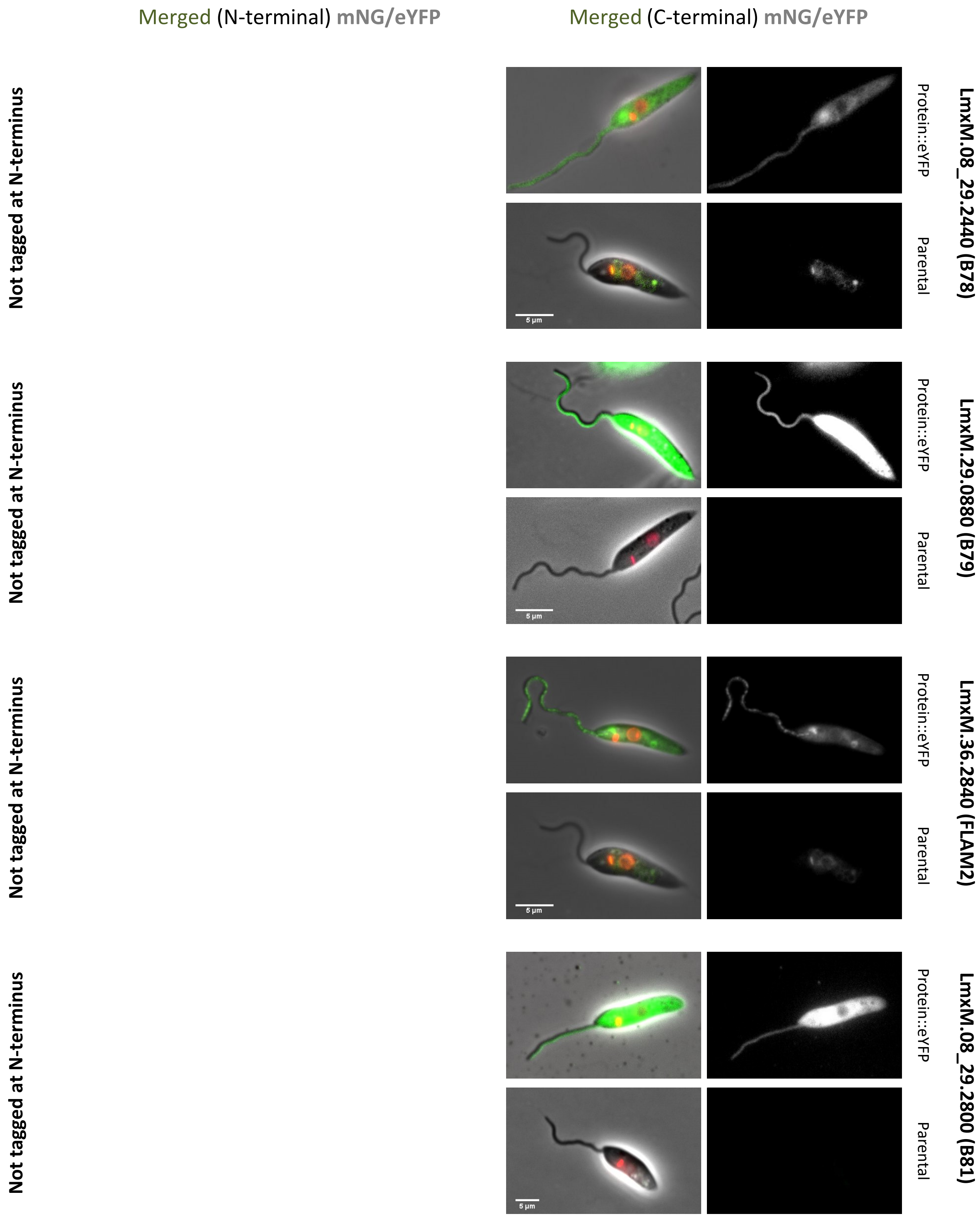

Merged (N-terminal) mNG/eYFP
Merged (C-terminal) mNG/eYFP
Not tagged at N-terminus
LmxM.08_29.2440 (B78)
Protein::eYFP
Parental
5 µm
Not tagged at N-terminus
LmxM.29.0880 (B79)
Protein::eYFP
Parental
5 µm
Not tagged at N-terminus
LmxM.36.2840 (FLAM2)
Protein::eYFP
Parental
5 µm
Not tagged at N-terminus
LmxM.08_29.2800 (B81)
Protein::eYFP
Parental
5 µm

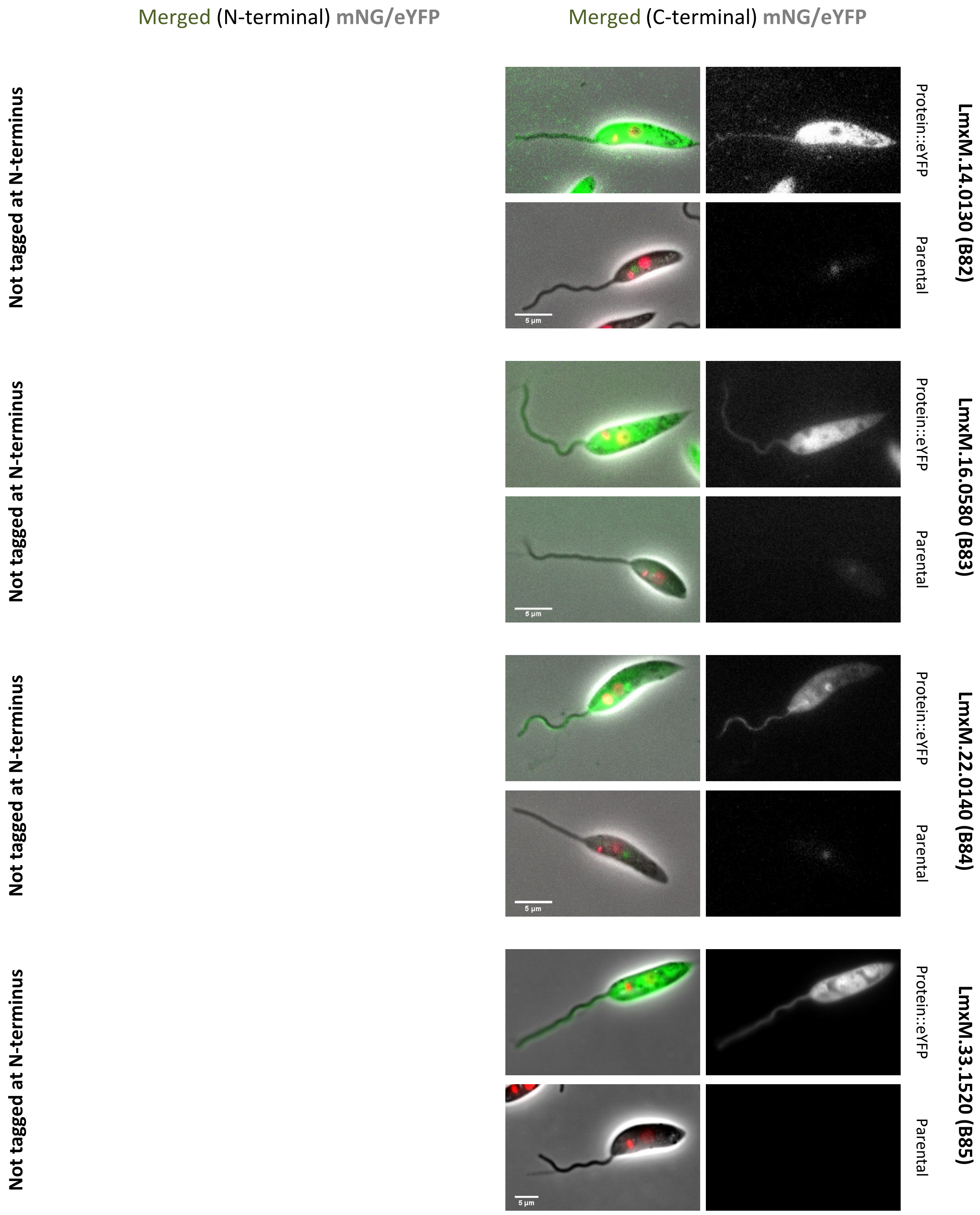

Merged (N-terminal) mNG/eYFP
Merged (C-terminal) mNG/eYFP
Not tagged at N-terminus
Not tagged at N-terminus
Not tagged at N-terminus
Not tagged at N-terminus
LmxM.14.0130 (B82)
LmxM.16.0580 (B83)
LmxM.22.0140 (B84)
LmxM.33.1520 (B85)
Protein::eYFP
Parental
Protein::eYFP
Parental
Protein::eYFP
Parental
Protein::eYFP
Parental
5 µm
5 µm
5 µm
5 µm

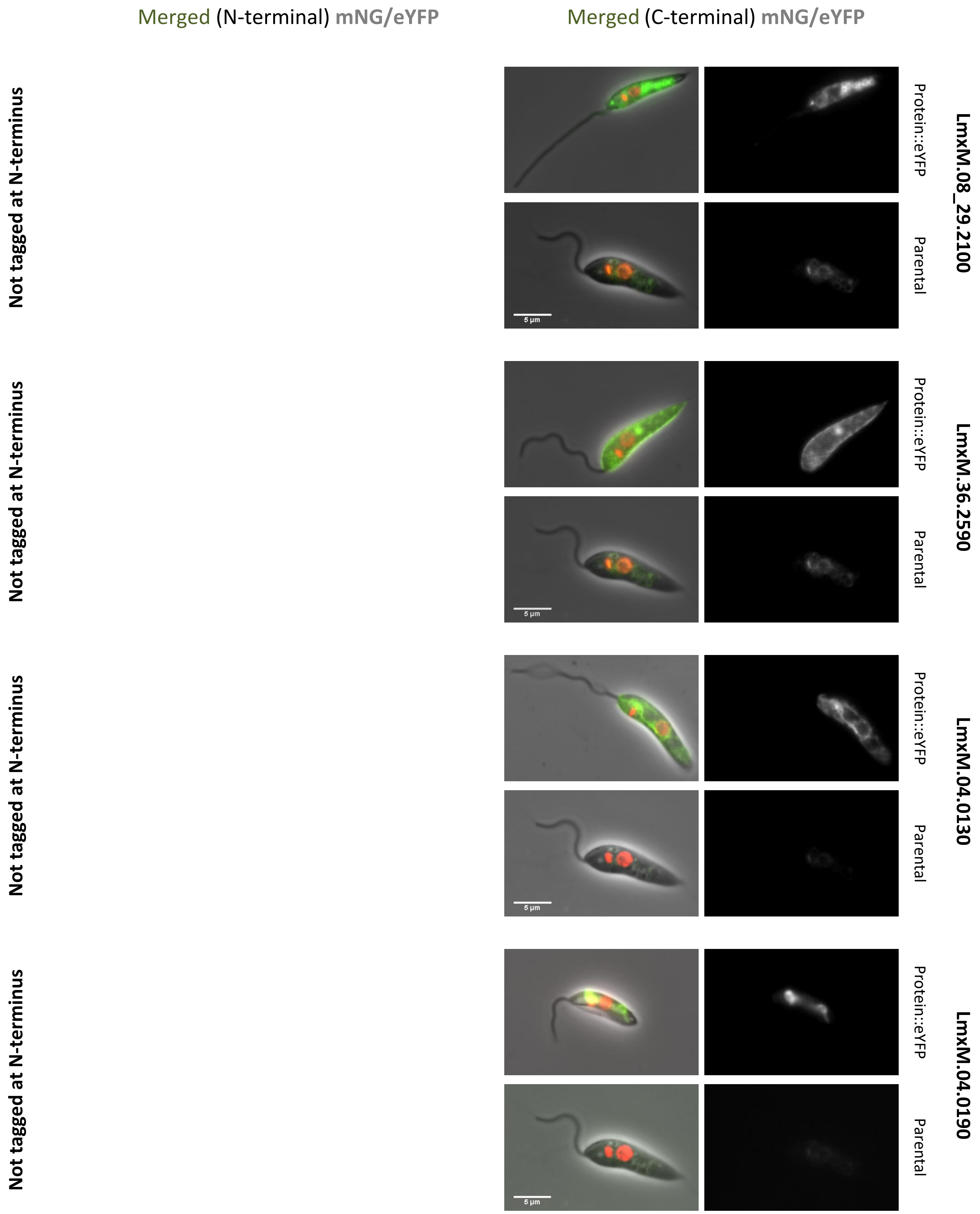
Merged (N-terminal) mNG/eYFP
Merged (C-terminal) mNG/eYFP
Not tagged at N-terminus
Not tagged at N-terminus
Not tagged at N-terminus
Not tagged at N-terminus
LmxM.08_29.2100
LmxM.36.2590
LmxM.04.0130
LmxM.04.0190
Protein::eYFP
Parental
Protein::eYFP
Parental
Protein::eYFP
Parental
Protein::eYFP
Parental
5 µm
5 µm
5 µm
5 µm

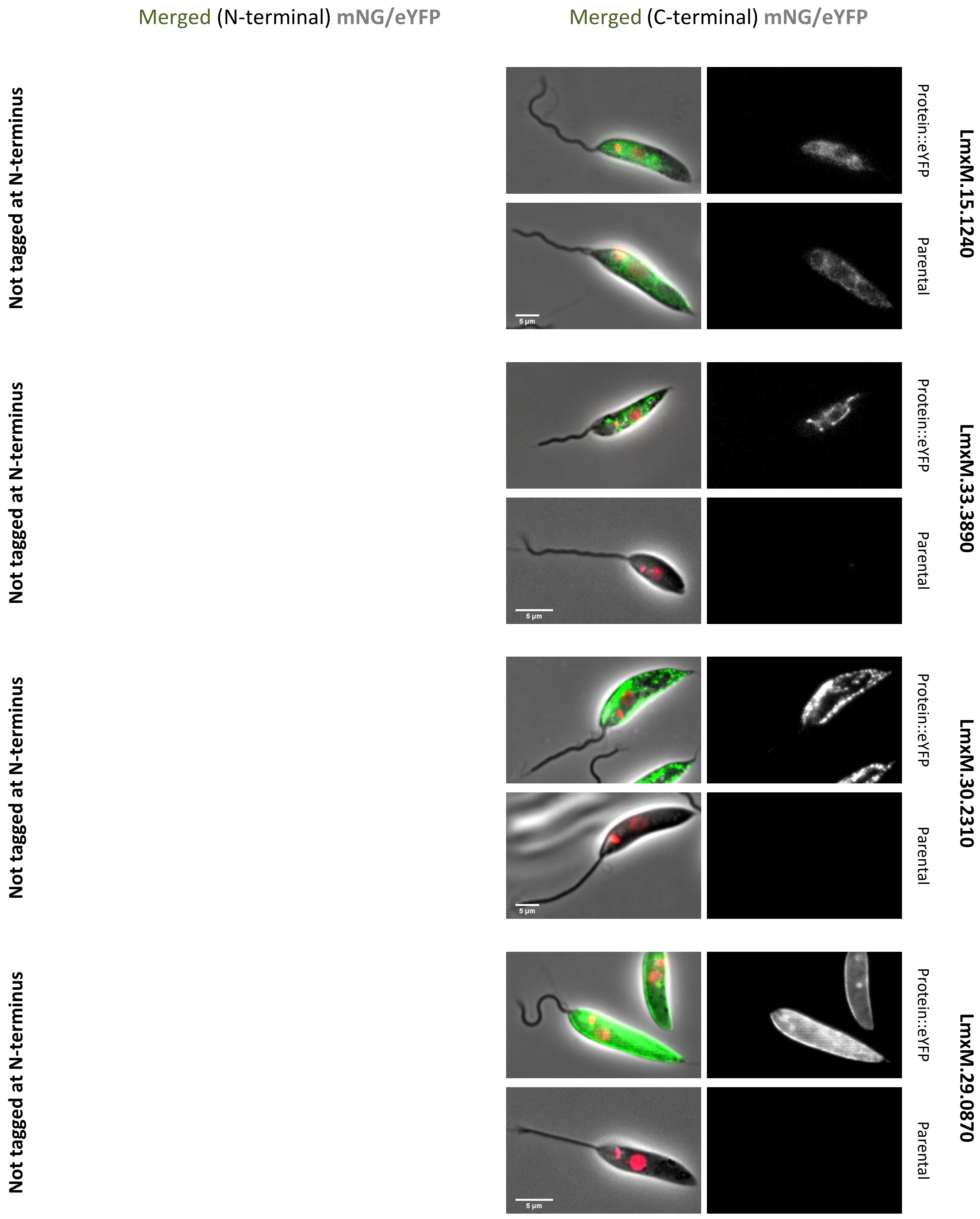
Merged (N-terminal) mNG/eYFP
Merged (C-terminal) mNG/eYFP
Not tagged at N-terminus
Not tagged at N-terminus
Not tagged at N-terminus
Not tagged at N-terminus
Protein::eYFP
Parental
LmxM.15.1240
LmxM.33.3890
LmxM.30.2310
LmxM.29.0870
5 µm

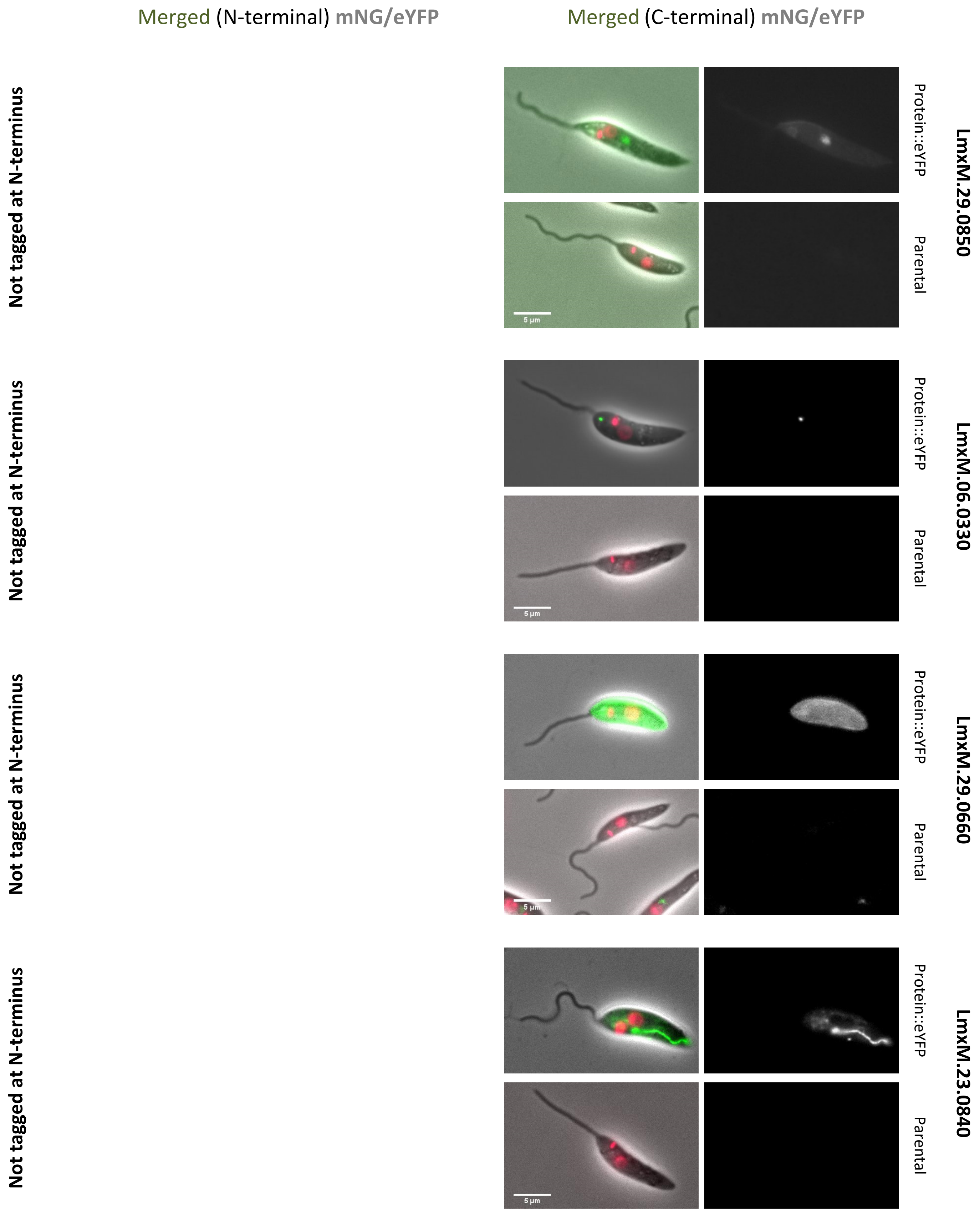
Merged (N-terminal) mNG/eYFP
Merged (C-terminal) mNG/eYFP
Not tagged at N-terminus
Protein::eYFP
Parental
LmxM.29.0850
5 µm
Not tagged at N-terminus
Protein::eYFP
Parental
LmxM.06.0330
5 µm
Not tagged at N-terminus
Protein::eYFP
Parental
LmxM.29.0660
5 µm
Not tagged at N-terminus
Protein::eYFP
Parental
LmxM.23.0840
5 µm

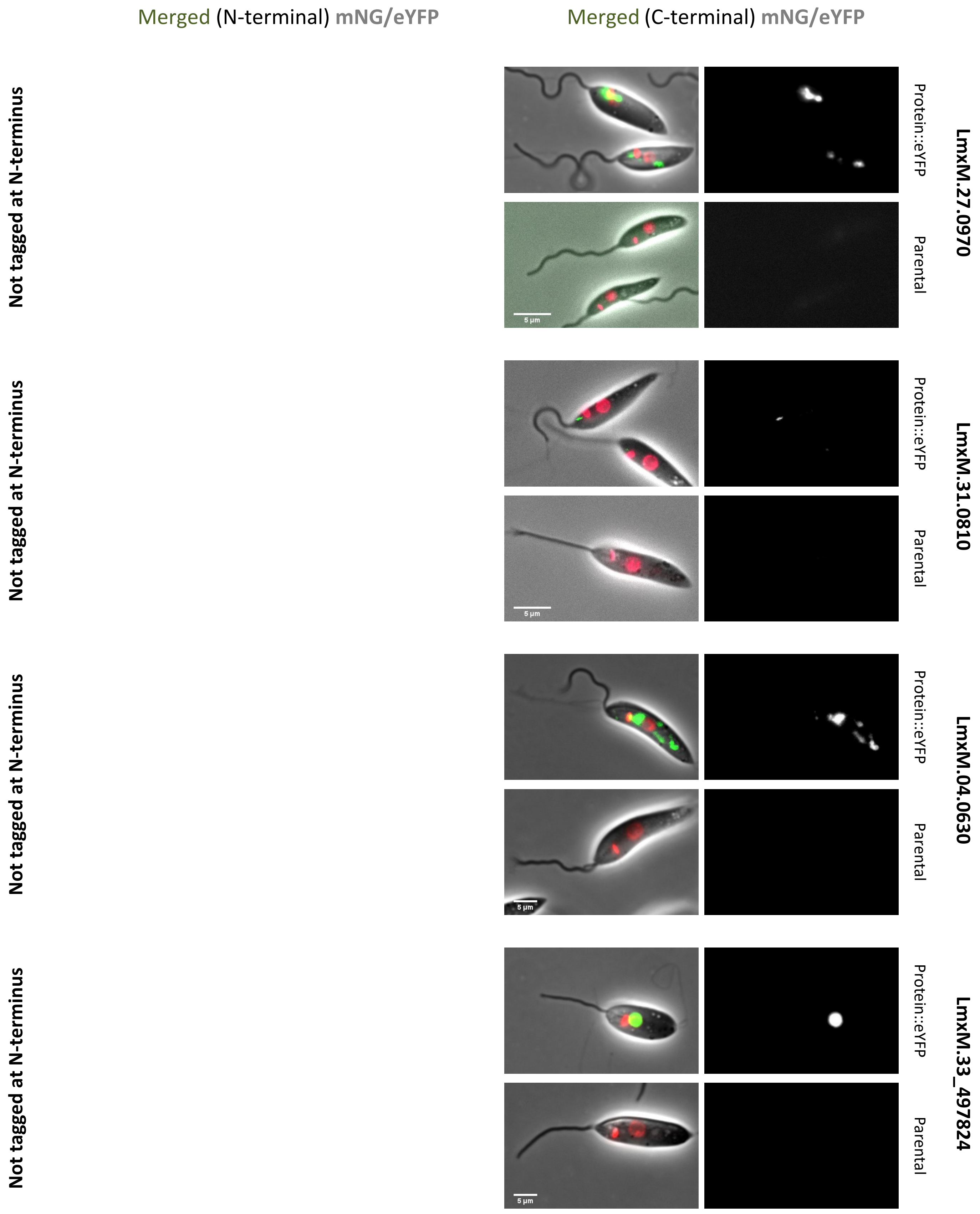

Merged (N-terminal) mNG/eYFP
Merged (C-terminal) mNG/eYFP
Not tagged at N-terminus
Protein::eYFP
Parental
LmxM.27.0970
5 µm
Not tagged at N-terminus
Protein::eYFP
Parental
LmxM.31.0810
5 µm
Not tagged at N-terminus
Protein::eYFP
Parental
LmxM.04.0630
5 µm
Not tagged at N-terminus
Protein::eYFP
Parental
LmxM.33_497824
5 µm

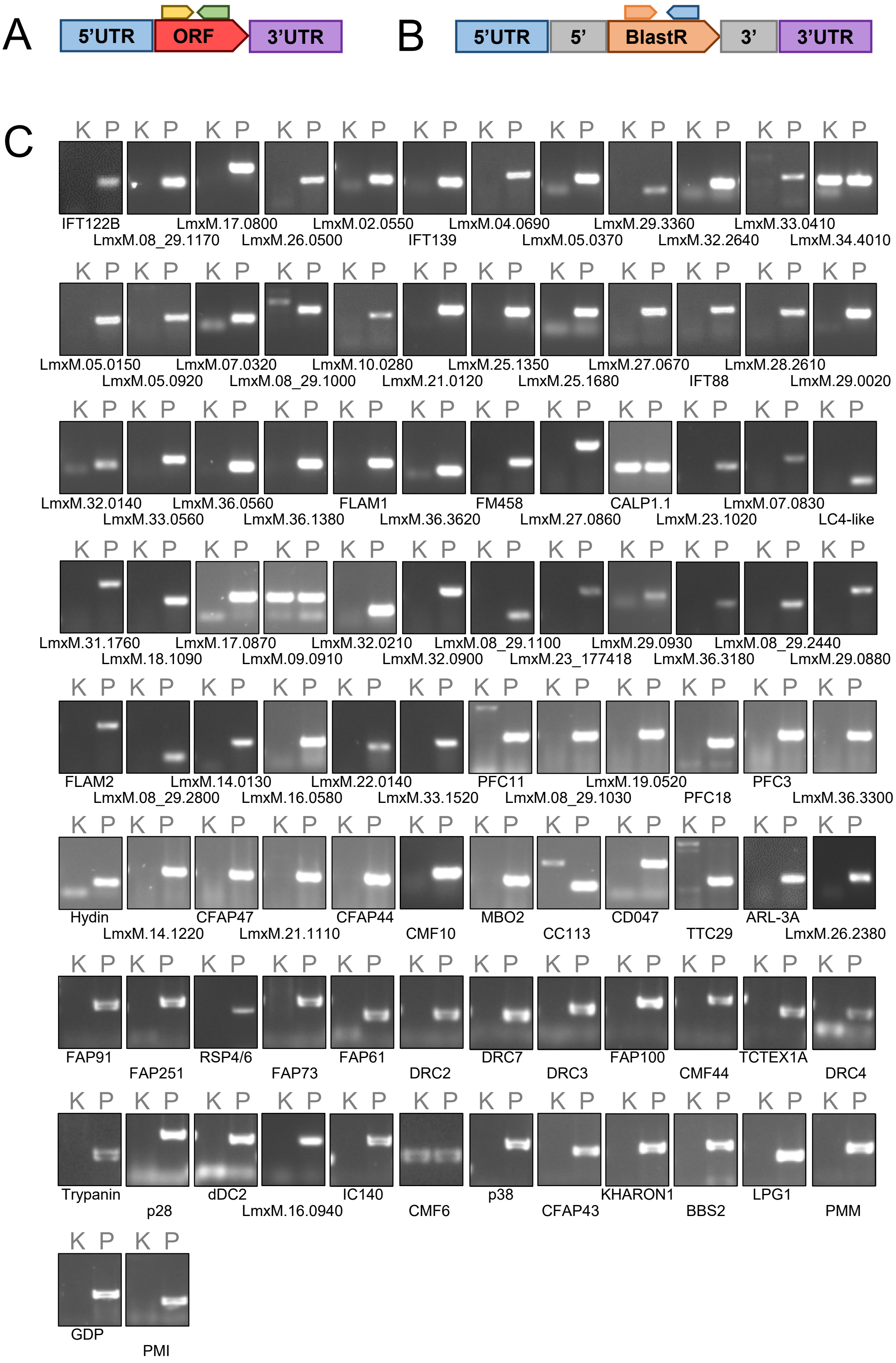
A
5'UTR
ORF
3'UTR
B
5'UTR
5'
BlastR
3'
3'UTR
C
K P
IFT122B
LmxM.08_29.1170
LmxM.17.0800
LmxM.26.0500
LmxM.02.0550
IFT139
LmxM.04.0690
LmxM.05.0370
LmxM.29.3360
LmxM.32.2640
LmxM.33.0410
LmxM.34.4010
LmxM.05.0150
LmxM.05.0920
LmxM.07.0320
LmxM.08_29.1000
LmxM.10.0280
LmxM.21.0120
LmxM.25.1350
LmxM.25.1680
LmxM.27.0670
IFT88
LmxM.28.2610
LmxM.29.0020
LmxM.32.0140
LmxM.33.0560
LmxM.36.0560
LmxM.36.1380
FLAM1
LmxM.36.3620
FM458
LmxM.27.0860
CALP1.1
LmxM.23.1020
LmxM.07.0830
LC4-like
LmxM.31.1760
LmxM.18.1090
LmxM.17.0870
LmxM.09.0910
LmxM.32.0210
LmxM.32.0900
LmxM.08_29.1100
LmxM.23_177418
LmxM.29.0930
LmxM.36.3180
LmxM.08_29.2440
LmxM.29.0880
FLAM2
LmxM.08_29.2800
LmxM.14.0130
LmxM.16.0580
LmxM.22.0140
LmxM.33.1520
PFC11
LmxM.08_29.1030
LmxM.19.0520
PFC18
PFC3
LmxM.36.3300
Hydin
LmxM.14.1220
CFAP47
LmxM.21.1110
CFAP44
CMF10
MBO2
CC113
CD047
TTC29
ARL-3A
LmxM.26.2380
FAP91
FAP251
RSP4/6
FAP73
FAP61
DRC2
DRC7
DRC3
FAP100
CMF44
TCTEX1A
DRC4
Trypanin
p28
dDC2
LmxM.16.0940
IC140
CMF6
p38
CFAP43
KHARON1
BBS2
LPG1
PMM
GDP
PMI

D

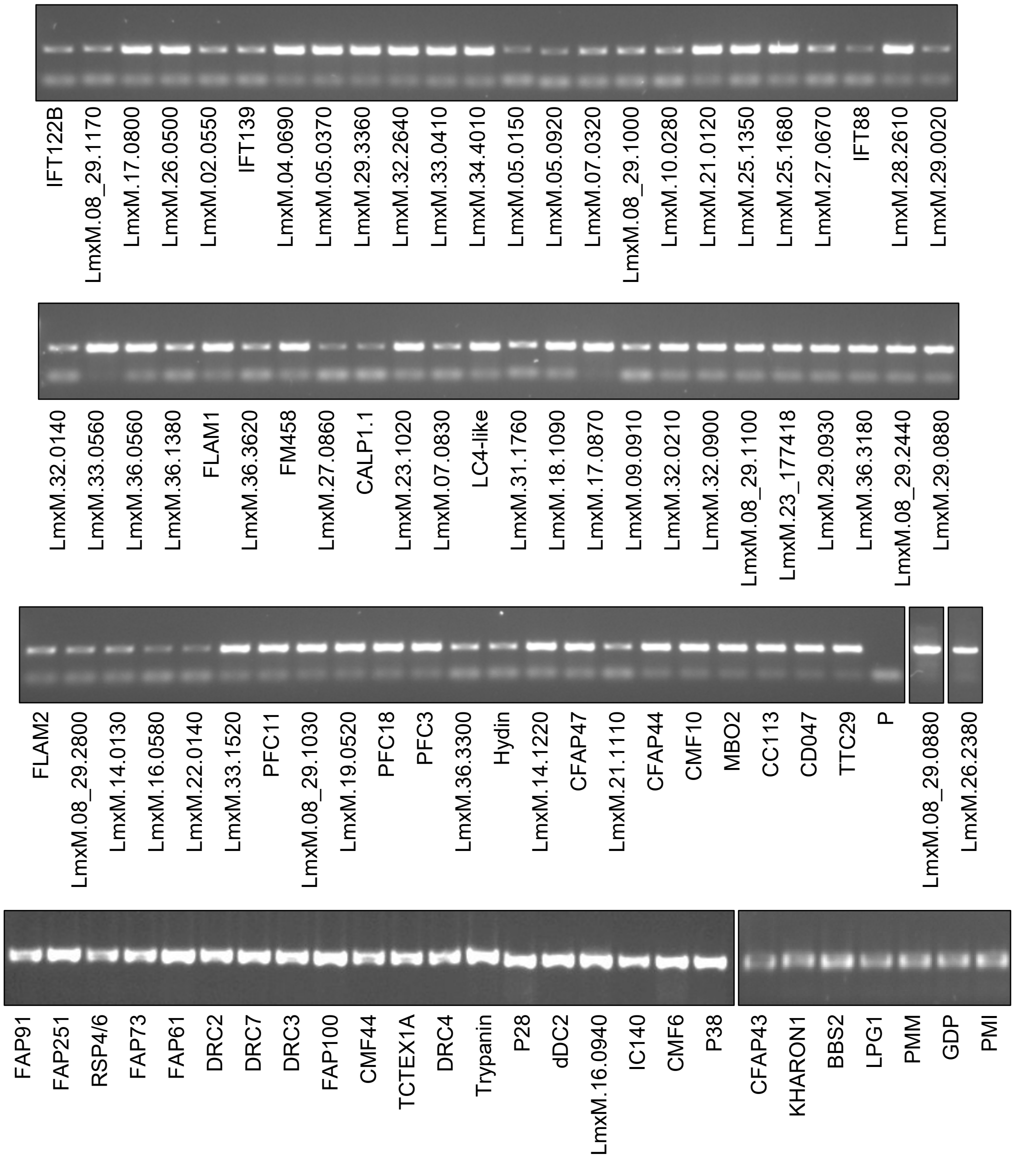
IFT122B
LmxM.08_29.1170
LmxM.17.0800
LmxM.26.0500
LmxM.02.0550
IFT139
LmxM.04.0690
LmxM.05.0370
LmxM.29.3360
LmxM.32.2640
LmxM.33.0410
LmxM.34.4010
LmxM.05.0150
LmxM.05.0920
LmxM.07.0320
LmxM.08_29.1000
LmxM.10.0280
LmxM.21.0120
LmxM.25.1350
LmxM.25.1680
LmxM.27.0670
IFT88
LmxM.28.2610
LmxM.29.0020
LmxM.32.0140
LmxM.33.0560
LmxM.36.0560
LmxM.36.1380
FLAM1
LmxM.36.3620
FM458
LmxM.27.0860
CALP1.1
LmxM.23.1020
LmxM.07.0830
LC4-like
LmxM.31.1760
LmxM.18.1090
LmxM.17.0870
LmxM.09.0910
LmxM.32.0210
LmxM.32.0900
LmxM.08_29.1100
LmxM.23_177418
LmxM.29.0930
LmxM.36.3180
LmxM.08_29.2440
LmxM.29.0880
FLAM2
LmxM.08_29.2800
LmxM.14.0130
LmxM.16.0580
LmxM.22.0140
LmxM.33.1520
PFC11
LmxM.08_29.1030
LmxM.19.0520
PFC18
PFC3
LmxM.36.3300
Hydin
LmxM.14.1220
CFAP47
LmxM.21.1110
CFAP44
CMF10
MBO2
CC113
CD047
TTC29
P
LmxM.08_29.0880
LmxM.26.2380
FAP91
FAP251
RSP4/6
FAP73
FAP61
DRC2
DRC7
DRC3
FAP100
CMF44
TCTEX1A
DRC4
Trypanin
P28
dDC2
LmxM.16.0940
IC140
CMF6
P38
CFAP43
KHARON1
BBS2
LPG1
PMM
GDP
PMI

Figure S8

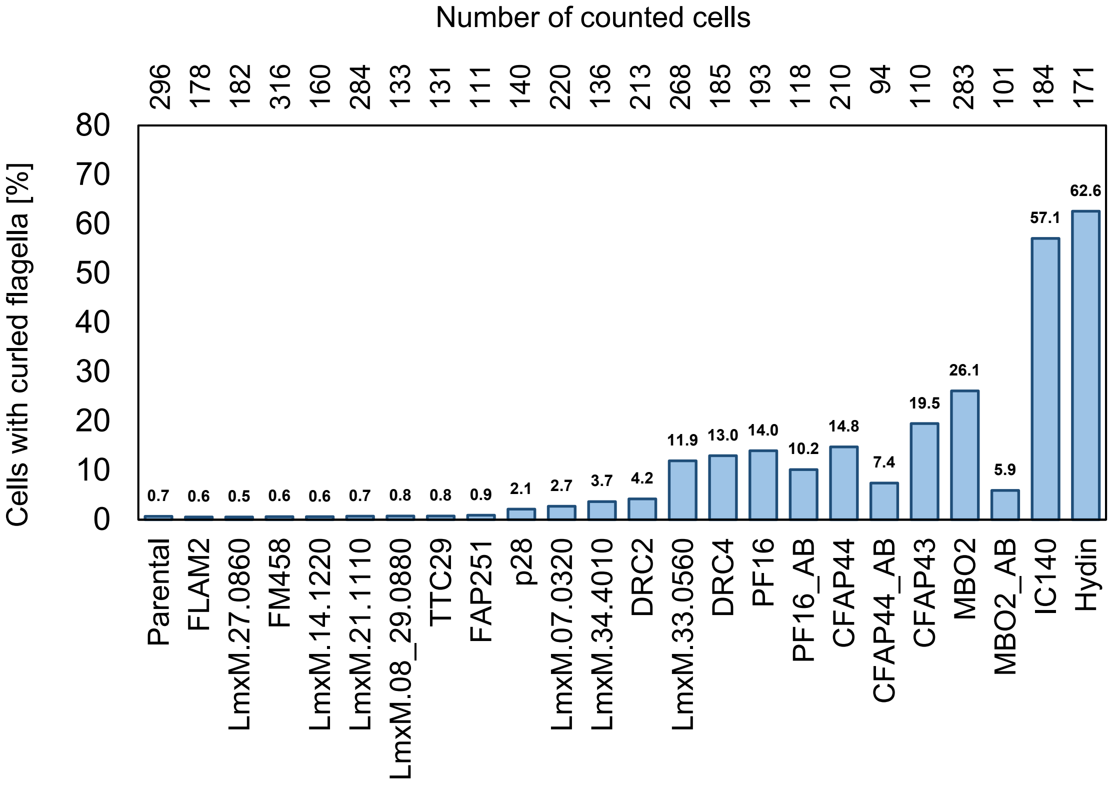
Number of counted cells
296 178 182 316 160 284 133 131 111 140 220 136 213 268 185 193 118 210 94 110 283 101 184 171
Cells with curled flagella [%]
0 10 20 30 40 50 60 70 80
0.7 0.6 0.5 0.6 0.6 0.7 0.8 0.8 0.9 2.1 2.7 3.7 4.2 11.9 13.0 14.0 10.2 14.8 7.4 19.5 26.1 5.9 57.1 62.6
Parental
FLAM2
LmxM.27.0860
FM458
LmxM.14.1220
LmxM.21.1110
LmxM.08_29.0880
TTC29
FAP251
p28
LmxM.07.0320
LmxM.34.4010
DRC2
LmxM.33.0560
DRC4
PF16
PF16_AB
CFAP44
CFAP44_AB
CFAP43
MBO2
MBO2_AB
IC140
Hydin

Figure S9

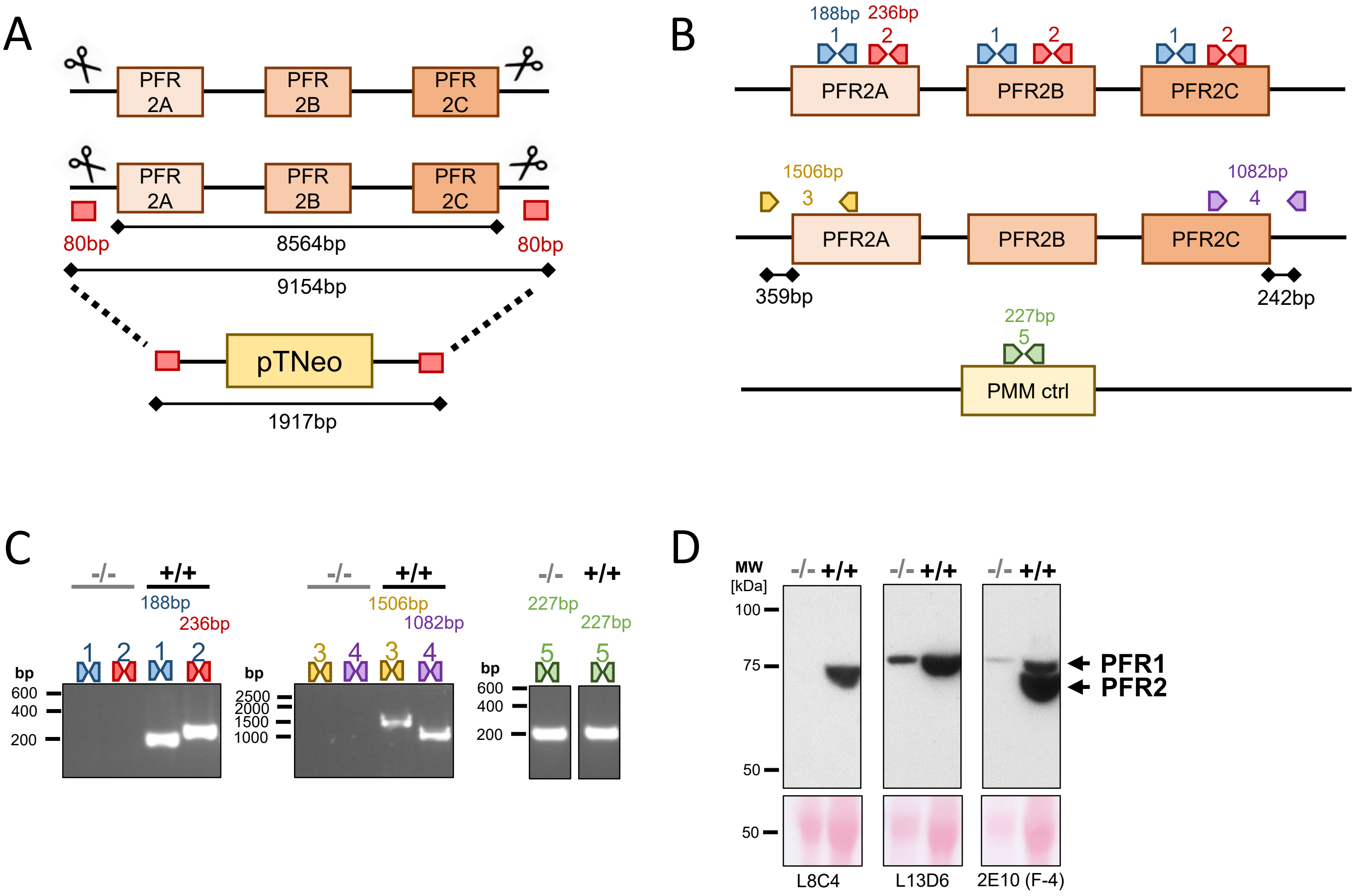
A
PFR 2A
PFR 2B
PFR 2C
80bp
8564bp
80bp
9154bp
pTNeo
1917bp
B
188bp
236bp
1
2
PFR2A
PFR2B
PFR2C
1506bp
3
1082bp
4
359bp
242bp
227bp
5
PMM ctrl
C
-/-
+/+
188bp
236bp
1506bp
1082bp
227bp
bp
600
400
200
2500
2000
1500
1000
D
MW
[kDa]
100
75
50
PFR1
PFR2
L8C4
L13D6
2E10 (F-4)

Figure S10

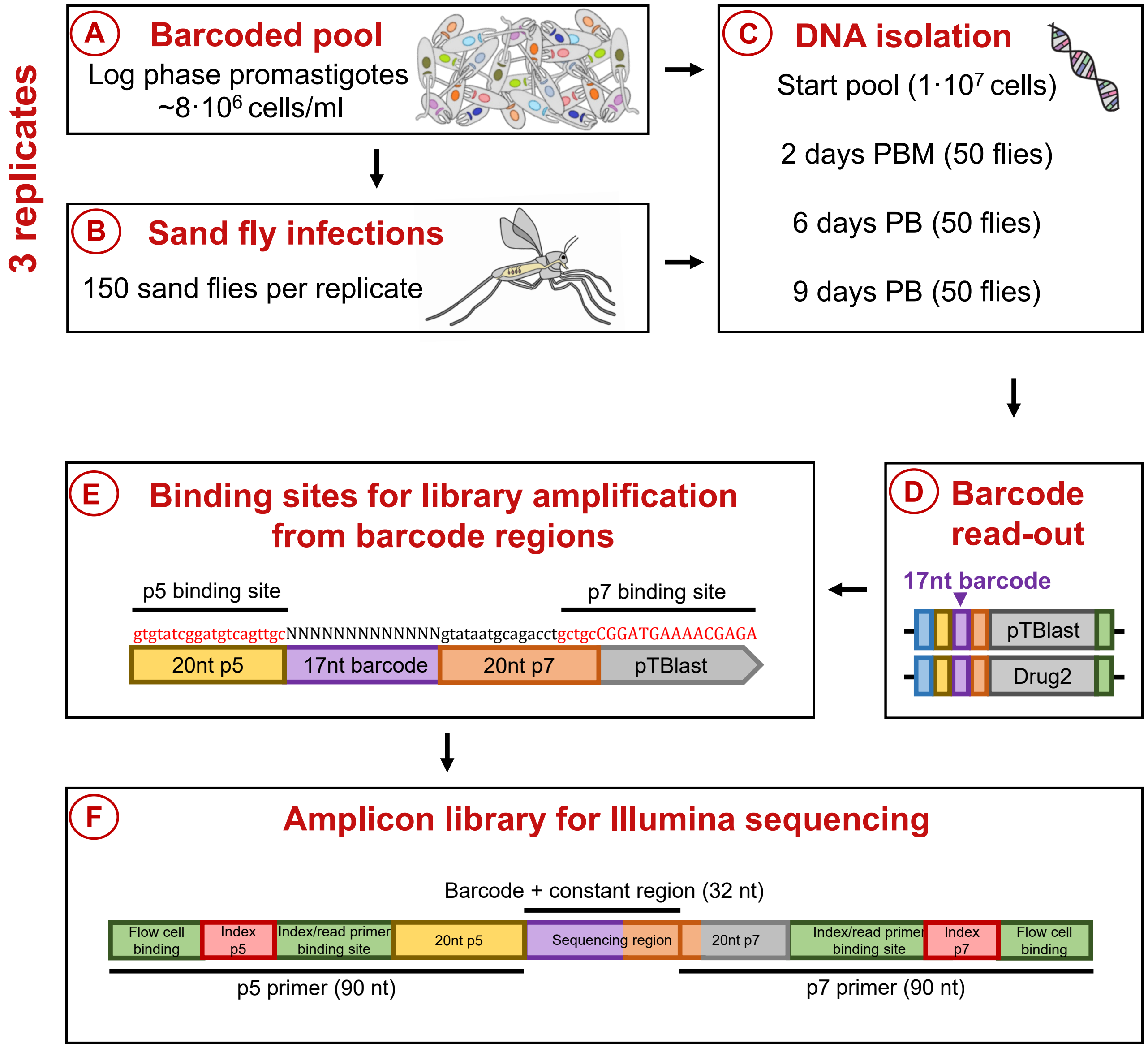
3 replicates
A
Barcoded pool
Log phase promastigotes
~8·10^6 cells/ml
B
Sand fly infections
150 sand flies per replicate
C
DNA isolation
Start pool (1·10^7 cells)
2 days PBM (50 flies)
6 days PB (50 flies)
9 days PB (50 flies)
D
Barcode read-out
17nt barcode
pTBlast
Drug2
E
Binding sites for library amplification from barcode regions
p5 binding site
p7 binding site
gtgtatcggatgtcagttgcNNNNNNNNNNNNNgtataatgcagacctgctgcCGGATGAAAACGAGA
20nt p5
17nt barcode
20nt p7
pTBlast
F
Amplicon library for Illumina sequencing
Barcode + constant region (32 nt)
Flow cell binding
Index p5
Index/read primer binding site
20nt p5
Sequencing region
20nt p7
Index/read primer binding site
Index p7
Flow cell binding
p5 primer (90 nt)
p7 primer (90 nt)

Figure S11
